# GFAP Degradation in TBI: Linking Novel Modified Products to Astrocyte Pathology and Patient Outcome

**DOI:** 10.1101/2025.08.01.668181

**Authors:** Ina-Beate Wanner, Julia Halford, Jonathan Lopez, Sean Shen, Yu Chen, Hayden Zhao, Carolina Salas, Rachel R. Ogorzalek Loo, Steven Robicsek, Benjamin M. Ellingson, Jeffrey Gornbein, Timothy E. Van Meter, Gerry Shaw, Joseph A. Loo, Paul M. Vespa

## Abstract

Glial fibrillary acidic protein (GFAP) is a significant clinical biomarker of traumatic brain injury (TBI), yet understanding the nature, timing, and impact of its degraded and modified products would inform clinical utility. We report novel GFAP breakdown products (BDPs) and post-translational modifications (PTMs) that are unique to TBI including fragment- and patient-specific citrullination signatures that destabilize GFAP filaments. GFAP and its fragments were sequenced by mass spectrometry (MS) from severe TBI patients’ cerebrospinal fluid (CSF) and sera, identifying two distinct TBI-specific jointly generated product sets within the rod-domain covering coil1 (20-26kDa) and coil2 (15-19kDa). Their endings differed from GFAP fragments described in Alzheimer’s and Alexander disease. Label-free quantification showed biofluid co-product abundance differences with coil1-BDPs enriched over coil2-BDPs. Coil imbalance was independently confirmed by immunoblot densitometry in twenty-three TBI patients. Measurements over ten days showed injury day peaks of full-length and end-clipped GFAP fragments, while proteolytic 37/39kDa and small fragments remained elevated. Six-month outcome (Extended Glasgow Outcome Scale) correlated with GFAP fragments, whereas levels of uncleaved and end-clipped GFAP did not. A human astrocyte culture trauma model provided mechanistic insights linking fluid GFAP levels, subcellular localization, and injury-induced astrocytopathy. A new cleavage site between the two coils was identified through selective epitope loss. Coil1-BDPs were fluid-released post-injury, while coil2-BDPs remained intracellular. Their cellular retention was explained by non-filamentous aggregation of citrulline-modified coil2-BDP’s in pathological astrocytes. Trauma-triggered proteolysis involved calpains and caspases in specific astrocyte injury states using protease inhibitors and live-dye-reporter imaging. These novel data link TBI biofluid GFAP fragments with assembly-defective GFAP aggregation in pathological astrocytes. Clinical TBI outcome correlated with GFAP degradation rather than with overall GFAP release. These translational findings indicate that TBI biomarker GFAP undergoes degradation and modification alongside astrocyte pathology, providing a new conceptual framework for investigating biomarker-associated astrocyte proteinopathy potentially linked to post-traumatic neurodegeneration.

## INTRODUCTION

Globally an estimated 61 and 69 million people sustain a traumatic brain injury (TBI) each year with the majority being mild TBIs [25, 61]. U.S. population-based survey data and a recent nation-wide meta-analysis suggest that nearly one in five U.S. adults has a lifetime history of TBI, making it the most common neurological disorder [21, 45]. More than half of mild TBI and 85% of moderate-to-severe TBI patients do not achieve long-term recovery [68]. TBI is a neurodegenerative disease with long term loss of brain structure, function and cognitive ability. Identifying which patients will develop chronic neurological deficits after acute injury is limited but would be useful for effective intervention. The underlying mechanisms of evolving neurodegeneration are not well understood and remain an important public health challenge [78]. Chronic impairment and post-traumatic neurodegeneration involve multiple dementia etiologic pathways after TBI, but the driving pathology and molecular mechanisms remain unclear [22, 83]. This study molecularly defines early biomarker signals and trauma-induced proteolytic changes that may persist as the injury progresses and be potentially relevant to post-traumatic neurodegeneration [14, 62]. This was done through detailed study of distinct TBI associated pathological proteoforms of the intermediate filament glial fibrillary acidic protein (GFAP) and its breakdown products (BDPs).

A pathologic hallmark of neurodegenerative diseases is the accumulation of toxic proteoforms arising from disrupted protein homeostasis, characterized by excessive posttranslational modifications (PTMs) and protein degradation [31, 78, 99]. Trauma increases membrane permeability, excitotoxicity, and sustained intracellular calcium elevation that activates modifying and proteolytic enzymes in a series of events that are well documented in preclinical TBI models [4, 8, 23, 80, 104]. Disrupted proteostasis in TBI and neurodegeneration likely affects many proteins, only a few of which are well-studied, including tau, ß-amyloid and α-synuclein. Protein modification and degradation start early after injury but can persist and thereby could serve as indicators of detecting ongoing neurodegeneration and, possibly, to predict recovery [96].

Two complementary strategies could detect ongoing post-TBI neurodegeneration (1) The link of TBI biomarkers to established neuropathology [74], and (2) determining pathological changes in the biomarkers themselves. Calpain and caspase activation have been monitored using αII-spectrin breakdown products (SBDPs), but their ubiquitous expression limits their clinical utility as brain-specific biomarkers [76, 85]. In contrast, astroglial GFAP is highly brain-enriched and extensively studied as biomarker of TBI. GFAP concentrations in biofluids increase after intracranial injuries, rise fast, peak 6-18 hours postinjury, and then decline [33, 72]. Elevated serum GFAP have been associated with poor outcome and increased mortality [73, 79]. GFAP blood testing is FDA approved within a narrow context of use (COU) on injury day [6, 71]. While well-established as an acute diagnostic biomarker, the longitudinal monitoring and investigation of altered GFAP proteoforms remain open priorities [96].

GFAP breakdown products (BDPs) have been reported in biofluids after TBI, and are detected by commercial GFAP assays, but TBI patient-specific fragment sequences and their degradation kinetics remain poorly defined [33, 63, 108]. The mechanistic link between biofluid GFAP levels and the altered proteoforms in human injured astrocytes has not been defined and may provide an important indicator of pathology [75]. In neuropathology, GFAP upregulation is widely used as a reactive astrogliosis indicator, but trauma-release occurs faster than *de novo* GFAP expression [36, 84]. GFAP modification, misfolding, and degradation are known to have detrimental consequences for astrocytes [43, 64, 90]. Those consequences are critical because astrocyte dysfunction would jeopardize critical cerebral functions including maintaining intact neurovascular coupling, integrity of the blood-brain barrier, metabolite support and neural network function [9]. Astrocyte degeneration and injury states after trauma are beginning to be acknowledged [33, 67, 106]. GFAP proteinopathy is relevant for neurodegeneration, since studies of GFAP mutations in Alexander disease have shown that modification and fragmentation of the protein lead to neurotoxic Rosenthal fibers [24, 35, 60]. Elevated biofluid GFAP levels accompany this inherited astroglial pathology [40]. This study aims to define GFAP degradation and modification in TBI biofluids and in injured human astrocytes, and correlates GFAP fragments to postinjury outcomes, which could improve biomarker interpretation and prognosis.

To bridge these translational gaps, we first sequenced and quantified TBI-induced GFAP-BDPs, and mapped their post-translational modifications (PTMs). We sought to determine whether distinct patterns of destabilizing modifications were present, such as citrullinations and acetylations, and determined their temporal changes after TBI in clinical biofluids. Our principal finding is that elevated temporal CSF profiles of fragments, rather than of total GFAP release were associated with poorer TBI patient outcome. Further, we used a human astrocyte trauma model to determine the mechanistic steps in GFAP cleavage and to show the distinct localization fates of the resulting GFAP-BDPs. Together, these findings advance the molecular understanding of GFAP degradation dynamics as an expanded set of neurotrauma biomarkers that may relate to astrocyte pathology during neurodegeneration.

## MATERIALS AND METHODS

### Aim and study design

The objective of this study was to investigate human GFAP degradation and modification in TBI biofluids and in traumatized astrocytes. GFAP proteolysis and key PTMs were examined in TBI patient biofluids using label-free mass spectrometry (MS). GFAP-BDPs were validated by quantitative immunoblot densitometry in an observational TBI patient cohort. Temporal degradation profiles were described, and their trajectories correlated to clinical outcome. These findings were further investigated using a human astrocyte culture trauma model for mechanistic insight. GFAP release and groups of fragments have already been characterized in this trauma model for severity, elapsed time and correlation with cell wounding and cell death [33]. Accordingly, the current study focused on the distribution fates and abundance of trauma-produced individual GFAP-BDPs. Proteolytic enzymes and astrocyte injury phenotypes carrying defective GFAP aggregates were characterized.

### Setting: Severe TBI patients’ demographic and clinical data and brain tissue specimen procurement

This prospective observational TBI cohort study included 112 cerebrospinal fluid (CSF) samples and 3 serum samples from 26 consented adult severe TBI patients admitted to neurocritical care units at level 1 trauma centers at the University of California Los Angeles (UCLA) and the University of Florida, Gainesville (UF). A commercial control serum sample was included (Precision Med; Supplemental Tables S1, S2). The median patient age was 40±19 years. Twenty-two subjects were male and 4 were female. The median score on the Glasgow Coma Scale (GCS) was 6 ±3.4. Enrollment at UCLA occurred between 2018-2023 and at UF between 2007-2010. Deidentified baseline demographic and clinical parameters are documented (Supplemental Table S1). Recovery scores on the Glasgow Outcome Scale (GOS) were available for all patients except one. All except two patients had outcome interviews by phone at six-month postinjury for extended GOS (GOSE).

Cerebral astrocyte cultures were prepared from de-identified human fetal brain specimen aged 16-19.5 gestational weeks, obtained from pre-shelved tissue at the UCLA Translational Pathology Core Laboratory. Tissues were donated after voluntarily aborted pregnancies following stringent ethical regulations outlined at the end (ethics approval and consent to participate).

### Immunoprecipitation and mass spectrometry sequencing of GFAP from TBI patients

MS experiments were performed on six CSF and serum samples from four TBI patients. GFAP was immunoprecipitated using a rabbit polyclonal anti-GFAP antibody cross-linked to protein G conjugated magnetic beads (Dynabeads). Cross-linking used dimethylpimelimidate-2 (DMP, freshly prepared 20mM DMP in 200mM triethanolamine, pH8.6). Immunoprecipitates were denatured by heat and ice and reduced with 1% ß-mercaptoethanol [13, 17, 65]. Eluted material was separated by preparative SDS polyacrylamide gel electrophoresis (SDS-PAGE) using 10% NuPAGE gradient gels in morpholinoethansulfonic acid buffer under reducing conditions along with a molecular weight standard (Benchmark). Gels were Sypro-Ruby stained and fixed in 10% methanol, 7% acetic acid. GFAP bands were cut out under UV light and placed into tubes for in-gel trypsin digestion. A fraction (∼1/9^th^) of the eluate was blotted and GFAP was detected using mouse monoclonal antibodies (mabs) to GFAP (mab1A64-244, Abbott Diagnostics or mab 3E10, EnCor Biotechnology Inc., Supplemental Table S3) to provide reference signals for gel band excision. Bands at 50 and 25kDa were omitted to avoid immunoglobulins (Supplemental Fig.S1). Gel bands were fully de-stained, dried, and then reduced with 10mM tris(2-carboxyethyl) phosphine in 100mM ammonium bicarbonate buffer at 56°C for 30min, followed by alkylation with 100 mM iodoacetamide in the same buffer at 37°C for 15min. Gel bands were equilibrated with acetonitrile (ACN) and dried. Proteins were digested using 0.01mg/ml sequencing grade trypsin in 100mM ammonium bicarbonate buffer at 37°C for 12h. Peptides were extracted from the gel bands by three rinses of 50% ACN/0.1% formic acid solution at 37°C for 15min. and the combined extracts were dried in a vacuum concentrator. The peptides were measured by liquid chromatography-tandem MS (LC-MS/MS) using an Ultimate 3000 nano-LC instrument (1h gradient) that was coupled to a Q-Exactive Plus Orbitrap mass spectrometer. Full spectra (MS1) were collected at a resolution of 70,000 at full width at half maximum (FWHM). The 15 most abundant ions of each scan were subsequently fragmented by higher-energy collisional dissociation (HCD; MS2) with a resolution of 17,500 at FWHM and a dynamic exclusion of 45s. Defined amounts of recombinant human GFAP were digested serving as technical controls (Supplemental Fig.S2). MS data were searched against the human GFAP protein sequence via Proteome Discoverer (PD) version 1.4. SEQUEST-engine Uniprot (https://www.uniprot.org/uniprotkb/P14136/entry). Confirmed peptides were unique to GFAP (not shared by e.g. vimentin or neurofilaments). ‘High’ confidence peptides (green on maps) were defined as peptides with at least two matches that passed a ≤1% false discovery rate (FDR) threshold based on cross-correlation (X-corr) scores, for peptide matches contingent on ion precursor charge states. There were a few ‘medium’ confident peptides (yellow on maps in Fig.1A), defined with at least two peptides that passed a ≤5% FDR. PTMs were identified from TBI patients’ GFAP fragments and included acetylations and citrullinations. Citrullinations were validated by the unique signatures of an immonium ion at mass-to-charge ratio (m/z) of 130.0975Da and/or a neutral loss of isocyanic acid at 43.0058Da. Lysine acetylations were validated by the presence of a cyclized immonium ion at m/z 126Da. Additional validation was provided by MS2 products of b- and y-ions documenting mass increments consistent with these modifications (Supplemental Fig.S5). Citrullinations were further validated by prior reported citrullination sites after treatment with peptidylarginine deiminase (PAD) enzyme (see Tables S4) [42].

**Fig. 1.**
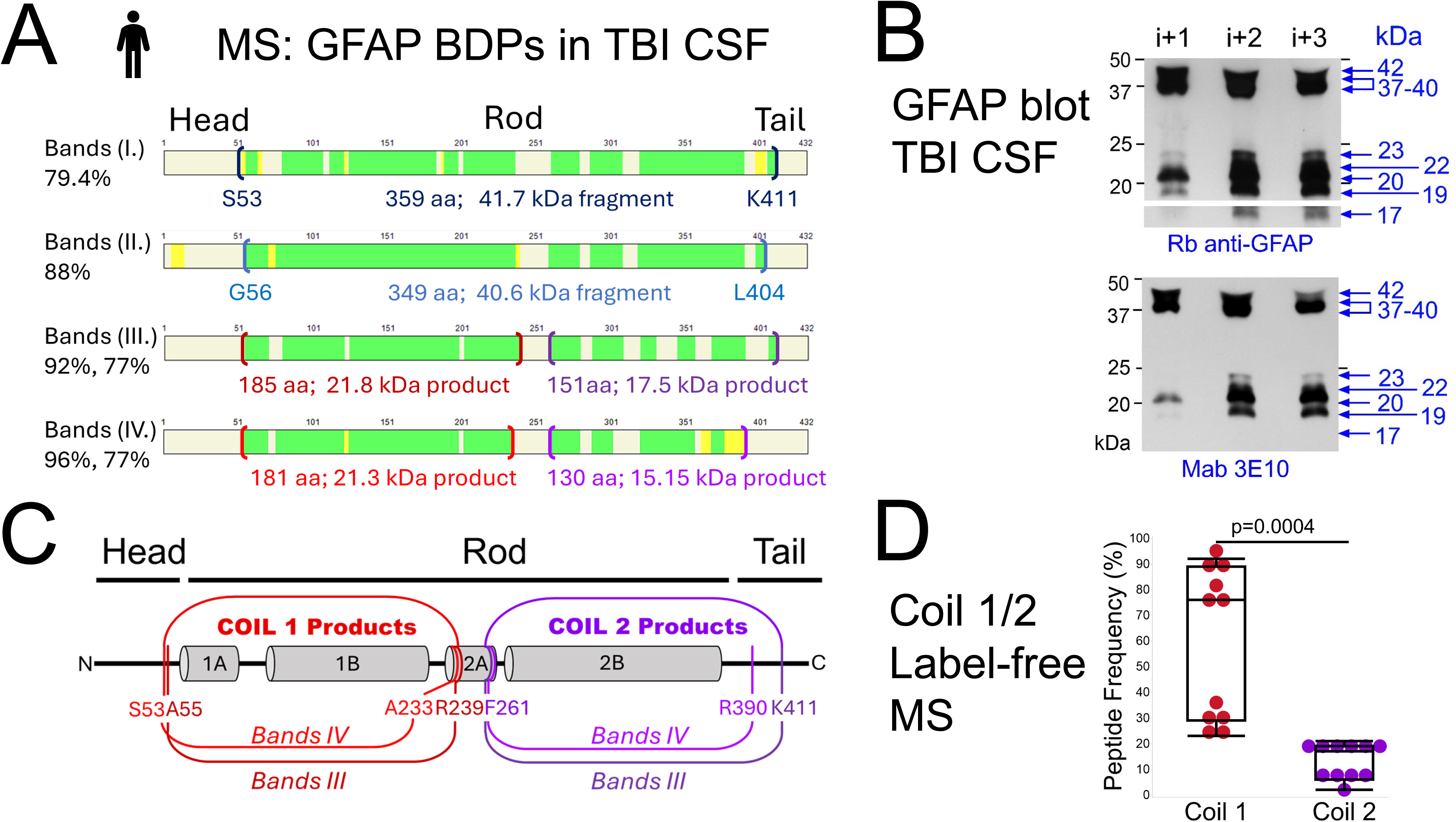
Variable endings and abundance of small GFAP breakdown products in TBI patients’ CSF **A)** Mass spectrometry (MS) sequences of GFAP breakdown products (BDPs) with peptide maps and percent coverage of immunoprecipitated GFAP from four excised gel slices (Bands I.-IV) from six TBI CSF samples (Supplemental Fig.S1). High confidence peptides are shown in green (FDR≤1%), and repeatedly identified medium confidence peptides in yellow (FDR≤%). **Bands I.** contained a 359 amino acids (aa) fragment with calculated molecular weight of 41.7kDa and new serine N-terminus (S53) and lysine C-terminus (K411). **Bands II.** spanned 349aa from glycine (G56) to leucine (L404), encompassing 37/38kDa fragments that varied between TBI samples. **Bands III.** contained two GFAP-BDPs, a coil1 product (185aa, 92% coverage, estimated molecular weight 21.8kDa), and a coil2 product (151aa, 77% coverage, 17.5kDa). **Bands IV.** contained slightly smaller coil1 product (181aa, 96% coverage, 21-21.6kDa ±PTMs) and slightly smaller coil2 product (130aa, 77% coverage, 15-15.3kDa). **B)** GFAP immunoblots at one and three postinjury days (PIDs) confirm the MS fragments. TBI CSF samples are from a 84-year-old male TBI patient with subarachnoid and intraventricular hemorrhage and intraparenchymal hematomas after a fall. Probed with a polyclonal (top) and a monoclonal (clone 3E10, bottom) antibody, observed GFAP bands were: **Bands I.**, 40-42kDa, **Bands II.**, 36-39kDa, **Bands III.**, 20.4-23kDa, and **Bands IV.**, 17-19kDa, the latter detected only by the polyclonal antibody. **C)** Predicted endings of new GFAP products after TBI. **Coil1 products** had either an alanine or serine N-terminus (A55, S53) and a C-terminus ending in alanine (A55) or arginine (R239). **Coil2 products** had a consistent phenylalanine N-terminus (F261) and ended with either lysine (K411) or arginine (R390). They were sequenced from two excised gel slices (Bands III and IV) of different patient samples. **D)** Label-free MS quantification documents peptide detection frequencies within coil1 (59± 29%) and in coil2 (13±7%) products across eleven MS runs of **Bands III.** and **IV.** The difference in peptide detection frequency between the two coils was significant, in a mixed-effect model testing with patient ID as random effect, followed by a two-tailed T-test with unequal variances (p=0.0004; validated by ion peak heights, Suppl. Fig.S3).

### Partial chemical cleavage of GFAP for monoclonal antibody epitope mapping

The binding region of mab anti-GFAP clone 5C10 was determined initially by binding to the peptide covering amino acids (aa) 233-261 of the human GFAP coil2A sequence using dot blots (AALKEIRTQYEAMASSNMHEAEEWYRSKF). The specific epitope was subsequently identified using partial chemical cleavage at tryptophan (W), of which there is only one in GFAP, at position W256. Recombinant GFAP (33µg) was treated in excess with 1% BNPS-Skatole (100µg; 1mg/ml glacial acetic acid; 3-bromo-3-methyl-2-(o-nitrophenylsulfenyl)indolenine, Sigma, Supplemental Fig.S9), an oxidant and brominating agent that selectively cleaves proteins at tryptophan residues [34, 92]. An equal volume of distilled water precipitated BNPS-Skatole, which was subsequently removed by centrifugation (5min at 12,000rpm). The supernatant was lyophilized and GFAP and its tryptophan-cleaved BDPs were separated by SDS-PAGE. After blotting, fragments were detected by incubation of mab 5C10 to identify the 5C10 epitope. In parallel, coil1 fragment was identified by mab 3E10, and a pan-intermediate filament antibody, IFA identified the coil2 harboring fragment (hybridoma supernatant, epitope coil2B aa 358-377: ALDIEIATYRKLLEGEENRI) [53, 77].

### Western blotting using scaled densitometry and GFAP calibration

SDS-PAGE was conducted as previously described using 7.5% acrylamide gels for αII-spectrin and 12% gels for the majority of GFAP data, and 15% gels for small GFAP-BDP size validation (Fig.1B) [33]. Protease inhibitors (cocktail Complete Mini, Roche 1:50) and EDTA (10mM) were added to TBI patient CSF. Samples were equilibrated in Laemmli sample buffer and boiled for 5min. Samples were then placed on ice and ß-mercaptoethanol was added to 1%. A volume of 34 µl per lane was loaded. The same procedure was followed for astrocyte lysates and concentrated culture media. Samples and molecular weight standards (Kaleidoscope, Biorad) were collected upon initial run for 30min at 100V, followed by separation for 1h at 120V until the bromophenol blue front reached the bottom. Proteins were transferred cold at 120V for 3h onto nitrocellulose and stained with Ponceau S. Membranes were blocked in 10% non-fat milk in Tris-buffered saline with Tween (TBST) at room temperature (RT). Incubation with primary antibody was done in 5% BSA in TBST overnight at 4°C. Secondary antibodies conjugated with peroxidase (Pierce) were incubated in 5% BSA in TBST for 1h at RT. Enhanced Chemiluminescence (ECL) substrate was applied to washed membranes for 5min at RT (Pico and Femto ECL, Pierce). Multiple exposures were used either using Kodak films, or with an iBright imager (ThermoFisher Scientific). Calibrants were recombinant human full-length GFAP and a 20kDa His-tagged GFAP-BDP (aa 72-217, EnCor Biotechnology Inc.).

### Calibration, densitometry and precision

Calibrants were diluted with 0.5% BSA in TBS at defined concentrations (0.08-2.6ng). Signals were measured across multiple exposure times (calibration curve, see Supplemental Fig.S6). Densitometry was performed using eight exposures of each blot (2s-20min) after background correction. Sub-saturated band measurements were standardized to 1min by multiplying shorter exposures and dividing longer exposures using scaling factors between exposures that were empirically determined from the same unsaturated on two exposures (Supplemental Table S5). Band sizes were computed from the migration distances of each molecular weight marker from the gel-front using curve-fit equations. Reproducibility is reported with the coefficients of variation (CV) between experiments’ technical replicates for expressing precision of immunoblot densitometry (CV=SD/mean x100, Supplemental Table S6B).

### Antibodies used on TBI CSF and astrocyte samples

All antibodies are tabulated with known epitopes, sources, dilutions, and Research Resource Identifiers (RRIDs, Supplemental Table S3). A rabbit polyclonal GFAP antibody (DAKO) was used for immunoprecipitation and immunoblotting. For densitometry the antibody was applied initially at a high dilution (1:20,000-150,000) to detect full-size GFAP and large GFAP-BDPs, followed by a lower dilution (1:5,000-7,000) to detect small GFAP-BDPs. Signals were adjusted in subsequent scaling procedure (see above). Mab 3E10 was generated to a recombinant construct containing aa 72-217 of human GFAP, comprising coil1A and 1B. The epitope for 3E10 was localized to aa 196-210 (EIRFLRKIHEEEVRE), via peptide competition testing with a set of 20 nested aa peptides, each overlapping by 5aa. Additional mabs were 5C10 (EnCor Biotechnology Inc.), 1A64-244 (Abbott Diagnostics), a mixture of three clones (4A11, 1B4, 2E1, BD Pharmingen), and mab GA5 (Sigma [16, 18]). GFAP-specific citrullination was detected using mouse clone CTGF-1221 that was kindly provided by Akihito Ishigami (Professor, Tokyo Metropolitan Institute for Geriatrics and Gerontology). CTGF-1221 detected citrullinated GFAP (cit-R270-GFAP) at aa sites R270 and R416 [39]. An anti-peptidyl-citrulline antibody clone F95 (EMD Millipore) validated citrullinations. Several additional rabbit monoclonal citrullination site-specific GFAP antibodies were characterized, including cit-GFAP-270 with epitope at GFAP aa 270 in coil2 (BRAINBox Solutions, Inc.). A mab against αII-spectrin was used at 1:3,000-1:5,000 dilution (Enzo Life Sciences).

### Human cerebral astrocyte trauma model

#### Human culture trauma model

In vitro differentiated fetal cerebral astrocytes were prepared as previously described [95]. Briefly, cortical tissue was mechanically dissociated, filtered and astrocytes were enriched by Percoll gradient centrifugation. Cells were expanded in DMEM/F12 with 10% fetal bovine serum. Residual oligodendrocytes and progenitor cells were removed by shaking. Astrocytes were seeded on collagen-I-coated deformable silastic membranes at 138,000-180,000 cells/9.62cm^2^. Astrocytes were differentiated for three weeks by stepwise serum withdrawal in a 1:1 mixture of DMEM/F12: Brainphysiology medium (StemCell Technology, supplemented with SM1 and N2A). Astrocytes were traumatized by 50ms pressure pulses (4.9-5.3psi) that abruptly stretched the culture bottom. This inflicted cell shearing that compromised membrane integrity and led to delayed cell death [33].

#### Live microscopy imaging

Live imaging was done 5h after stretch injury on a Leica DMI8 microscope in a gas and temperature-controlled chamber (Okolab). Cells were bathed for 5min in 60µM propidium iodide (PI, Sigma) to label damaged membranes prior imaging. Washed cultures were live-imaged in the presence of two permeable fluorescent protease substrate reporters for calpains (30µM CMAC tBoc Leucyl-methionine, ThermoFisher), and for caspases (5µM DEVD-NucView488, Biotium).

#### Astrocyte protein sample preparation for gel electrophoresis

Whole cell lysate (WCL) of adherent astrocytes was prepared by adding 150µl lysis buffer/9.62 cm^2^ wells (150mM NaCl, 20mM Tris, 5mM EDTA, 10mM NaPyrophosphate, 30mM phosphatase substrate, 5mM dithiothreitol, DTT, Calbiochem, 1% NP40 and 1 tablet/10ml protease inhibitors, Mini Protease Inhibitor Cocktail, Roche). Lysis buffer was incubated for 5min at 4°C, lysates were collected by scraping and transferred into low binding tubes. Proteins were homogenized by 3x5s sonification bursts without foaming (Branson Digital Sonifier, 90% or constant amplitude duty cycle, output level 5). Culture medium conditioned by astrocytes (CM) was collected in absence of serum or protein supplements. Added were 5mM DTT and 1tablet protease inhibitors/10ml CM. Lifted cells and debris were pelleted by initial centrifugation (5,000g, 5min, 4°C). Pellets were then lysed for 5-10min in 0.5% lysis buffer followed by 3x5s sonication. Remaining insoluble fractions were removed by centrifugation (10min at 16,000g 4°C). Media were concentrated by centrifugation until reaching ∼1/10^th^-20^th^ of the original volume (3kDa cutoff Vivaspin filtration columns, 4-8h at 5000g, 4°C). CM yielded a protein concentration of 1-1.5 µg/ml. All samples were immediately frozen in aliquots at -80°C. Immunoblotting and densitometry were performed as described for TBI CSF samples.

#### Protease inhibition in the human trauma culture model

A non-toxic, cell-permeable peptide aldehyde pan-calpain inhibitor, calpeptin was used at 13µM (Z-Leu-nLeu-H, Millipore/Sigma) [57]. A cell-permeable, irreversible pan caspase inhibitor, ZVAD-FMK was used at 11µM (benzyloxycarbonyl-Val-Ala-Asp(OMe)-fluoromethylketone, Enzo) [27]. ZVAD-FMK is reported to also inhibit calpain [50]. Inhibitors were applied to cultures 10min after stretching, omitting gentamicin from the medium. Drugs were incubated with cells for 48h and were replenished every 8-10h. Other tested calpain inhibitors included leupeptin (20µM), SJA6017 (10µM, Calpain Inhibitor VI, Calbiochem), PD1500606 (100µM, Calbiochem), MDL28170 (4µM, Z-Val-Phe-al, Calbiochem). These were not further pursued because they had limited effect or were toxic by increasing cell permeability or cell death.

### Statistical analysis approaches

GFAP band measurements were initially assessed for normality using the Shapiro Wilk test for Gaussian distribution fit. Measurements deviated from normal distribution (significantly lowered W-values) and were therefore log-transformed establishing normal distribution. Zeroes were replaced with half of the lower limit of detection (0.7pg or a concentration of 27pg/ml; Supplemental Fig.S7).

The primary statistical method was comparison of biomarker means over time and between subgroups using a linear mixed-effects model (LMM) for repeated patient measurements. Elapsed time after trauma was a fixed effect, with patient ID modeled as a random effect. Variance components were estimated using restricted maximum likelihood (REML), a conventional and less biased method of estimating random variances. Means were adjusted for age. Body temperature did not have a significant influence on biomarker levels and was therefore not retained in the final model. Sex and race were recorded, but sparse strata in this small cohort precluded stable adjustments, these factors may therefore contribute to unexplained residual variance. For graphing, temporal profiles were estimated by knotted splines (Fig.3C-H).

**Fig. 2.**
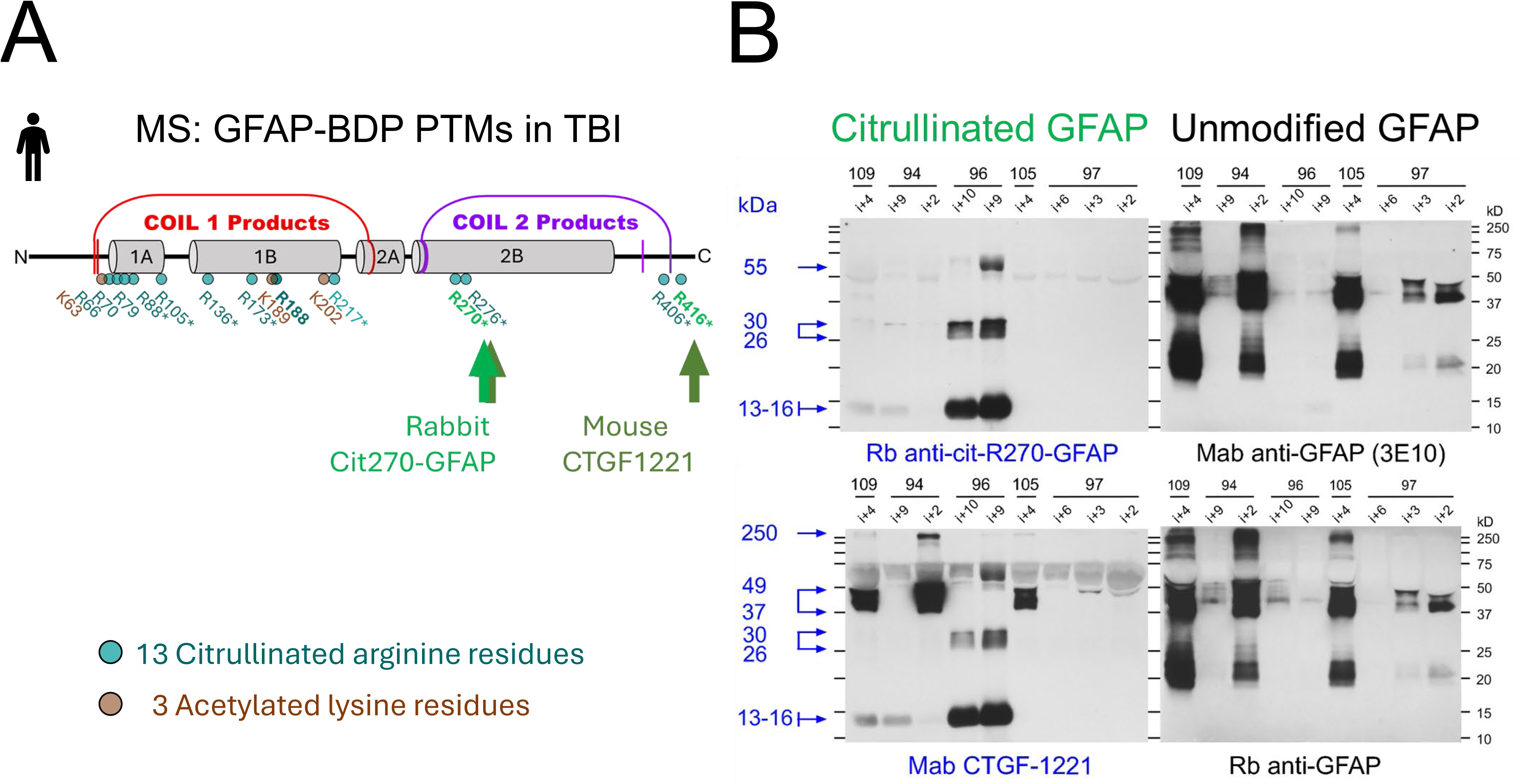
Citrullination and acetylation PTMs are new signatures of TBI patients’ GFAP breakdown products **A)** MS mapping of PTMs in GFAP coil1 and coil2 products from six TBI CSF samples. There were three acetylations (brown) and 13 citrullinations (green), including a novel citrullination site (R188). All PTMs were validated by signature ions and expected mass shifts in MS2 spectra (Supplemental Table S4, Supplemental Fig.S5). A subset of citrullinations (*) is further confirmed by prior identified cell-free peptidylarginine deiminase (PAD) digests [53]. Arrows indicate neo-epitopes at R270 and R416 detected using citrullinated-GFAP-specific antibodies. **B)** *Left*: TBI patient-specific CSF immunoblots at indicated postinjury days (PIDs i+2 to i+10) show citrullinated GFAP bands with strong BDP signals of both antibodies and additional higher molecular weight signals with CTGF-1221[50]. *Right*: Unmodified GFAP in same TBI CSF samples is detected by cloil1 mab3E10 (*top*) or a rabbit polyclonal GFAP antiserum (*bottom*) that document proportional differences in unmodified GFAP signals versus modified ones.

**Fig. 3.**
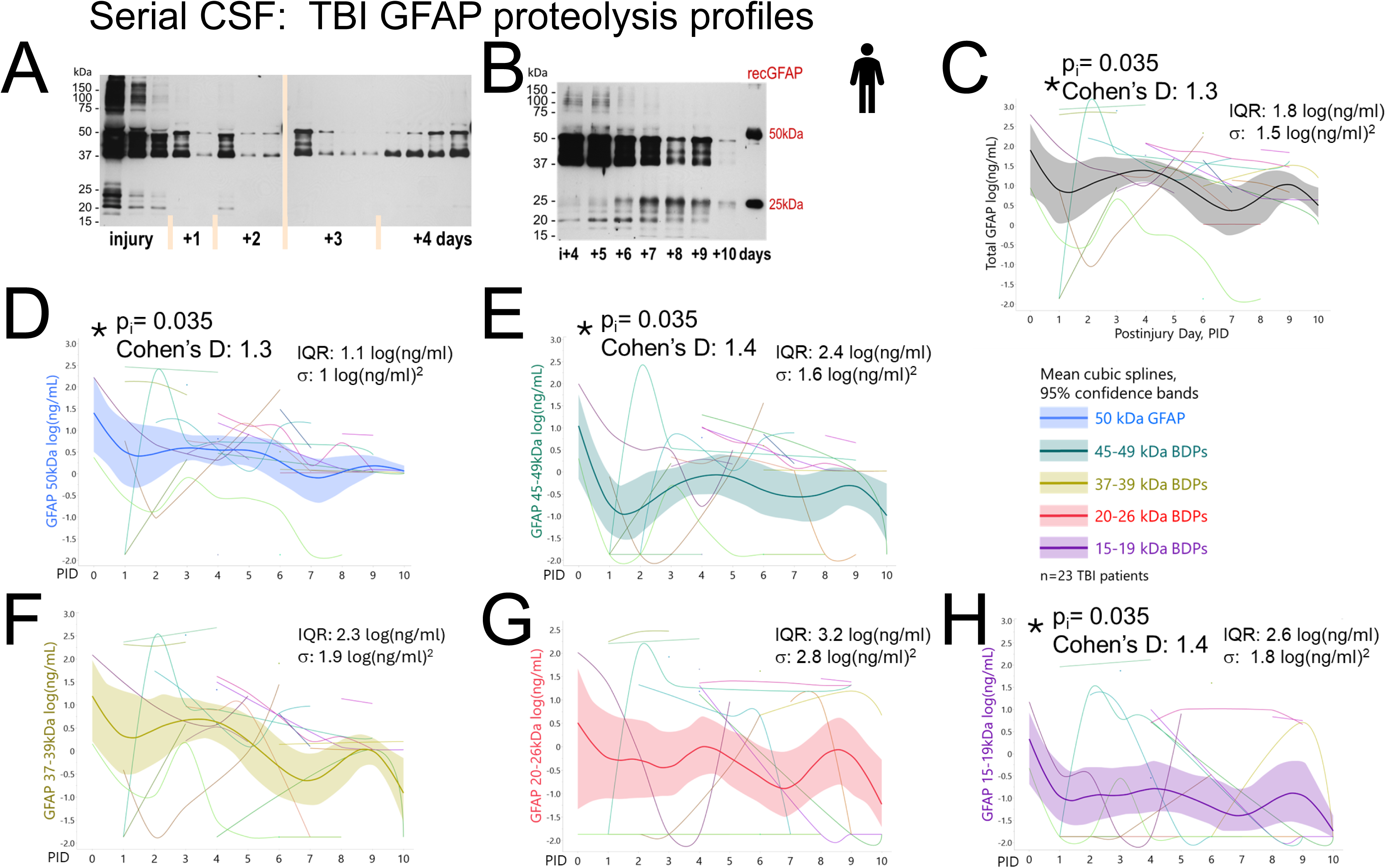
Longitudinal GFAP proteolysis in TBI patients’ CSF with shared peaks and individual profiles **A)** GFAP immunoblot of serial CSF samples over four PIDs from a 40-year-old male TBI patient (admission GCS 3, GOSE 8) shows highest GFAP and fragment levels at 6 hours with additional peaks on PIDs 2-4. **B)** Serial CSF GFAP data over PIDs 4-10 from a 48-year-old male TBI patient (admission GCS 10, GOSE 6) shows declining 50-37kDa bands with later variably increasing small 15-26kDa BDPs. Calibrants are human recombinant full-length GFAP (50kDa) and his-GFAP-BDP (25kDa). **C-H:** Ten-day profiles in 23 TBI patients (102 samples) with resampled mean cubic splines and 95% CI. Graphed are log-transformed calibrated concentrations of subsaturated band densities (calibration curves in Supplemental Fig.S6; densitometry see Supplemental Table S5). **C)** “Total” GFAP signals show a significant injury day peak with 1.3 SD units above adjusted mean across time using a LMM, controlling for random donor and age variance. **D)** Uncleaved GFAP (50kDa) displays a significant injury day peak with decline over 10 PIDs. **E)** End-clipped GFAP-BDPs (45-49kDa) show significant injury day peaks (effect sizes see Table 1A). **F)** Mean 37-39kDa GFAP fragment levels remained above time-adjusted mean during first three PIDs and were below it after PID5. **G)** 20-26kDa GFAP products showed the greatest dispersion with highest interquartile range (IQR) and variance (σ) yet displayed mean secondary elevations on later days. **H)** 15-19kDa GFAP products display an injury day peak and lowest profile, due to a higher fraction of absent bands, yet when present, these bands had strong signals.

**Table 1:**
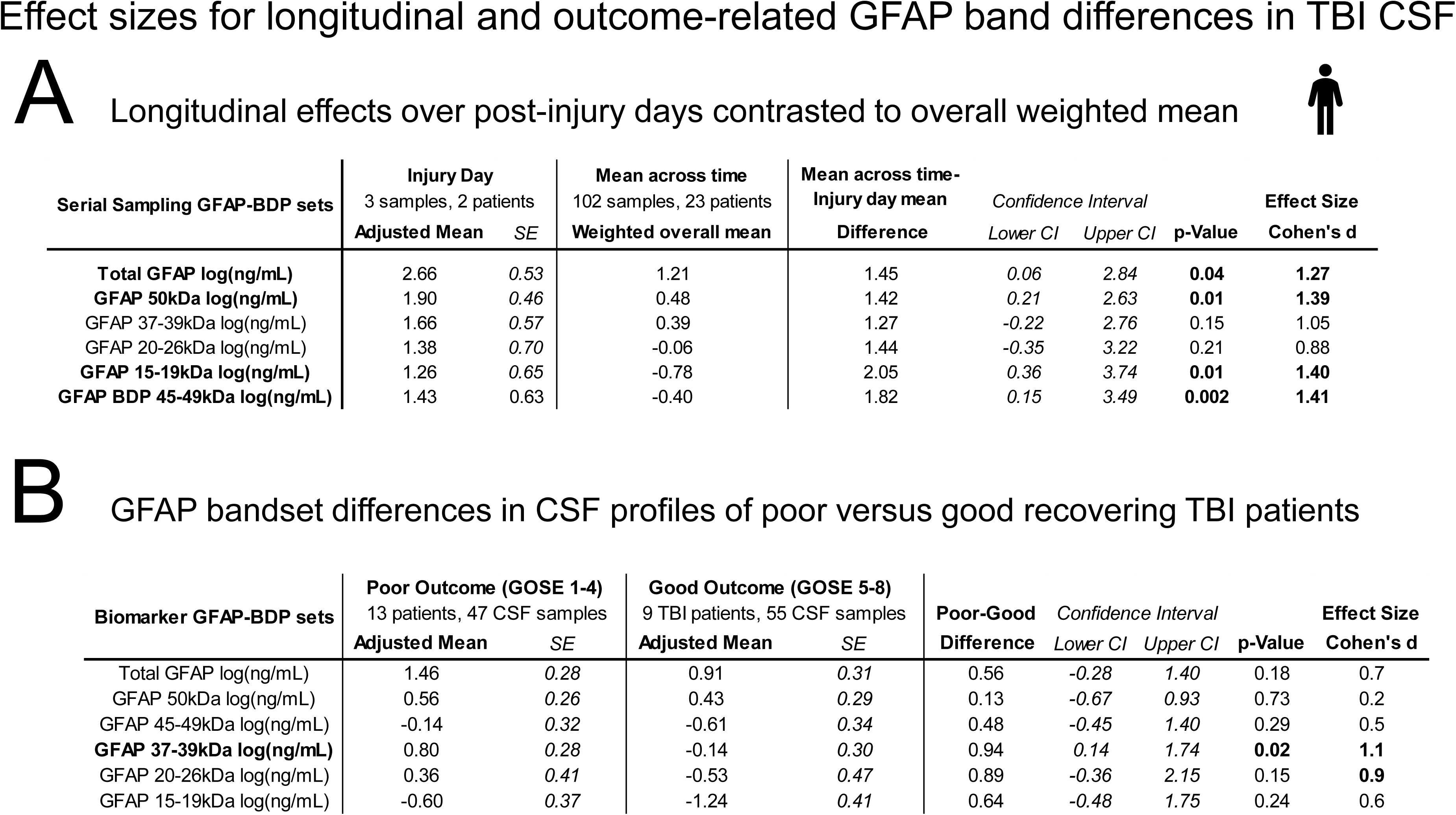
Effect size for longitudinal and outcome-related GFAP band differences in TBI CSF **C)** Longitudinal effects over PIDs in contrast to overall weighted mean. This table shows injury day GFAP signal contrasted to overall mean signals from serial samples weighted by available samples [37]. Means of each PID were adjusted for age and each day’s difference from the overall average across time was computed with its upper and lower 95% confidence intervals (CI). Cohen’s d effect size values were computed dividing this difference by the squareroot of the entire random variance component of this REML linear mixed model conservatively accounting for within and between subject variance. Effect sizes for each GFAP bandset are given with t-statistic-derived p-value. Only injury day GFAP signals had marked differences, other PIDs are not shown. **D)** GFAP bandset differences in CSF profiles of TBI patients with poor versus good six month recovery. Total, uncleaved and fragment GFAP bands are compared between 13 TBI patients with poor outcome (GOSE 1-4, 47 CSF samples) and 9 patients with good outcome (GOSE 5-8, 55 CSF samples). Age-and time-adjusted means of each group are presented with their SE from a REML linear mixed model including subject ID as random effect. For each GFAP bandset, Cohen’s d was computed as the adjusted mean difference between the two groups divided by the residual variance, reflecting within subject variation. 95% CI for these differences and corresponding p-value for these effect sizes are provided.

| Serial Sampling GFAP-BDP sets | Injury Day<br>3 samples, 2 patients |  | Mean across time<br>102 samples, 23 patients | Mean across time-Injury day mean | Confidence Interval |  | p-Value | Effect Size |
| --- | --- | --- | --- | --- | --- | --- | --- | --- |
|  | Adjusted Mean | SE | Weighted overall mean | Difference | Lower CI | Upper CI |  | Cohen's d |
| Total GFAP log(ng/mL) | 2.66 | 0.53 | 1.21 | 1.45 | 0.06 | 2.84 | 0.04 | 1.27 |
| GFAP 50kDa log(ng/mL) | 1.90 | 0.46 | 0.48 | 1.42 | 0.21 | 2.63 | 0.01 | 1.39 |
| GFAP 37-39kDa log(ng/mL) | 1.66 | 0.57 | 0.39 | 1.27 | -0.22 | 2.76 | 0.15 | 1.05 |
| GFAP 20-26kDa log(ng/mL) | 1.38 | 0.70 | -0.06 | 1.44 | -0.35 | 3.22 | 0.21 | 0.88 |
| GFAP 15-19kDa log(ng/mL) | 1.26 | 0.65 | -0.78 | 2.05 | 0.36 | 3.74 | 0.01 | 1.40 |
| GFAP BDP 45-49kDa log(ng/mL) | 1.43 | 0.63 | -0.40 | 1.82 | 0.15 | 3.49 | 0.002 | 1.41 |

Table 1
| Biomarker GFAP-BDP sets | Poor Outcome (GOSE 1-4)<br>13 patients, 47 CSF samples |  | Good Outcome (GOSE 5-8)<br>9 TBI patients, 55 CSF samples |  | Poor-Good | Confidence Interval |  | p-Value | Effect Size |
| --- | --- | --- | --- | --- | --- | --- | --- | --- | --- |
|  | Adjusted Mean | SE | Adjusted Mean | SE | Difference | Lower CI | Upper CI |  | Cohen's d |
| Total GFAP log(ng/mL) | 1.46 | 0.28 | 0.91 | 0.31 | 0.56 | -0.28 | 1.40 | 0.18 | 0.7 |
| GFAP 50kDa log(ng/mL) | 0.56 | 0.26 | 0.43 | 0.29 | 0.13 | -0.67 | 0.93 | 0.73 | 0.2 |
| GFAP 45-49kDa log(ng/mL) | -0.14 | 0.32 | -0.61 | 0.34 | 0.48 | -0.45 | 1.40 | 0.29 | 0.5 |
| GFAP 37-39kDa log(ng/mL) | 0.80 | 0.28 | -0.14 | 0.30 | 0.94 | 0.14 | 1.74 | 0.02 | 1.1 |
| GFAP 20-26kDa log(ng/mL) | 0.36 | 0.41 | -0.53 | 0.47 | 0.89 | -0.36 | 2.15 | 0.15 | 0.9 |
| GFAP 15-19kDa log(ng/mL) | -0.60 | 0.37 | -1.24 | 0.41 | 0.64 | -0.48 | 1.75 | 0.24 | 0.6 |

#### Age covariate effect

The effect of age was summarized using partial omega squared (ω_p_^2^) the proportion of variance in GFAP levels explained by age. This method is commonly used because it provides a less biased estimate of the population effect size than eta squared, suited for smaller samples with unequal group sizes [54]. ω_p_^2^ was computed from the F-statistic of the age term according to equation (1, SAS/Jmp):

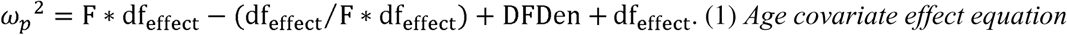

df: degrees of freedom; DFDen, denominator term of degrees of freedom (see Supplemental Table S7).

***Temporal profile effect size:*** Longitudinal mean profiles were graphed by resampled spline fit lines from daily CSF draws with 95% confidence intervals (CI, Fig.3C-H). LMM-based effect sizes were Cohen’s d mean differences of any postinjury day age-adjusted means from the overall weighted mean in standard deviational (SD) units using the squareroot of the total residual variance as pooled SD according to equation (2): [37, 55].

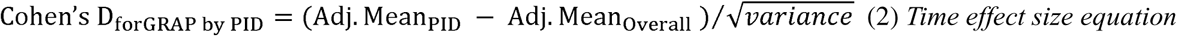

Adj.Mean PID: Age-adjusted mean on an individual postinjury day Adj. Mean overall: Weighted overall age-adjusted mean [37].

Temporal effect sizes are reported with upper and lower 95% confidence intervals (CI) of the mean difference on injury day versus overall mean with p-value of its difference (Table1).

#### Bivariate GFAP and fragment correlations

To assess GFAP substrate-product and product-product relationships, all log-transformed GFAP band signals were analyzed using pairwise Pearson correlations and reported as unadjusted correlation coefficients between continuous variables (Fig.4A). Biplots used orthogonal (Deming) fit, assuming equal variance between GFAP bands and are reported with coefficient of determination (R^2^) and Benjamini-Hochberg FDR-adjusted significance expressed as Logworth, -log_10_(p-values) (Fig. 4B, C). Outliers were flagged using a robust rule, defined as percentage of datapoints exceeding three times the residual SD. Correlation outliers that could influence associations between variables were inspected using the Mahalanobis distance, which identifies potential bivariate outliers by an upper control limit determined from the sample size and the number of variables, applying robust weighted estimation [32].

**Fig. 4.**
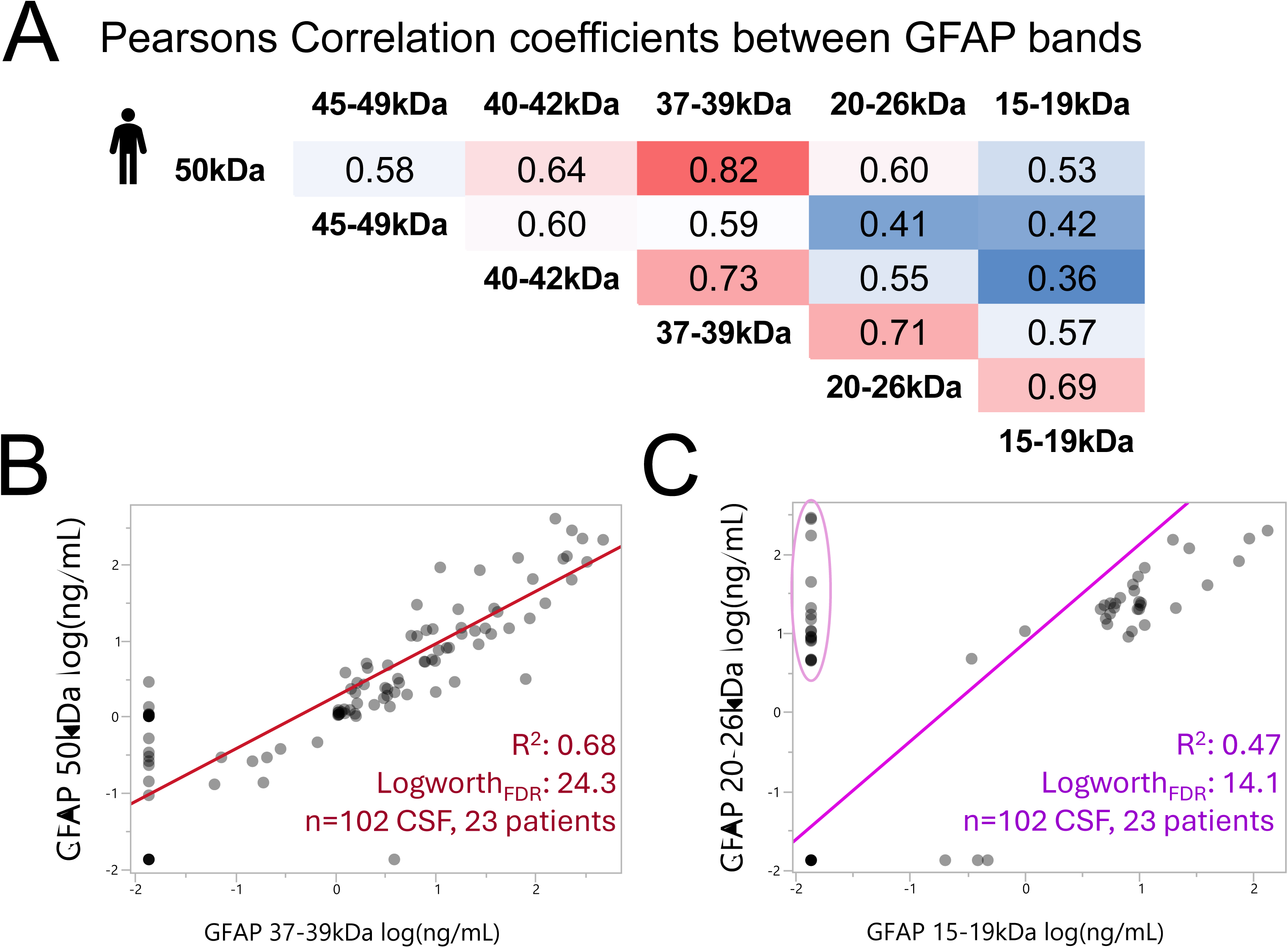
Correlation between GFAP uncleaved, fragments, and co-products in TBI CSF **A)** Pearson’s correlation matrix lists coefficients between uncleaved GFAP and fragment sets measured in 102 CSF samples from 23 TBI patients. **B)** Bivariate plot of full length (50kDa) GFAP substrate and calpain-generated fragments (37-39kDa) using an orthogonal fit (R^2^ with equal error weights) and FDR-corrected Logworth (log_10_p-value). **C)** Bivariate plot of GFAP coproducts shows weaker correlation, due to higher frequency of undetected 15-19kDa versus 20-26kDa bands (circle, 28% 15 undetected cases).

#### Outcome differences among GFAP species

The purpose was to determine whether GFAP profiles over time differed between patients with good versus those with poor six months outcome to qualify among GFAP and its fragment bands. Group differences were assessed by comparing adjusted mean temporal trajectories between the two groups using LMM model-derived and reporting the magnitude of the difference with effect size using Cohen’s d. Here Cohen’s d was calculated from the model-derived mean difference between the two outcome groups for each GFAP bandset expressed in units of the pooled residual SD (Equation 3). Effect sizes are reported with lower and upper 95% CI and corresponding posthoc t-statistic-derived p-values for each bandset (Table1). GFAP trajectories for good (GOSE 5-8), and poor (GOSE 1-4) outcomes were graphed using common cubic knotted spline curves with same curvature parameters (λ= 0.28) and 95% CI (Fig. 5).

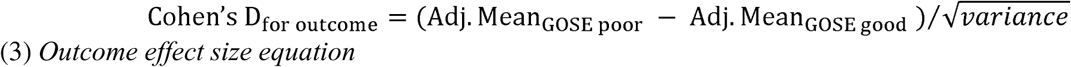

**Fig. 5.**
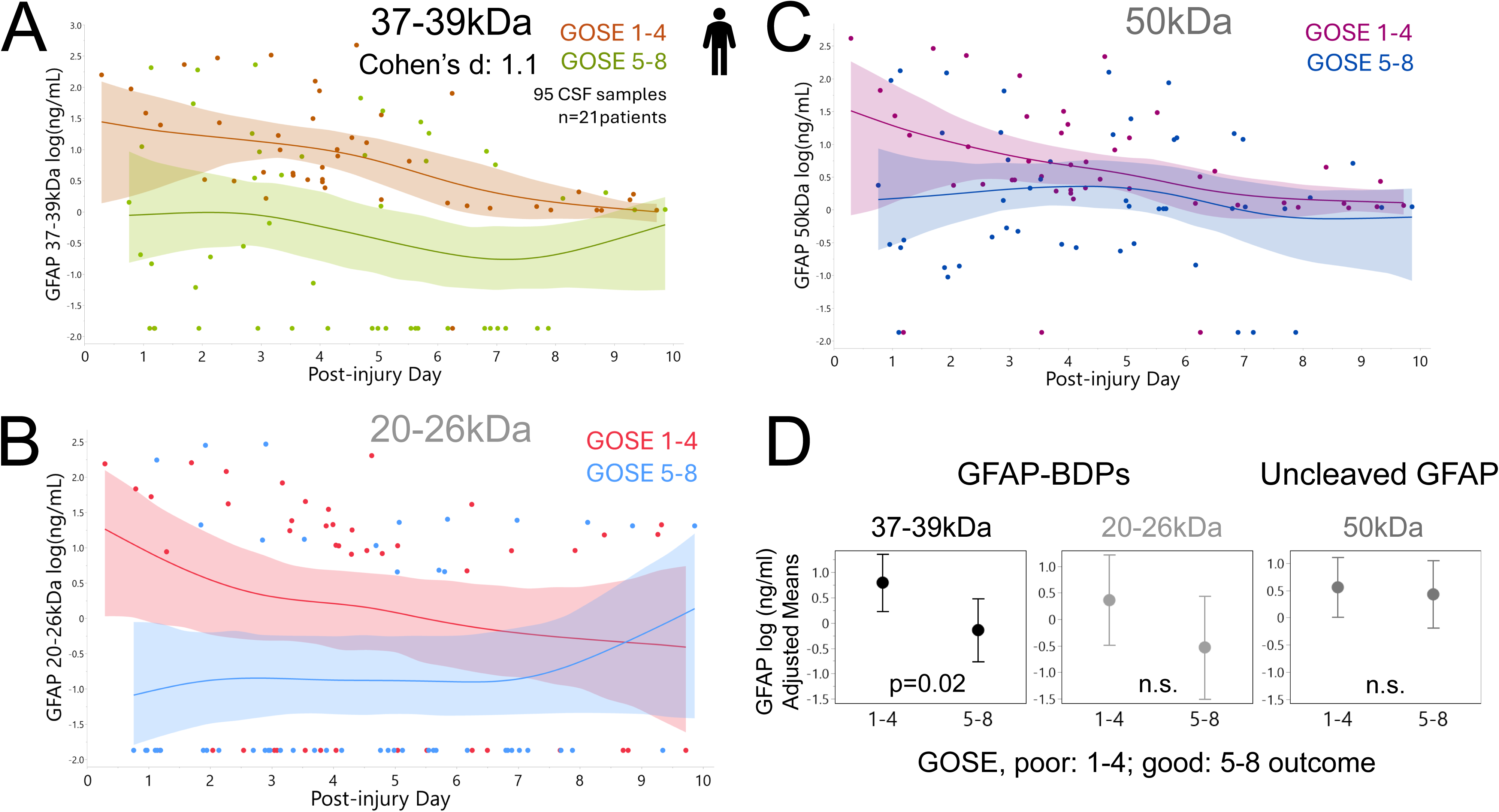
Longitudinal GFAP proteolysis correlates with TBI patient outcome, not that of full-length GFAP. Compared are trajectories of GFAP bandsets over 10 PIDs in TBI patients with good recovery (GOSE 5-8) versus poor outcome (GOSE 1-4). Graphs show mean cubic splines and 95% CI (21 patients; 95 observations, effect sizes see Table 1B). **A)** Trajectories of calpain-derived GFAP fragments (37-39kDa) differed between outcome groups. **B)** Levels of small GFAP-BDPs (20-26kDa) tended to be higher in the poor versus the good outcome group. **C)** Full-length GFAP (50kDa) profiles overlapped across time between groups. **D)** GFAP band LMM-adjusted for elapsed time, age and ID are graphed with their SE. These adjusted means illustrate differences in GFAP bands between TBI patients with good versus poor recovery.

Adj.Mean_GOSE poor_ : Each GFAP bandset CSF concentration means (47 CSF samples 13 TBI patients with GOSE scores 1-4) were adjusted for age and elapsed time (fixed effects) and patient ID (random effect).

Adj.Mean_GOSE good_: Each GFAP bandset CSF concentration means (55 CSF samples, 9 TBI patients with GOSE scores 5-8), with same adjustment structure.

Cohen’s d denominator: Pooled SD: Square root of the residual variance from the model (within subject variability).

#### Label-free MS analysis

The frequency of peptide presence was compared between coil1- and coil2-derived peptides using a LMM with unequal variances, with patient ID as a random effect and GFAP-coil as a fixed effect (Fig.1D). The same structure was used to compare the peak ion abundance of coil1 versus coil2 ions, excluding undetected peptides (Supplemental Fig.S3).

#### Cell culture fluid GFAP 37 and 20kDa band analysis over time post-injury

Densitometry was done as described previously for 4-9 donors (7-12 pooled samples per condition) [33]. GFAP 37kDa and 20kDa band signals were compared at 0.5, 5, 24 and 48h post-stretching with unstretched samples as well as over time (Supplemental Fig.S12). LMM random donor adjusted means with time as fixed effect was performed and differences between stretched and unstretched adjusted mean fluid GFAP levels were reported as effect size using Cohen’s d with residual variance as pooled SD.

## RESULTS

These integrative translational studies were designed to (1) define the GFAP degradome in TBI patients by rigorous MS sequencing and PTM determination with validation via immunoblotting; (2) delineate temporal profiles and relate fragment patterns to clinical outcomes in an exploratory cohort; (3) obtain mechanistic insight on fragment release after cleavage by proteases; and (4) localize pathologically aggregated GFAP-BDP proteoforms within dystrophic astrocytes after stretch injury.

### Novel trauma-specific GFAP breakdown products in TBI patients show fluid imbalance and unique PTMs

#### Consecutive GFAP cleavage events with defined fragment sequences in cerebrospinal fluid of TBI patients

GFAP displays a range of fragments that increase after TBI [23, 33]. TBI-specific GFAP fragments were immunoprecipitated and labeled as ‘Bands I.-IV.’ (Supplemental Fig.S1). Bands were sequenced by MS with observed peptides used to construct band-specific maps (Fig.1A).

Molecular masses of each slice were estimated from each band’s gel migration and observed sequences from combined MS output. Band I contained a 41.7kDa fragment, and Band II sequences covered a region within 40.6kDa and migrated between 36-39kDa as shown by GFAP immunoblots (Fig.1B). Bands III yielded two sequence blocks in the alpha-helical rod domain covering a 21.8kDa coil1-product and partially covering a 17.5kDa coil2B-product. Bands IV included shorter versions of both fragments, a 21.3kDa coil1-product and a 15.2kDa coil2-product (Fig.1A, C). These small GFAP-BDP sets indicate consecutive GFAP proteolysis, with some variability after TBI. Jointly, these sequences confirm previously observed 42, 37, and 18-25kDa GFAP-BDPs and newly identified coil1 and coil2 products [33].

### GFAP breakdown products in TBI differ from Alzheimer’s and Alexander Disease fragments

Coil1 products were identified with a common N-terminus but different C-termini resulting in a longer and shorter fragment (Fig.1B). These C-termini extend beyond a previously reported caspase-6 cleavage site (VELD225) associated with Alexander disease, indicating distinct processing in TBI [16]. Immunoblots using a polyclonal GFAP antibody as well as mab 3E10, recognizing an epitope in coil1, both detected small GFAP-BDPs with molecular masses between ∼19-23kDa (Fig.1B). Coil2-products were also identified with a common N-terminus and different C-termini, producing longer and shorter BDPs (Fig.1C). The polyclonal GFAP antibody detected an additional ∼17kDa band that was not recognized by coil1-specific mab 3E10, providing immunoblot evidence for coil2 fragments (Fig.1B). These coil2-products start at phenylalanine (F261), consistent with an Alzheimer’s disease (AD)-reported caspase-3 cleavage motif (aa258-266, RSKFA↓DLTD). Yet these TBI observed coil2 products were considerably smaller than the ∼30kDa AD-associated fragment [66]. None of the small BDPs contained peptides in the linker region coil2A, while this region was readily detected in full-length recombinant GFAP (Supplement Fig.S2).

This work documents, for the first time, differences in GFAP-BDPs produced following TBI compared to other neurodegenerative diseases. The data underscores that neurological diseases can differ in their proteolytic activities resulting in distinct GFAP-BDP patterns. The findings on trauma-specific GFAP fragments also speak against the generalized GFAP fragmentation observed in cell-free systems through in vitro digests of recombinant GFAP, or after treating cells with toxins [103].

### Imbalance of coil1 and coil2 GFAP breakdown products in TBI patients’ serum and CSF

High-confidence peptide coverage was greater in coil1 than in coil2 products. This imbalance was quantified by how frequently peptides were detected across MS runs, documenting significantly higher coil1 (59±29%) versus coil2 (13±7%) detection frequencies (Fig.1D). Ion peak intensities were also significantly higher in coil1 versus coil2 peptides, further supporting greater coil1 abundance in TBI CSF (Supplement Fig.S3).

In serum, TBI patients showed more intense 37kDa and 20kDa bands than controls (Supplemental Fig.S4A). Immunoprecipitated GFAP from TBI serum samples identified a coil1-product of 97 residues with calculated mass of 18.2kDa (aa51 to R217 Supplement Fig.S4). Of note, coil2-containing peptides were absent in serum, providing additional evidence of an imbalance between the two coils in biofluids. In summary, deep sequencing of GFAP-BDPs in TBI patient biofluids identified proteolytic processing expected to generate stoichiometrically equimolar coproducts, yet coil1-products were overrepresented relative to coil2 products.

### Citrullinated GFAP-BDPs show a TBI patient signature distinct from that of classic GFAP

Given the observed GFAP product imbalance, we suspected fragment PTMs could help further characterize their disparity. Acetylation and citrullination promote selective destabilization and aggregation and are associated with neurodegeneration [44, 75]. Notably, citrullination has a destabilizing effect on intermediate filaments and has been reported in rodent injury models [56, 98]. To our present knowledge, GFAP-PTMs have not been characterized in human TBI patient samples.

### Validated mapping of citrullination and acetylation in GFAP fragments from TBI patients

Mass spectra of small GFAP-BDPs were searched for evidence of acetylation and citrullination as PTM signatures in TBI patients. GFAP modifications could represent GFAP pathology following TBI and may support biomarker assay development. Acetylated and citrullinated peptide fragments were validated through the presence of signature ions and neutral mass losses (Supplemental Table S4, representative spectra see Supplemental Fig.S5). Coil1 products contained three validated acetylated lysine residues and four citrullinated arginine residues, including a potential novel citrullination site (R79). Three citrulline residues were validated by MS2 for Coil2 products. Additionally, five citrullinations detected in these small GFAP-BDPs had been previously reported in GFAP by PAD treatment [42]. Coil1 and Coil2 acetylation and citrullination were robust and repeatedly detected in 6 CSF samples from 4 TBI patients (Fig. 2A).

### TBI patients have different citrullinated versus unmodified GFAP proportions in cerebrospinal fluid

Citrullination was further analyzed using site-specific citrullinated GFAP antibodies. Two cit-GFAP antibodies with neoepitopes in coil2 (Fig.2A) were characterized here in TBI CSF samples. Both citrullinated GFAP antibodies detected patient-specific signals that differed from those detected by antibodies to unmodified GFAP in the same TBI samples (Fig.2B). The Cit-GFAP antibody detected strong signals in smaller bands, including 13-16kDa and 26-30kDa GFAP-BDPs, relative to its detected signals of uncleaved and larger GFAP-BDPs. Antibodies to unmodified GFAP favored full-length GFAP and other larger fragments but did not detect smaller 13-16kDa and 26-30kDa BDPs. Large GFAP-BDPs were detected by CTGF-1221, an antibody with a second epitope (R416) to the more C-terminal region located outside of the coil2 fragments. On the contrary, cit270-GFAP is directed solely to R270 within coil2 products. This epitope difference could explain further downstream citrullinated larger GFAP fragments detected by CTGF1221 (37-49kDa). In addition, larger citrullinated bands may reflect ubiquitinated or multimeric forms [98]. Substantial patient-to-patient variation highlights individual differences in proteolysis and PAD activities after TBI. These preliminary observations invite future studies on citrullinated GFAP-BDPs for molecular profiling of TBI patients.

### Temporal GFAP proteolysis and fragment-specific imbalance in TBI patients

#### Temporal dynamics of GFAP degradation after TBI

MS fragment sequences were independently validated by quantifying MS-matching band-sets of GFAP-BDPs in serial CSF samples over ten postinjury days (PIDs, 23 patients). Scaled immunoblot densitometry, standardized across multiple exposures, was calibrated with known dilutions of recombinant full-length and fragment GFAP proteins (calibration curves see Supplemental Fig.S6). Exposure scaling expanded the dynamic range to over four-five orders of magnitude, with limits of detection and quantification given in Supplemental Table S6A. Inter-experimental replicates of immunoblot band densitometry had an average technical reproducibility of 28% CV (Supplemental Table S6B).

Longitudinal GFAP profiles document substantial variability over ten PIDs expressed by interquartile range (IQR) and variance (σ) (Fig.3). Age accounted for 23-38% of the GFAP variance after adjusting for time and donor, indicating higher GFAP levels in older patients (age effects sizes see Supplemental Table S7, Supplemental Fig.S8). Temporal trends were analyzed by day relative to mean levels across time using age-adjusted effect sizes (Cohen’s d, see Methods, Table 1A). Summated “total” GFAP signals peaked on injury day (p=0.035) with a Cohen’s d of 1.3 SD units above overall weighted mean signals. Strongest GFAP signal contribution came from full-length GFAP (d=1.3) and large fragments (d=1.4) with their peaks on injury day (Fig.3D-E). The 37-39kDa BDPs exceeded their mean during the first four PIDs and fell below it by day 7 onwards (Fig.3F). The 20-26kDa products showed largest variance, yet their mean remained elevated at later PIDs (Fig. 3G). The 15-19kDa products were sporadically detected with highest levels on injury day (1.4 SD above mean, Fig.3H). These temporal CSF profiles indicate initial release of uncleaved and partially digested GFAP with subsequent decline, while smaller fragments displayed prolonged elevation, supporting progressive GFAP proteolysis over 10 days. To further address the broad variances observed, we next inspected correlations among GFAP species in this cohort.

### Fragment covariation supports overall GFAP “injury burden” and imbalance of small GFAP-BDPs

Because multiple factors influence GFAP release and proteolysis, we next examined bivariate correlations among GFAP bands to distinguish overall GFAP acute “injury burden” in CSF from fragment-specific behavior in this cohort (Fig. 4A). Strongest covariance was observed between 50kDa GFAP uncleaved substrate and 37-39kDa fragments (r=0.82, R^2^:0.68, LogWorth=24.3; Fig.4B). The 40-42kDa bands correlated similarly with the 37-39kDa fragments (r=0.73, LogWorth=16.3), plausibly reflecting co-release of precursors and intermediary products. The 15-19kDa bands showed weaker relations with precursor bands, and had 28% more undetected values than the 20–26kDa BDPs. Consequentially this lowered the correlation between these presumed co-products (Fig.4C). Overall, strong covariance across most bandsets indicates robust within-sample match of multiple GFAP species, supporting an general patient-specific GFAP injury burden concept. To the contrary, the selective absence and weaker relations of the 15-19kDa bands with other fragments adds evidence of imbalance. This pattern is consistent with the coil1/coil2 imbalance observed by MS.

These findings are further explored in a trauma culture model. Given both uncleaved and fragment GFAP species in our dataset, we next asked which GFAP trajectories would better correlate with recovery after moderate-severe TBI.

### GFAP proteolysis, but not bulk GFAP release signal, correlates to TBI outcome

This CSF dataset provides the opportunity to assess whether overall GFAP release, or its postinjury proteolysis, correlate to TBI recovery. Longitudinal CSF levels of full-length GFAP and its fragments were compared between patients with good versus poor six-month outcomes using GOSE. Age- and time-adjusted CSF means of 37-39kDa GFAP-BDPs were higher in patients with poor outcome (GOSE 1-4) than in those with good outcome (GOSE 5-8, Fig.5A, effect sizes given in Table 1B). Small 20-26kDa BDPs separated good versus poor GOSE early after injury, showing lower levels in patients with favorable outcome, whereas their means converged during later PIDs (Fig.5B). In contrast, temporal profiles of full-length GFAP and large fragments showed substantial overlap of mean spline confidence bands between outcome groups, with similar adjusted means. Similar results were observed when applying a stricter partitioning of favorable outcome (GOSE7+8), indicating these findings were robust to different ways of GOSE dichotomizing (Supplemental Table S8). Considering that full-length and end-clipped GFAP constituted the majority of the “total” GFAP signal measured in CSF, these data suggest that progressive proteolysis, rather than overall GFAP release signal, captures injury processes that are relevant to six-month recovery.

### Mechanisms of GFAP proteolysis and modification in a human astrocyte trauma model

A series of epitope-defined GFAP antibodies was used to map coil1 and coil2 products, track their compartmental distribution, and define the center cleavage site that generates them after trauma. Protease inhibitors and live-cell reporters then documented calpain and caspase contributions to traumatic GFAP degradation in membrane-wounded human astrocytes, lifted debris and fluid-released compartments. Lastly, neoepitope-coil2-directed citrullinated GFAP staining showed this GFAP-BDP proteoform as non-filamentous aggregates within pathological astrocytes, providing further evidence for selective fragment retention.

### Trauma-generated GFAP coproducts are destined to different locations

Primary fetal human cerebral astrocytes were in vitro differentiated on elastic membranes and were reproducibly injured by abrupt pressure-pulse stretching [95]. In this trauma model, GFAP filaments are disassembled, degraded and released [33]. To identify coil-specific fragments, mabs with defined GFAP epitopes were used to probe immunoblots of whole cell lysates (“cells”) and concentrated culture medium (“fluid”) from unstretched and stretched cultures of same donors (Fig.6). Epitopes of these clones are aligned with coil-specific fragment sequences, using colors matching MS-derived maps (Fig.6AB). All clones detected full-length GFAP and large fragments but differed in detecting smaller bands. Clone 3E10 detected a 20kDa band in culture fluid that increased after trauma yet failed to detect a corresponding band in cells (Fig.2C, red box, left). Clone GA5 recognized an epitope at the end of coil2, at the beginning of the GFAP tail [18]. It detected a set of 17-28kDa bands in cells but not in fluids (Fig.6C, purple box, right). A cocktail of three mabs served as positive control, confirming the presence of smaller fragments in cells and fluids from same donors (Fig.2C, blue box). Thus, following mechanical trauma of human astrocytes, coil1-harboring products were preferentially released into fluids, while coil2-containing products were selectively retained within cells. This new insight from the human trauma model supports the findings of enriched coil1-BDPs in biofluids of TBI patients, compared to coil2-BDPs.

**Fig. 6.**
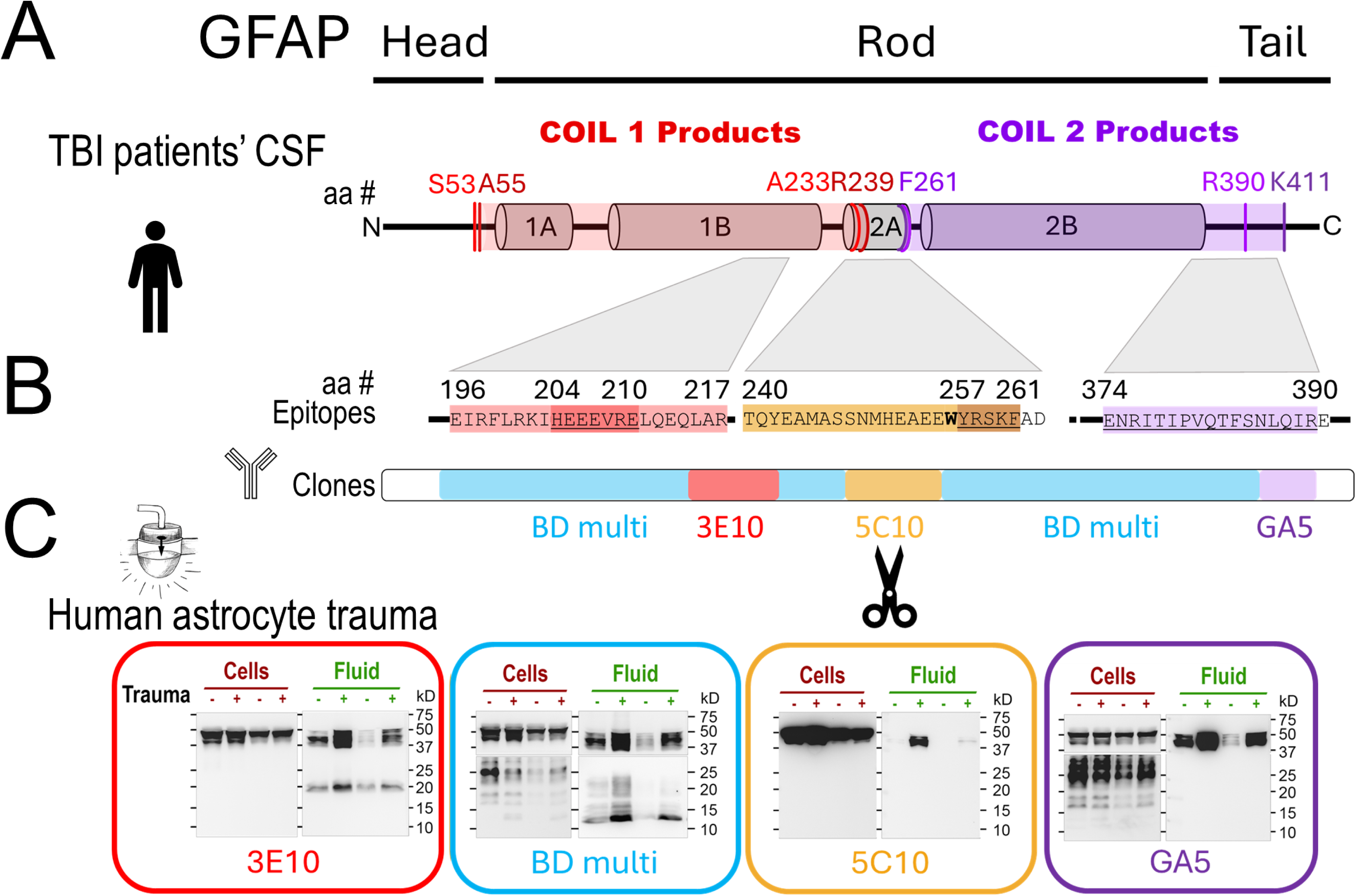
Novel GFAP cleavage creates coproducts with distinct compartmental fate after trauma. Monoclonal antibody mapping confirms the in TBI-sequenced GFAP co-products. Antibodies also identified center cleavage and document distinct compartmental fate of GFAP co-products in a human astrocyte trauma model. **A)** MS-derived coil1/2 sequences are aligned with the epitopes of used antibodies. **B)** Binding regions of GFAP monoclonal antibody (mab) clones: Mab 3E10 targets coil1B (aa 196-217) with epitope HEEEVRE (aa204-210, “stutter” motif). Mab 5C10 targets coil 2A (aa 240-260) with newly deciphered core epitope (aa 257-260, YRSKF). Mab GA5 binds near the end of coil 2B (aa 374-390)[22]. The BD multiclone (4A11, 1B4, 2E1) recognizes both coils. **C)** Fluid and cell lysate immunoblots document that coil1-products were released into fluids, whereas coil2-products remained cell-associated. Two donor’s whole cell lysates (Cells) and concentrated astrocyte-conditioned medium (Fluid) from control (-) and 2 days post-stretched human astrocyte cultures (“+”, 4.8-5.4psi) were probed with antibodies listed in B. All clones detected full-length GFAP and large fragments in Cells and Fluids. 3E10 detected 20kDa products in Fluids but not Cells. BD Multiclone detected intracellular 13-28kDa GFAP-BDPs and small bands in Fluids. 5C10 did not detect any small fragments, and GA5 detected 17-28kDa GFAP-BDPs in Cells, but not in Fluids.

### Mapping a novel GFAP trauma cleavage site by defining an antibody epitope with selective chemical scission

Clone 5C10 detected full size and large GFAP-BDPs in cells and fluids, but no fragments smaller than 40kDa in either compartment (Fig.6C, yellow box). This suggests that 5C10 recognized an epitope that was destroyed during further trauma proteolysis of GFAP. Thus, identifying this antibody’s epitope would inform where GFAP was cleaved after trauma. To pinpoint the 5C10 epitope to a specific position within the center of the rod domain, we performed a chemical-mediated partial digest of GFAP using BNPS-skatole that cleaves after tryptophan residues, which there is only a single tryptophan in GFAP (W256) [34, 92]. BNPS-skatole-treated GFAP produced two distinct products, a N-terminal 34.6kDa product and a C-terminal 20.5kDa product, and 5C10 bound avidly to the ∼20kDa product on western blots (Supplement Fig.S9C). The identity of this ∼20 kDa BNPS-skatole fragment was confirmed using anti-IFA, a pan-intermediate filament antibody with a defined epitope in coil2B [53, 77]. The coil1 antibody 3E10 confirmed that the larger BNPS-skatole fragment corresponds to the N-terminal BNPS-skatole product (Supplemental Fig.S9D). Together, these two controls with 5C10 indicated that the sequence immediately following W256 from Y257 to F261 is a critical component of the 5C10 epitope (YRSKF, brown epitope map Fig.6B). Because 5C10 does not detect small GFAP-BDPs after trauma, this places the trauma-induced cleavage site within this region. It likely involves elements of the neighboring caspase-3 cleavage site (YRSKFA↓DLTD). MS analysis from TBI-patients showed that coil1-BDP sequences extended no further than R239, whereas coil2-BDPs consistently begun at F261, and thus pins the center trauma cleavage of GFAP within this area. These antibody-mapping data therefore identify a novel trauma-cleavage upstream of F261 with potential caspase-3 acting downstream of this position.

Calpain and caspase activation is established in rodent trauma models [5, 23, 51]. To determine their contribution to generating small GFAP-BDPs in human traumatized astrocytes, we applied inhibitors and live-imaging reporters in the human trauma model.

### Generation of small GFAP-BDPs in human traumatized astrocytes involves calpains and caspases

Human traumatized astrocytes were treated with calpain inhibitor calpeptin, or caspase inhibitor ZVAD-FMK, or both. Two days later lysates of adhered and lifted cells as well as fluids were analyzed for cytoskeletal degradation. The positive control was αII-spectrin, which generates defined well-known calpain and caspase spectrin breakdown products (SBDPs) and was expressed in human astrocytes [49]. Both calpain and caspase SBDPs increased two days after stretch-injury in cells, lifted material, and fluids, and decreased with respective inhibitors (Supplemental Fig.S10). Of note, SBDPs displayed selective compartmental distributions. The 120kDa caspase-SBDP was particularly enriched in detached material of demised cells. The cellular calpain-double-band (145/150kDa) separated with the 150kDa SBDP found in lifted cells, while the 145kDa SBPD was detected in the fluid. These findings suggest that intracellular αII-spectrin digestion may also result in distinct cell- and fluid-destinies after trauma.

With specificity and effectiveness of these inhibitors confirmed in the human model, we next examined trauma-induced GFAP-BDPs. Two days after trauma without inhibitors, GFAP-BDPs were elevated in cells, lifted material, and fluid shown on immunoblot bands and densities are graphed for 35-38kDa and 19-26kDa bandsets across compartments (Fig.7AB). Calpeptin and ZVAD-FMK both separately reduced 35-38kDa BDP, and co-application produced an additive effect. The smaller BDPs (19-26kDa) decreased with calpeptin and diminished further with ZVAD-FMK, and combined drugs eliminated these bands across all compartments. These preliminary human trauma model findings parallel rodent injury model data, indicating that both calpains and caspases were involved in trauma-driven GFAP degradation [23]. Resting astrocytes had low-level GFAP-BDPs, consistent with baseline cytoskeleton turnover, and displayed similar inhibitor sensitivities [75]. Full-length GFAP increased in lysates after trauma, consistent with increased solubility of monomers by breakage of insoluble filaments (Supplemental Fig.S11A). Uncleaved 50kDa accumulated further with calpain inhibition, supporting substrate buildup (Supplemental Fig.S11B). This data provides initial evidence of endogenous calpain- and caspase-driven GFAP cleavage in human traumatized astrocytes extending prior rodent work [103].

**Fig. 7.**
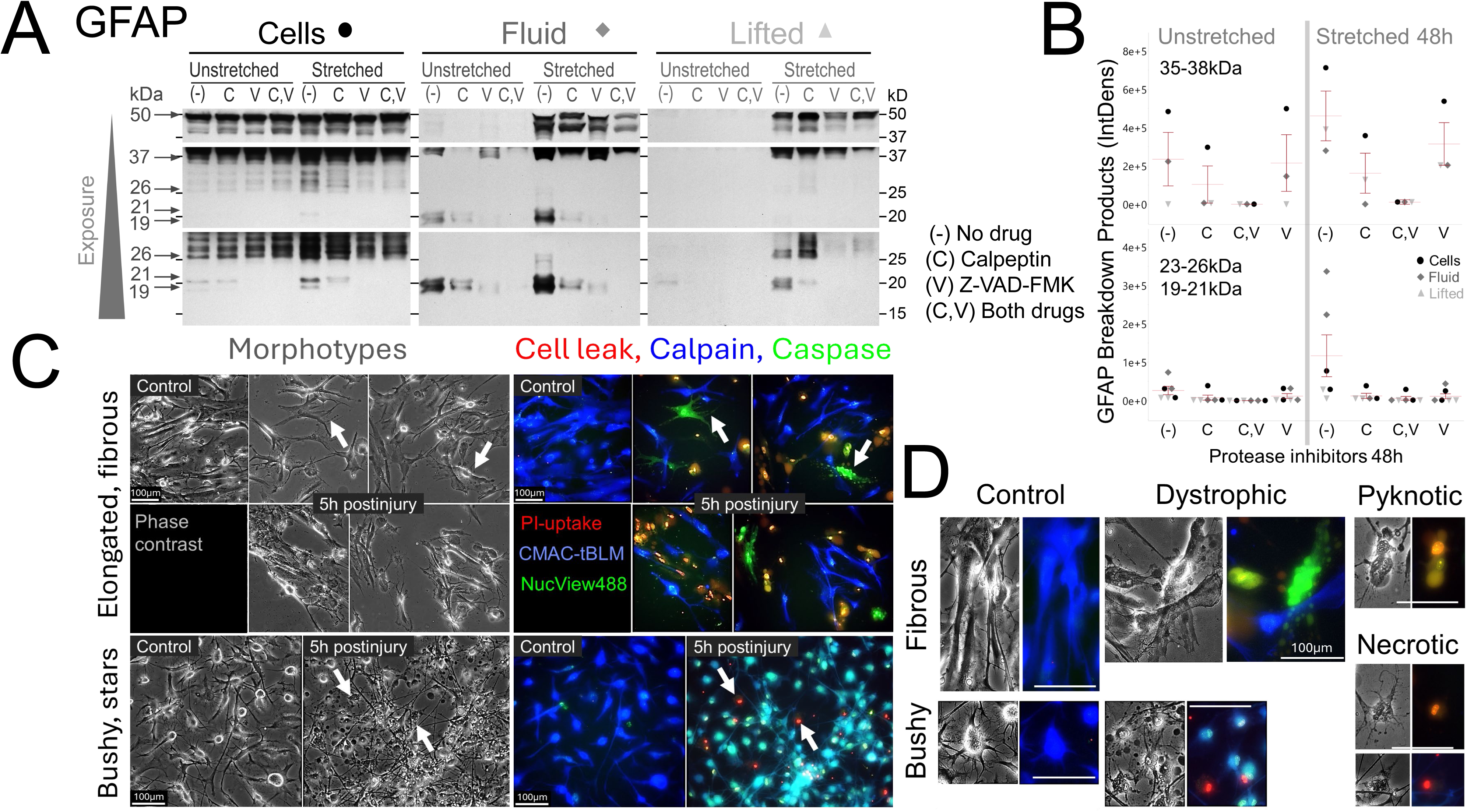
Calpain and caspase activities in compartmental GFAP degradation and astrocyte phenotyping **A)** Shown is GFAP degradation at 48h postinjury across adherend cells, released pool and lifted material with or without calpain and caspase inhibitors. Unstretched and stretch-injured human astrocytes were treated with calpeptin (C, 13µM), Z-VAD-FMK (V, 11 µM), or both (C,V) and whole cell lysates (Cells), conditioned medium (Fluid), as well as pelleted and then lysed lifted material (Lifted). Samples were then analyzed for GFAP degradation. Immunoblots (Mab 3E10) of increasing exposures show strongest BDP signals after stretching without inhibitors (-) and BDPs were variably reduced with inhibitors. **B)** Densities of 35-38kDa BDPs (top) as well as 19-22 and 23-26kDa (bottom) BDPs are graphed across all compartments shown in A. Calpain-inhibition reduced all shown BDPs, while caspase inhibition more clearly reduced small products, and combined drugs abolished signals of both GFAP-BDPs bandsets. Resting samples showed baseline 35-38kDa bands and low-level 19-26kDa bands, with all fragments showing inhibitor suppression. **A)** Live imaging of human astrocyte morphotypes uninjured and 5-6 hours postinjury show injury stages defined by membrane permeability with protease activity. Phase contrast shows elongated, aligned (top), or star-shaped tiling (bottom) in control astrocytes. These uninjured astrocytes had intact processes. Immunofluorescence shows calpain activity (CMAC-tBOC-leucyl-methionine, blue). Phase images of stretched cultures displayed morphologically damaged and blebbing cells (arrows). Subgroups had permeable membranes detected by PI-uptake (red/yellow) that were devoid of calpain, but positive for caspase activity (NucView488, green, fibrous cells), or were active for both proteases (tiel, star-shaped cells). **B)** Zoomed-in views show individual control fibrous and star-shaped morphotypes with calpain activity. Stretch-injured dystrophic astrocytes display structural process damage with thin, truncated, or beaded clasmatodendrotic processes that were either caspase active and calpain-inactive (fibrous type) or retained both protease activities (star-shaped type). Among the subgroup of cells with advanced integrity-compromise were pyknotic morphologies with nuclear caspase activity (pyknotic) and cells devoid of protease activity (necrotic). Phase-dense vacuoles were unstained (see Supplement Fig. S13 for individual channels). Scale bars: 100 µm.

Given the limited knowledge of human astrocyte injury states compared with the expanding literature on GFAP as a TBI biomarker, we next combined live-cell protease activity reporters with membrane-leak dependent dye uptake to connect cell injury with proteolysis. We lastly determined fixed citrullinated GFAP immunofluorescence to further characterize astrocyte injury phenotypes. These initial microscopy-based pathophysiological studies begin to characterize human traumatized astrocyte injury states.

In this model GFAP fluid elevation was previously analyzed over time and correlated to the percentage of cell wounding and cell death [33]. Here temporal postinjury of two band-specific fragments are reported. Significant 37kDa fragment elevation in fluid occurred already at 0.5 and 5h (p<0.05 Cohen’s d=1.1) that increased further by 48h (p<0.01, d=3.4; 4-9 donors, Supplemental Fig.S12B). The 20kDa product release was elevated by 24h (p<0.05; d=1.6) and consistently at 48h postinjury (p<0.001, d=3; Supplemental Fig.S12C). These quantified band-specific release profiles support progressive proteolysis following astrocyte trauma.

### Human astrocyte pathophysiology relates to membrane wounding and proteolysis

Acute astrocyte injury states after stretch trauma were examined using live-fluorescent calpain and caspase reporters together with PI to assess membrane permeability. Differentiated human astrocytes displayed two morphotypes, fibrous elongated cells and bushy, star-shaped cells, resembling white and gray matter astrocyte morphologies [1] (Fig.7CD). Under control conditions, both subpopulations showed calpain but rarely caspase activity, consistent with fetal neocortical expression of calpains and low levels of calpastatin [81].

Five hours after stretch injury, a mix of astrocytes with intact and permeable membranes document selective plasma membrane damage as quantified before [33]. Injured fibrous astrocytes lacked calpain activity and frequently displayed beaded, clasmatodendrotic processes with caspase activity (Fig.7CD) [19, 33]. Among injured bushy astrocytes were many cells displaying simultaneous membrane leak and protease activity, will majority showing calpain and caspase activity (individual channels see Supplement Fig.S13). These injured cells lost radiating processes and displayed blebbing with phase-dark vacuoles devoid of enzyme activity (Fig.7C). Acutely pyknotic cells, identified by phase-dense appearance and shrinkage, showed advanced integrity loss with bright PI-stained nuclei in both subpopulations. These pyknotic cells were either protease-inactive (red), consistent with necrosis, or caspase-active and integrity compromised (yellow), suggesting apoptotic phenotypes (Fig.7CD). These initial observations show distinct concurrent degenerating phenotypes of acutely injury human astrocytes with protease activity and membrane wounding hours after trauma illustrating a spectrum of early vulnerability.

This model offers an opportunity for future analyses of associated aberrant proteoforms, although our studies to date have been limited to antibody probes.

### Citrullinated GFAP forms non-filamentous aggregates that identify pathological astrocytes after trauma

Distinguishing filament-assembled from aberrant GFAP proteoforms in traumatized astrocytes is critical to begin addressing astrocyte pathology beyond classical reactive astrogliosis. As indicated above, coil2-GFAP-BDPs were retained in cells, hence we chose two coil2-specific antibodies to compare unmodified and citrullinated GFAP forms: the rabbit mab cit270-GFAP, recognizing a citrullinated neoepitope in coil2, and mab 5C10 with epitope beyond coil2 in uncleaved GFAP. Cit270-GFAP signals were granular, not filamentous, a specific pattern that was confirmed by absence of granules with rabbit IgG negative control, while a polyclonal GFAP antibody labeled cytoskeletal filaments serving as positive control for GFAP expression (Supplemental Fig.S14A). Cit270 antibody stained clusters, speckles and large granules, but did not stain cytoskeletal fibers that were co-stained for unmodified GFAP, emphasizing that citrullinated GFAP aggregates are structurally distinct from filament-assembled GFAP (Fig.8). Stretch-injured cultures showed larger and more abundant cit270-GFAP clusters than controls both at 5 and 24h postinjury (Fig.8, see Supplemental Fig.13BC for separate channels). Cit-GFAP stained either cells that were negative or excessively bright stained for classic GFAP. Increased GFAP antigenicity is associated with filament-disassembly following acute wounding, calcium ionophore-exposure, or calpain activation in astrocytes [33, 58]. Astrocytes with cit-GFAP granules displayed morphotypes of reactive, shrunken, or swollen dystrophic astrocytes devoid of processes (Fig.8B, Supplemental Fig.S14B-D for individual channels). Cit270-stained puncta were also peppered around astrocyte processes that appeared extracellular, as delineated astrocyte shapes on phase-contrast images showed (Supplemental Fig14E).

**Fig. 8.**
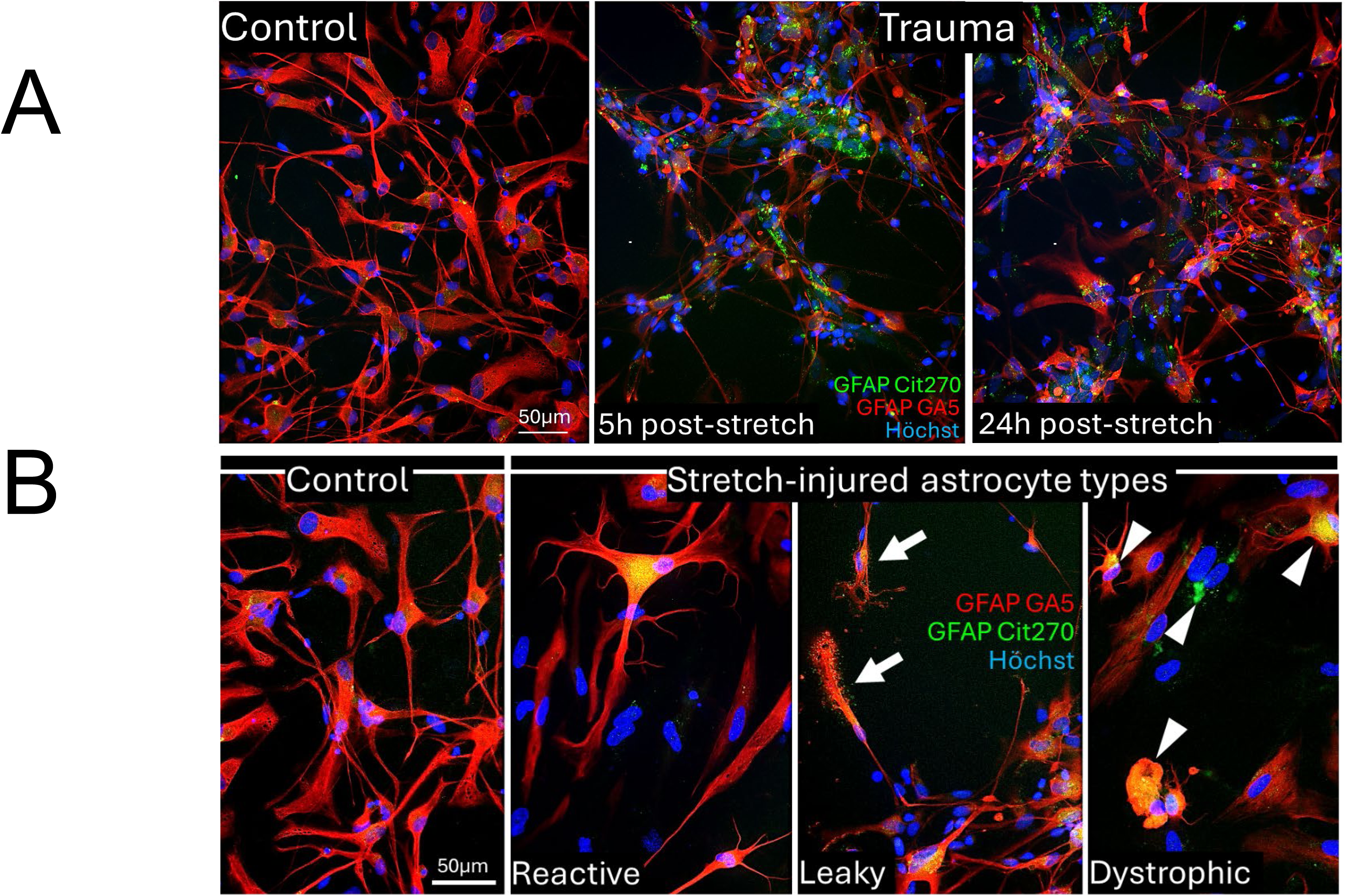
Trauma induced citrullinated GFAP aggregates in pathological human astrocytes **A)** Immunofluorescence of unmodified coil2 GFAP (GA5, red) and citrullinated coil2 GFAP (cit270-GFAP, green) shows abundant accumulation of non-filamentous citrullinated GFAP clusters at 5 and 24h post-stretching (right), and little to no citrullinated GFAP staining in unstretched control astrocytes (left). **B) Control** astrocytes (left) and three pathological astrocyte morphotypes (right) after stretch-injury: **Reactive** astrocyte shows typical thick, hypertrophied processes, bright cytoskeletal GFAP and fine-grained cytosolic citrullinated GFAP (yellow). **Leaky** astrocytes display fanning processes surrounded by citrullinated GFAP sprinkles (arrows). **Dystrophic** astrocytes with swollen cell bodies and malformed, truncated or missing processes contain larger citrullinated GFAP aggregates and had non-filamentous or malformed GFAP cytoskeletal fibers or were devoid of unmodified GFAP signals (arrowheads; see Supplement Figure S14 for individual channels and phase images). Scale bars: 50 µm.

Thus, mechanical injury and filament disassembly was associated with abundant GFAP citrullination of assembly-defective GFAP proteoforms in aggregates that identified pathological astrocytes. These findings provide evidence for trauma-associated coil2-aggregation that may contribute to its selective intracellular retention and that led to coil2 underrepresentation in culture medium and TBI patients biofluids. They further establish citrullinated R270-GFAP-BDPs as a new markers to study astrocytopathy and astrocyte degeneration relevant to traumatic pathophysiology.

In summary, TBI patients show GFAP destabilization by citrullination and acetylation, and trauma-specific proteolysis producing unique small co-products with distinct temporal release behaviors, explained by citrullination-modified and assembly-defective GFAP aggregation in pathological astrocytes. CSF trajectories of GFAP proteolysis, not bulk release of uncleaved GFAP carried outcome distinction, underscoring the value of monitoring postinjury degradation processes as being part of disturbed proteostasis in TBI patients.

## DISCUSSION

The principal findings of this study are that following a TBI proteolysis generates unique GFAP-BDP proteoforms with distinct temporal profiles that associate with clinical outcome. The results indicate that GFAP biomarker profiles reflect downstream injury processes with perturbed astrocyte proteostasis rather than a primary trauma insult alone. Moreover, the assembly-defective GFAP proteoforms are unique to TBI, and distinct from GFAP fragments seen in other neurodegenerative diseases. These data support a concept that disrupted proteostasis may contribute to an astrocyte proteinopathy with potential consequences for neurodegeneration and patient recovery. These initial mechanistic insights provide a framework for future studies linking GFAP proteolysis and long-term post-traumatic neurodegeneration.

### Unique GFAP-BDPs in TBI initiated by mechanical force

Our results show that the GFAP-BDPs are unique to TBI and distinct from other neurodegenerative diseases such as AD or Alexander disease. This is an important feature of this work because it argues against the concept that TBI triggers AD related processes but instead involves distinct, trauma-specific molecular cascades and targets that are summarized elsewhere [33]. Direct trauma-induced protein changes are shown by high-speed biomechanical force stretching intermediate filaments to disassemble or break exposing the rod-domain [11]. Stretching of rat brain astrocytes activates PAD2 expression and protein citrullination [2]; and membrane injury induces localized calpain activation [46], both in place to destabilize and make GFAP susceptible to further core cleavage.

### The uniqueness of the GFAP co-products appears to be related to a coil 2A cleavage sensitivity

Cleavages that produce TBI-associated GFAP-BDPs are distinct from established protease sites described in other neurodegenerative diseases. A caspase-6 GFAP cleavage (VELD_225_-VAKP) has been reported in mouse culture models of Alexander disease, splitting GFAP within linker2 [3, 16, 18]. Here we identified longer coil1-products with a TBI-cleavage map between residues 240-260. GFAP-BDPs reported in AD disease are generated by a caspase-3 cleavage at _258_RSKFA↓DLTD [66]. Our biochemical and epitope mapping studies instead indicate _257_YRSKF_261_ as the likely cleavage region flanking this caspase-3 site that generates the observed new N-terminal coil2-product, which is shorter than the reported AD fragments. However, the shorter TBI coil1-product end cannot be fully explained by this cleavage. Coil2A appears particularly sensitive to proteolysis after TBI and our data suggest involvement of calpains and caspases, although other proteases, including possible endogenous trypsin-like activity, could contribute to the novel coil1 terminus (LTAALKEIR↓T; R239). Endogenous trypsin activity has been described after brain injury, compromising astrocytes [28, 101]. Joint protease activities can account for the observed sequence gap within coil2A, between the two product sets, warranting further studies.

### Structural evidence for protease sensitivity of the TBI cleaved region

Intact GFAP molecules anneal via α-helical coiled-coil interactions to form parallel dimers, antiparallel tetramers and ultimately 10nm diameter units that grow in length to mature filaments [47]. Computer modeling and structural data show that the entire rod segment of vimentin, a close homolog to GFAP, adopts a continuous α-helix [29]. This is despite several regions, historically referred to as linkers or “stutters”, that lack the typical hydrophobic heptad repeat pattern of α-helical coiled-coils. These regions contain proline and other turn-promoting residues or incorporate 11-residue (hendecad) sequences that disrupt the heptad repeats and are therefore thought to represent non-helical linkers. The Alexander disease-associated GFAP cleavage falls within one such non-helical region (“linker12”). Notably, the region containing the 5C10 epitope and caspase-3 site is historically referred to as “stutter1”, where a hydrophobic heptad repeat is replaced by a single hendecad sequence, disrupting the local packing and rendering this segment particularly susceptible to proteases [86].

### Consequences of GFAP modification, degradation, and aggregation as a possible astrocyte proteinopathy in neurodegeneration

Assembled GFAP filaments are essential for astrocyte morphology, mechanical resilience, ATP release and other key functions [43]. Low baseline calpain activity contributes to cytoskeleton remodeling and is thus consistent with our data in resting astrocytes [75]. Since cytoskeleton integrity provides structural resilience and metabolic competence, excessive GFAP proteolysis can exacerbate astrocyte vulnerability. Pathological GFAP proteoforms can critically undermine astrocyte function, compromise glutamate buffering, and neurovascular unit integrity, which matter for functional network activity and synaptic plasticity. In our human astrocyte trauma model, we demonstrate pathological changes associated with these GFAP-BDPs. Specifically, the trauma-driven citrullinated GFAP aggregates in pathological astrocytes suggest that trauma-induced GFAP proteoforms could potentially represent a harmful proteinopathy, and at minimum concurred in swollen dystrophic astrocytes.

### Significance and implication of biofluid and pathological astrocyte GFAP breakdown products

Proteolysis of GFAP after TBI produces unequal abundance of BDPs in patient biofluids, due to differences in release, or in susceptibility to further degradation, or both. We show preferential coil1-release and selective coil2-retention as non-filamentous citrullinated GFAP aggregates in traumatized human astrocytes. An in vivo study documents strongly upregulated citrullination in the contused rat brain localized to astrocytes [56]. Thus, citrullinated GFAP aggregates inform on an GFAP fluid biomarker-underlying astrocytopathy.

TBI CSF samples carry both co-products, yet coil2-BDPs were either absent or elevated, depending on the patient studied. This varied finding could involve adverse patient events causing glial integrity loss, with release of otherwise retained proteoforms.

On one hand, fluid coil1-GFAP may be matched by a latent coil2-GFAP aggregation burden in the injured brain. On the other hand, once coil2 products are detected in fluids, this may signal worsening conditions with increasing astrocyte demise. Membrane damage, blebbing and swelling, can lead to oncotic cell death seen as cytotoxic edema and delayed cell death [59, 93]. Thus, use of coil- and citrullination-specific GFAP antibodies, as those characterized here, can serve as pathological astrocyte markers, and as tools for TBI assays. Possibilities for future research that address clinical needs include: (1) Examination of citrullinated GFAP-aggregates in postmortem brains in the context of post-traumatic neurodegeneration to further investigate traumatic astrocytopathy. (2) Larger cohort studies to validate GFAP-BDP proteoform profiles along with linkage to disrupted proteostasis and posttraumatic neurodegeneration [38].

### Relevance of GFAP-BDPs for assay design and interpretation

GFAP testing following TBI is already incorporated into large multisite clinical studies for acute patient assessment using commercial antibody-based platforms [6, 7]. However, beyond general specificity for GFAP, it is unclear which proteoforms are being measured in these assays, as this information is typically proprietary or unknown and it is presumed that total released GFAP is captured. The distinct release and modifications of coil1 and coil2 products can therefore inform both the design and interpretation of GFAP assays.

### Potential for new GFAP ‘contexts of use’ in TBI monitoring

Our sequence-defined GFAP degradome and connected cytological data support possible future clinical applications of GFAP in longitudinal TBI patient monitoring. Three interrelated potential contexts of use (COUs) are considered: *(1) Longitudinal monitoring*; (*2) Astrocyte degeneration and secondary injury; (3) Outcome prognosis*.

#### (1) Longitudinal GFAP monitoring post-TBI controlling for age

Tracking GFAP degradation kinetics offers a longitudinal window into TBI patient monitoring during intensive care for early clinical trial endpoints [26]. GFAP-BDPs are abundant in the blood, while CSF carries both uncleaved GFAP and BDP fragments [33, 63, 108]. CSF GFAP analysis shows an initial post-injury abundance of uncleaved and end-clipped GFAP (e.g., missing N- and C- termini), where both are detected at peak elevation in the first 24 hours from injury. This temporal pattern is similar to the kinetics of other TBI biomarkers, including UCH-L1, S100β, brain lipid binding protein, aldolase C, neuron-specific enolase and SBDPs [33, 96]. Although with different half-lives, they share primary injury-related release associated with acute mechanoporation and necrosis [33, 72, 76, 87].

Secondary elevations of GFAP-BDPs, particularly of small products, involve caspase activation and occur in a subset of TBI patients. This pattern aligns with reports of delayed serum GFAP peaks in some older TBI patients with high CT lesion burden, poor outcome, high mortality and features of more severe TBI [79]. Age-related increases in baseline GFAP and neurofilament light (NfL), two intermediate filaments with pronounced trauma-associated fragmentation, associate with ongoing atrophy during neurodegeneration [15, 30, 82, 107]. Elderly TBI patients may also clear CSF less efficiently due to decreased CSF pulsation, vulnerability of ependymal cilia, and disrupted cerebral blood flow, all of which can influence biomarker levels [12, 48, 100]. Taken together, long-term monitoring of GFAP fragments could add further insight into secondary TBI pathophysiology controlling for age-related reference values [20, 89] .

#### (2) Secondary adverse events after TBI may reflect delayed astrocyte pathology

Patient-specific secondary GFAP-BDP release can occur during injury progression. Neuroimaging reports that contusions expand into peri-lesional tissue, a process attributed to cytotoxic edema [69, 88]. Cytotoxic edema is characterized by microstructural changes of astrocyte swelling and subsequent integrity loss, driven by ion dysregulation and metabolic depression [91, 93]. These interrelated mechanisms may underlie variable secondary elevation of cell death-associated GFAP fragments observed in some TBI patients. Consistent with this interpretation are data from progressive astrocyte degeneration, quantified in autopsy brains of TBI patients, showing reduced astrocyte numbers between acute (<6h) and subacute (>6hrs to 3PID) intervals in cerebral white matter and hippocampus [59]. Thus, GFAP-BDP profiling may indicate secondary postinjury events after TBI.

#### (3) GFAP-BDP trajectories for evaluating TBI outcome

Elevated GFAP is widely recognized as a predictive biomarker after severe TBI [33, 52, 73, 79]. Our findings indicate that in CSF the time course of GFAP-BDPs distinguishes poor from good 6-month outcome, rather than total GFAP signal, which is based on uncleaved and end-clipped GFAP in CSF. This finding was robust to different threshold scores for dichotomizing GOSE, which is relevant as recovery evaluation is individual [97]. The results suggest that monitoring temporal GFAP proteolysis can provide predictive information, which may be useful in GFAP assays with binding agents that could target specific BDP fragments. This thesis aligns with the broader clinical understanding that early injury progression, rather than the initial injury status alone, shape recovery [10, 102]. Because GFAP testing is currently only FDA-approved for an early limited post-injury window for evaluation within ∼12 hours, and thus is not positioned to assess the temporal evolution of GFAP proteolysis through serial sampling. Yet longitudinal TBI patient assessment is critical to control events affecting their recovery [52].

### Limitations of the study

The clinical cohort used to establish the current findings was relatively small, and included patients that had incomplete longitudinal data, due to staffing and logistical constraints during the COVID-19 pandemic. This limited statistical power and warrant future studies in additional TBI patients to validate the findings. Steps were taken to mitigate this, including truncating collection at 10 days post-injury, to avoid imbalance in variances, and by using appropriate effect size metrics. Effect sizes were calculated using Cohen’s d to injury day differences from means over time; as well as for time- and age-adjusted differences between good versus poor outcomes across all GFAP band sets. Receiver operator characteristic modeling with area under the curve estimate statistics is inadequate to account for elapsed postinjury time, while effect sizes allowed adjusting for age and elapsed time simultaneously. Citrullination and acetylation of MS sequenced GFAP-BDPs from TBI patients was rigorously validated. However, immunoblot contrasting of citrullinated GFAP with unmodified GFAP was only done on selected samples due to limited availability of the citrullinated GFAP neoepitope antibodies underscoring the need for larger cohort analysis. Technical restrictions limited serum MS sequencing. CSF collection in TBI patients undergoing ventriculostomy offers unique biomarker insights but is limited to ICU studies and does carry a risk of infection [41, 70].

Cerebral astrocytes isolated from donated fetal brain specimens offer a unique opportunity to investigate human primary brain cells responding to injury using stretchable culture plates, but these labor-intensive experimental protocols limited the number of multi-donor assessments. Simultaneous live-protease activity and cell integrity studies provided mechanistic insights on mechanoporation injury-driven changes in proteolysis, but additional donors are needed for more detailed protease specific inhibitor effects on individual GFAP fragments. Fetal brain-derived astrocytes may only be partially representative of adult or aged astrocyte populations. *In vitro*-differentiated human cortical astrocytes used here display complex morphotypes and express markers of differentiated astrocytes, making this a useful human model [33, 95, 105]. Fluorescence live dye imaging did not allow tracking over time, due to high-intensity illumination. Despite these challenges, all key measurements were obtained comprehensively across samples, using appropriate controls and unbiased processing. Importantly, findings were cross validated using independent experimental approaches. As this is a discovery-focused study, rigorous orthogonal validation supports the reliability of the results. Certainly, larger standardized clinical TBI patient cohorts and translational preclinical studies across centers will be critical for future validation to extend the reported findings [94].

## CONCLUSION

### Translational significance and future directions

Collectively, our data positions GFAP-BDPs as molecular indicators of disrupted proteostasis and astrocyte pathology. Future work will aim to link citrullinated GFAP proteoforms with astrocytopathy in human brain tissue after TBI and examine their potential relationships with neurodegenerative conditions. Serial biofluid measurements of GFAP-BDPs may support patient monitoring in relation to physiological parameters and secondary in hospital insults occurring in neurocritical care units. By connecting GFAP biofluid signatures to citrullinated GFAP fragment aggregates, this work contributes to a translational framework for TBI phenotyping that could help to advance patient care. These dynamics provide new perspectives for investigating GFAP, and specifically GFAP-BDPs as a biomarker with the potential for latent proteinopathy in pathological astrocytes, which could bridge TBI and posttraumatic neurodegeneration.

## ABBREVIATIONS

Alzheimer’s disease (AD), amino acids (aa), acetonitrile (ACN), adjusted means (Adj.Means), breakdown products (BDPs), confidence intervals (CI), Conditioned medium (CM, “fluid”), CMAC-tBOC-leucyl-methionine (CMAC-tBLM), context of use (COU), cerebrospinal fluid (CSF), coefficients of variation (CV), degrees of freedom (df), denominator term (DFDen), Enhanced Chemiluminescence (ECL), false discovery rate (FDR), fixed effect statistic (F-statistic), Glasgow Coma Scale (GCS), glial fibrillary acidic protein (GFAP), Glasgow Outcome Scale (GOS), Extended Glasgow Outcome Scale (GOSE), higher-energy collisional dissociation (HCD; MS2), patient identifier (ID), interquartile range (IQR), Institutional Review Board (IRB), liquid chromatography-tandem MS (LC-MS/MS), linear mixed model (LMM), mouse monoclonal antibodies (mabs), mass spectrometry (MS), full mass spectra (MS1), National Institute of Neurological Disorders and Stroke (NINDS), peptidylarginine deiminase 2 (PAD2), Proteome Discoverer (PD), propidium iodide (PI), postinjury day (PID), posttranslational modifications (PTMs), restricted maximum likelihood (REML), Research Resource Identifier (RRID), room temperature (RT), αII-spectrin breakdown products (SBDPs), SDS polyacrylamide gel electrophoresis (SDS-PAGE), standard error (SE), Tris-buffered saline with Tween (TBST), Traumatic brain injury (TBI), Whole cell lysate (WCL, “cells”), partial omega squared (ω_p_ ^2^), University of California, Los Angeles (UCLA), UF Health Shands Hospital, University of Florida, Gainesville (UF).

## DECLARATIONS

### ETHICS APPROVAL AND CONSENT TO PARTICIPATE

For deidentified TBI patient’s CSF and serum collection with metadata, adult patients (>18 years of age) were enrolled at two Level 1 trauma centers, in Los Angeles, California, at the UCLA Ronald Reagan Hospital Neuro-Intensive Care Unit, and in Gainesville, Florida (UF Health Shands Hospital, University of Florida). The human subjects’ protocols included local IRBs that approved informed consent procedures at UCLA under Chair Paul Vespa and at the University of Florida Dept of Anesthesiology under Dr. Steven Robicsek. The Study “Biochemical markers for traumatic brain injury tissue and data bank” was approved by the Institutional Review Board (IRB) of the University of Florida with the IRB# 201703310. Patients met inclusion criteria if they were 18 years old with a non-penetrating head injury and had a GCS <8 requiring the placement of an intraventricular catheter. Patients were excluded if they had a history of preexisting end-stage organ disease or severe psychiatric illness. At UCLA, the Office for Human Research Protection (OHRP) approved the informed consent protocol for the Brain Injury Research Center study with the BIRC-IRB# 10-0929.

For deidentified fetal cerebral astrocyte cell cultures, the UCLA Office for Protection of Research Subject (OPRS) declared IRB exemption status for pre-shelved fetal brain specimen donation per HS-7 exemption amendment. Deidentified human fetal brain tissue from voluntarily aborted pregnancies was donated adhering to stringent ethical regulations. Tissue was preshelved at the UCLA Translational Pathology Core Laboratory (Sara Dry, Chair of Pathology and Laboratory Medicine). Consent was established through UCLA Dept of OB-GYN (Angela Chen, Division Chief of Family Planning). Signed informed consent occurred after decision on elective termination, complying with US Public Law 103-43 and NIH policy, ensuring no incentive (The Public Health Service Act, 42 U.S.C. § 289g-2).

### CONSENT FOR PUBLICATION

This manuscript contains no individual personal information, so individual consent for publication is not applicable. All authors have read and approved the manuscript in its present version.

### AVAILABILITY OF DATA AND MATERIAL

The datasets supporting the conclusions of this article are included within the article and its additional files with file names listed under Additional material. The CSF GFAP and BDP longitudinal quantification dataset of 24 TBI patients is freely accessible through the Open Data Commons for Traumatic Brain Injury repository (ODC-TBI), under the Wanner lab space with data dictionary, the unique persistent identifier, the hyperlink to this dataset is provided here: https://odc-tbi.org/reviewers/55e8d5e33ad69bd411313791f85sfn25

**Wanner IB,** Halford J, Chen Y, Lopez J, Shen S, Van Meter TE, Shaw G, Vespa P, Loo R and Loo J.: GFAP Degradation in TBI: Linking Modified Products to Astrocyte Pathology and Patient Outcomes 2025 BioRXiv Preprint: https://doi.org/10.1101/2025.08.01.668181

Mass spectrometry raw data of GFAP fragments’ peptide sequence PD files and label-free peptide/ion quantification of four TBI patients (six CSF samples) are publicly accessible at the Center for Computational Mass Spectrometry (CCMS) ProteoSAFe (Proteomics Software Environment) web server as a MassIVE (MS Interactive Virtual Environment) file folder provided here: https://massive.ucsd.edu/ProteoSAFe/dataset.jsp?task=460f951a666a4b599df2bfa37fa33fbd.

## Supporting information

GFAP-degradome-Supplemental Figures and Tables

## Figures and Supplements

This manuscript includes a PowerPoint file with eight figures called ‘*Figures-GFAP-trauma-Wanner’*. A supplement is included, which contains 14 Additional Figures (labeled Figures S1-S14) and eight Tables S1-S8 that are compiled into one pdf file called ‘*Supplement-GFAP-trauma-Wanner’*. Further, complete and uncropped immunoblot images of the figures are provided in *‘GFAP-uncropped blots-Supplementary-Wanner’*.

## COMPETING INTERESTS

**IBW** and **JAL** are inventors of the following patents licensed to BRAINBox Solutions Inc., where **TVM** serves as Chief Scientific Officer (CSO). **GS** is CEO and founder of EnCor Biotechnology Inc., which distributes several monoclonal GFAP antibodies used in this study.

**US Patent Issue No.: US 10,557,859 B2** original patent; Inventors **IBW** and **JAL**, Issue date: 2/11/2020; 25 claims prior publication 12/20/2018: “Astrocyte traumatome and neurotrauma biomarkers”.

Applicant Regents of the University of California, Oakland, CA. Licensee: BRAINBox Solutions Inc.

**US Issue No.: US 11,249,094 B2** Additional 13 claims in divisional patent 13 Claims, Filed 12/116/2019; Issue date: 2/15/2022; Inventors: **IBW** and **JAL**, UCLA “Astrocyte traumatome and neurotrauma biomarkers”. Assignee UCLA; Licensee: Brain Box Solutions Inc.

## FUNDING

Dana Foundation: Clinical Neuroscience “First in man” award # 20182601: “Clinical study and trauma culture model experiments”

Brain Injury Research Center (BIRC), State of California: “Clinical study and trauma culture model experiments”

National Institute of Health, Institute of Neuronal Disorders and Stroke, NIH/NINDS, R01 NS052831 “Biochemical markers of severe traumatic brain injury” NIH/NINDS Small Business Innovation Research, SBIR grant # 1R43NS106972-01 “Validation and GFAP proteoform characterization for assay use”

Sponsored Research Agreement, Abbott Diagnostics: “GFAP fragment analyses”

Sponsored Research Agreement, BRAINBox Solutions Inc., Richmond VA, USA: “Validation studies and patent licensing”

Semel Institute IDDRC and Brain Injury Research Center, BIRC, UCLA, Los Angeles, CA, USA. Facilities supplies for cell culture and microscopy work.

## AUTHORS’ CONTRIBUTIONS

**IBW** conceptualized and designed the study, wrote and revised the manuscript, curated data and conducted formal analysis, designed and prepared all visualizations, acquired funding, coordinated research activities, supervised laboratory assistants and students and conducted cell culture and live cell imaging studies. **JH** and **JL** conducted sample preparation and immunological experiments, assembled raw data and illustrations, curated and analyzed data. **JL, SS** and **YC** analyzed MS spectra. **YC** and **SS** conducted MS analyses. **HZ** provided the thumbnail infographic. **CS** analyzed data and provided visualizations. **ROL** and **JAL** provided MS data analysis guidance and **JAL** was involved in the study conceptualization. **TVM** and **GS** provided antibodies, peptides and recombinant proteins, assisted in writing and editing the manuscript and offered biochemical guidance. **GS** also conducted antibody binding and chemical cleavage studies and wrote the structural part of the discussion. **BME** provided data interpretation and manuscript revision. **JG** provided guidance on data curation and statistical analysis approaches. **SR** and **PMV** had oversight over clinical aspects of the study including TBI patient enrollment, IRB approval and sample provision. Further, **PMV** provided guidance, manuscript organization and contributed to written content and clinical interpretation of the findings. Resources included **IDDRC** laboratory infrastructure including cell culture, centrifuges, gel imagers. The **MIC** provided mass spectrometry instruments and proteomics facilities for the study. ***All authors*** *read, reviewed, edited and approved of the final manuscript*.

## ACKNOWLEDGEMENTS

The authors acknowledge with appreciation the receipt of mouse antibody CTGF-1221 to citrullinated GFAP (RRID: AB_3678889) that was kindly provided by Dr. Akihito Ishigami, Tokyo Metropolitan Institute of Gerontology, Tokyo, Japan. Special thanks goes to Dr. Sarah Dry and Ko Kiehle Translational Pathology Core lab, Ronald Reagan UCLA Medical Center, and Dr. Angela Chen, Family Planning & UCLA OBGYN Ronald Reagan Medical Center, for consenting and procuring voluntary donated fetal remains as pre-shelved specimen for cerebral tissues. Greatly appreciates for their grateful for input from are Laboratory assistant Kiohei Itamura, and UCLA Honors students Michelle Gong, Raghu Padmanabhuni, Aryana Sargsyan, Ana Karimi Bidhendi, Emerald Wong, Mhina Cham, Aryana Sargsyan, Kunal Ranat; David Sanville, and for the valuable critical care management assistance provided by Courtney Real and Jesus Ruiz-Tejada and the help from clinical fellow Azim Laiwalla.

## Thumbnail: Astrocyte traumatic injury states with distinct GFAP proteoforms and fates after TBI

The infographic depicts four distinct coexisting and progressive injury states of human astrocytes after trauma, each defined by distinct astrocyte morphologies, GFAP proteoforms, and distributions. Key pathways of GFAP post-translational modification (PTM), degradation and aggregation are linked to astrocyte dysfunction and underpin fluid biomarker signatures. The illustrated stages connect astrocytopathy with GFAP biomarker profiles to aid interpretation in TBI patient monitoring. ***(1) Uninjured, healthy astrocyte:*** Intact plasma membrane and fully assembled GFAP filaments. ***(2) Mechanoporated, wounded astrocyte:*** Mechanical forces cause membrane poration and process beading (clasmatodendrosis), with calcium influx and filament disassembly into monomers (black) and fragments. Proteolytic is initiated by calpain cleavage into large fragments (gray), followed by caspases and other proteases generating coil1 and 2 co-products: coil1-BDPs are readily released into fluids (red), while coil2 products form PTM-decorated aggregates (purple). ***(3) Degenerating, dystrophic astrocytes:*** Swollen, oncotic morphology with vacuoles, membrane blebbing, and severely damaged processes. Degenerating astrocytes contain accumulated PTM-modified GFAP granules marked by citrullination (green) and acetylation (brown) documenting assembly-defective GFAP aggregation. ***(4) Dying, necrotic, astrocytes:*** Loss of cell integrity with membrane rupture and lysis, release of modified coil2 products as markers of irreversible cell death.

---

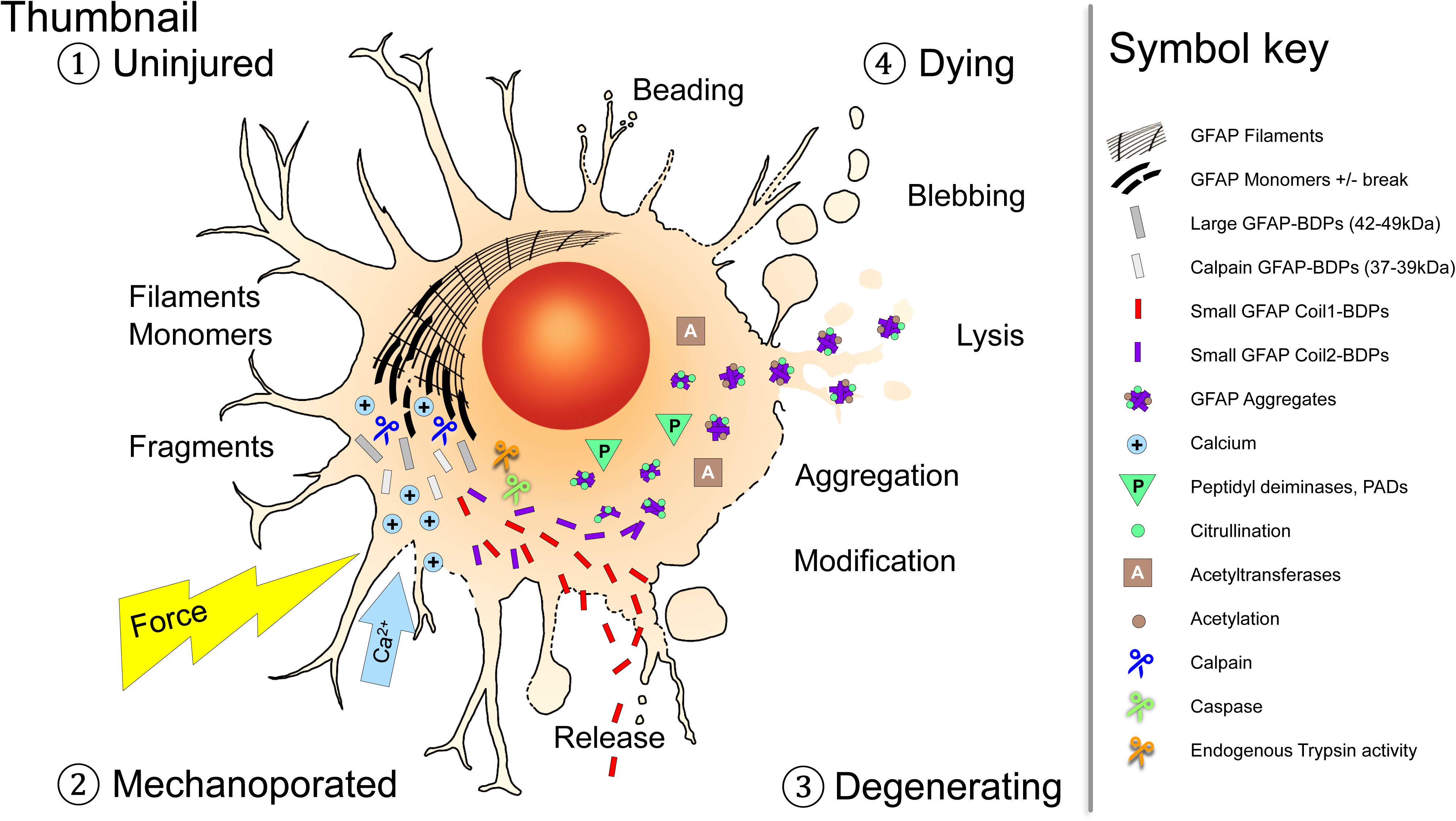

