## Supplementary material for "GFAP Degradation in TBI: Linking Novel Modified Products to Astrocyte Pathology and Patient Outcome": GFAP-degradome-Supplemental Figures and Tables

### Table S1

| Demographic | Intake Region | of Patients | |
|---|---|---|---|
| California | Los Angeles (UCLA) | 21 | |
| Florida | Gainesville Univ. of Florida | 5 | |
| Parameter | | Median | *SD* |
| **Age** | | 40 | *19* |
| **Severity** | GCS on Admission Day | 6 | *3.4* |
| **Outcome** | GOS at Discharge | 3 | *1* |
| | GOSE at 6 months | 4 | *2.1* |
| Parameter | | Number | Percent |
| **Race** | White | 15 | 57.7 |
| | Hispanic | 8 | 30.8 |
| | Black | 1 | 3.8 |
| | Asian | 1 | 3.8 |
| | Unknown | 2 | 7.7 |
| **Sex** | Male | 22 | 84.6 |
| | Female | 4 | 15.4 |
| **Pre-medical History** | Yes | 9 | 34.6 |
| | No | 12 | 46.2 |
| | Unknown | 5 | 19.2 |
| **Injury Cause** | Motor vehicle crash | 3 | 11.5 |
| | Motor cycle accident | 3 | 11.5 |
| | Fall | 9 | 34.6 |
| | Pedestrian versus auto | 8 | 30.8 |
| | Unknown (GNV) | 3 | 11.5 |
| **Injury locations on CAT scan** | Known | **24** | **92.3** |
| **Intracerebral** | Contusion | 12 | 50.0 |
| | Hemorrhage/hematoma | 14 | 58.3 |
| | Edema | 7 | 29.2 |
| | Midline shift | 10 | 41.7 |
| | Diffuse axonal injury | 5 | 20.8 |
| **Leptomeningeal** | Subdural blood | 8 | 33.3 |
| *hemorrhage, hematoma | Subarachnoideal blood* | 6 | 25.0 |
| ** including blood* | Extradural /epidural fluid** | 4 | 16.7 |
| **Intraventricular** | Ventricular blood | 11 | 45.8 |
| **Bone fractures** | | 6 | 25.0 |
| **Surgery** | Ventriculostomy | 14 | 58.3 |
| | Craniotomy | 5 | 20.8 |
| | Craniectomy | 10 | 41.7 |
| | Hemicraniectomy | 5 | 20.8 |

### Table S2

| Samples, Analysis | # Samples | # Patients | Samples / patient |
|---|---|---|---|
| Cerebrospinal fluid | 112 | 26 | 1 to 18 |
| Mass spectrometry (MS) | 6 | 5 | 1 to 3 |
| Immunoblot & densitometry | 108 | 24 | 3 to 18 |
| CSF-draw concurrent temperature | 69 | 23 | 1 to 9 |
| GFAP bandset trajectory analysis | 102 | 23 | 3 to 18 |
| GFAP bandset outcome prediction | 95 | 21 | 3 to 18 |
| Serum (for MS) | 2 (TBI), 1 (Control) | 2 TBI, 1 Ctrl | 1 |

**Table S1: TBI patient demographics and clinical parameters.**
Neuro-Intensive care (ICU, n=26) TBI patients from two centers. Age, admission Glasgow Coma scale (GCS) score and outcome score on hospital discharge (GOS) and at six months (GOSE) as well as pre-medical history and injury cause. CT scan report is summarized based on cerebral locations and surgical interventions.

**Table S2: TBI patient samples collected and analyzed.**
The number of samples collected per patient followed by listings of samples used for mass spectrometry (MS) and for immunoblot densitometric GFAP-BDP quantification and samples used for GFAP and GFAP-BDP profile outcome prognosis.

### Table S3

| Antibody name/clone | RRID | Species | Epitope | Source | Immunoblot (IB) dilution (ECL type) | Immunoprecipitation | IF dilution (with secondary antibodies, Jackson Immunoresearch, minimal cross-species reactivity) |
|---|---|---|---|---|---|---|---|
| Rabbit polyclonal anti-GFAP | RRID:AB_10013382 | rabbit | 10 epitopes defined | Agilent, DAKO | 1:7,000 to 1:150,000 (Pico) | capture antibody 1:1 (weight) to 1:12 (volumes) crosslinked to Protein G magnetic beads | N/A |
| Mouse anti-GFAP 1A-244 | RRID: AB_3711244 | mouse IgG | coil1 20kDa GFAP fragment | Abbott Diagnostics | 1:2,000 (serum ) 1:20,000 (CSF) | detection antibody (see IB) | N/A |
| MCA-3E10 | RRID:AB_10013382 | mouse IgG | raised to aa71-217 here determined: coil 1B motif: aa 196-210 epitope with binding motif : ESLEEEIRFLRKIHEEEVRELQEQLAR; | EnCor Biotechnology Inc. | 1:5,000 to 1:15,000 (Pico) | detection antibody (see IB) | N/A |
| CPCA anti-GFAP | RRID:AB_2109953 | chicken | polyclonal | EnCor Biotechnology Inc. | 1:20,000 (CSF) | detection antibody (see IB) | 1:700 (with donkey anti chicken Cy5 1:80) |
| MCA-5C10 | RRID:AB_2572311 | mouse IgG | between coil1B and 3B covering coil2A: epitope with motif: RTQYEAMASSNMHEAEEWYRSKFAD | EnCor Biotechnology Inc. | 1:1,000 to 1:2,000 (Pico) | N/A | N/A |
| Anti-GFAP clone GA5 | RRID:AB_477010 | mouse IgG | c-terminal rod-to-tail region (Chen et al., 2011; 2013) | Sigma | 1:1,000 to 1:2,000 (Pico/Femto) | N/A | 1:260 (with donkey anti mouse IgG-Cy3 1:250) |
| Anti-GFAP Cocktail (4A11, 1B4, 2E1) | RRID:AB_10013382 | 3 mouse IgGs | proprietary | Agilent, BD Pharmingen | 1:1,000 (Pico) | N/A | N/A |
| CTGF-1221 to citrullinated GFAP | RRID: AB_3678889 | mouse IgG | Citrullinated GFAP at R270 &R416 (Ishigami et al., 2015) | Ishigami et al., 2015 | 1:1,000 (Femto) | N/A | N/A |
| Anti-peptidyl citrulline clone F95 | RRID:AB_10013382 | rabbit polyclonal | citrullination, not specific to GFAP | Sigma, EMD Millipore | 1:500 (Femto) | N/A | N/A |
| cit-270-GFAP | RRID: AB_3678888 | rabbit monoclonal | Citrullinated GFAP at R270 | BrainBox Solutions, Inc. | 1:1,000 (Femto) | N/A | 1:180 (with donkey anti-rabbit-Alexa488 1:250) |
| Pan-intermediate filament ab, Anti-IFA | presently not commercial available | mouse hybridoma supernatant | coil2B aa 358-377: ALDIEIATYRKLLEGEENRI | Kouklis et al., 1992; Pruss et al., 1985; ATCC (unavailable) | 1:1,000 | N/A | N/A |
| mouse alpha-II Spectrin (Fodrin, clone AA6) | RRID:AB_2050678 | mouse monoclonal IgG | not provided, detects full-length and caspase and calpain cleavage products. | Enzo Life Science | 1:3,000 to 1:5,000 (Pico) | N/A | N/A |

**Table S3: Antibodies** Listed are GFAP, citrullination and αII-spectrin antibody characteristics with epitopes and binding motifs (if known). Dilutions used for immunoprecipitation and with electrochemiluminescence (ECL)-base immunoblotting, as well as secondary antibody choices for immunofluorescence (IF).

Fig. S1

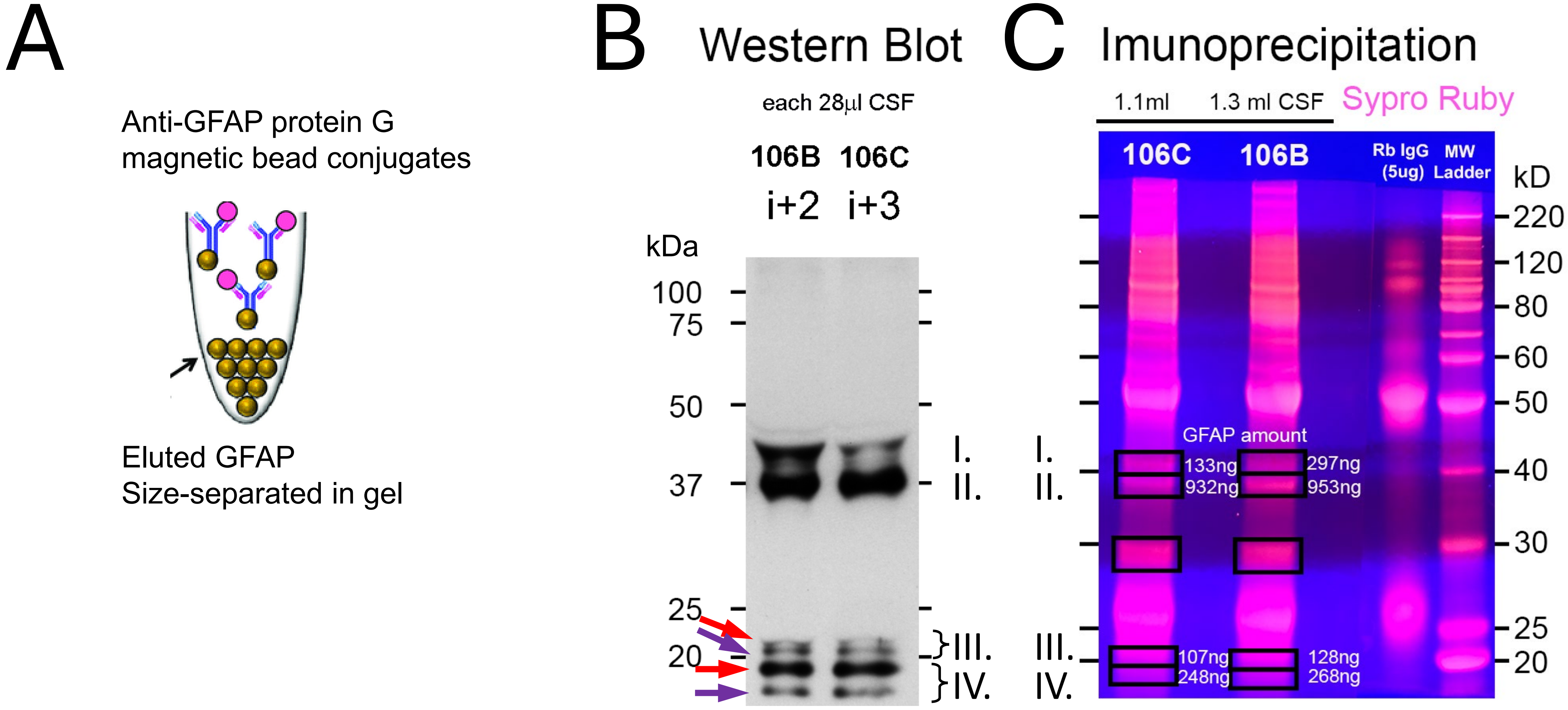


**Fig. S1: GFAP Immunoprecipitation (IP) for LC-MS/MS.** **A)** Drawing depicts protein G magnetic bead-based GFAP immunoprecipitation using a rabbit polyclonal GFAP antibody approach from TBI CSF (see Methods). **B)** GFAP immunoblot (mab 3E10) shows immunoprecipitated GFAP from CSF samples of a TBI patient at two and three PIDs (i+2, i+3). **C)** Sypro Ruby-stained preparative gradient gel of same samples alongside immunoglobulin and molecular weight indicator. Black boxes show 4 gel excised band sets sequenced via LC-MS/MS. Bands I.: 41-42kDa, Bands II. 36-40kDa, Bands III. 21-23kDa and Bands IV. 15-20kDa GFAP.

### Fig. S2

#### Recombinant GFAP , sequence coverage 96%

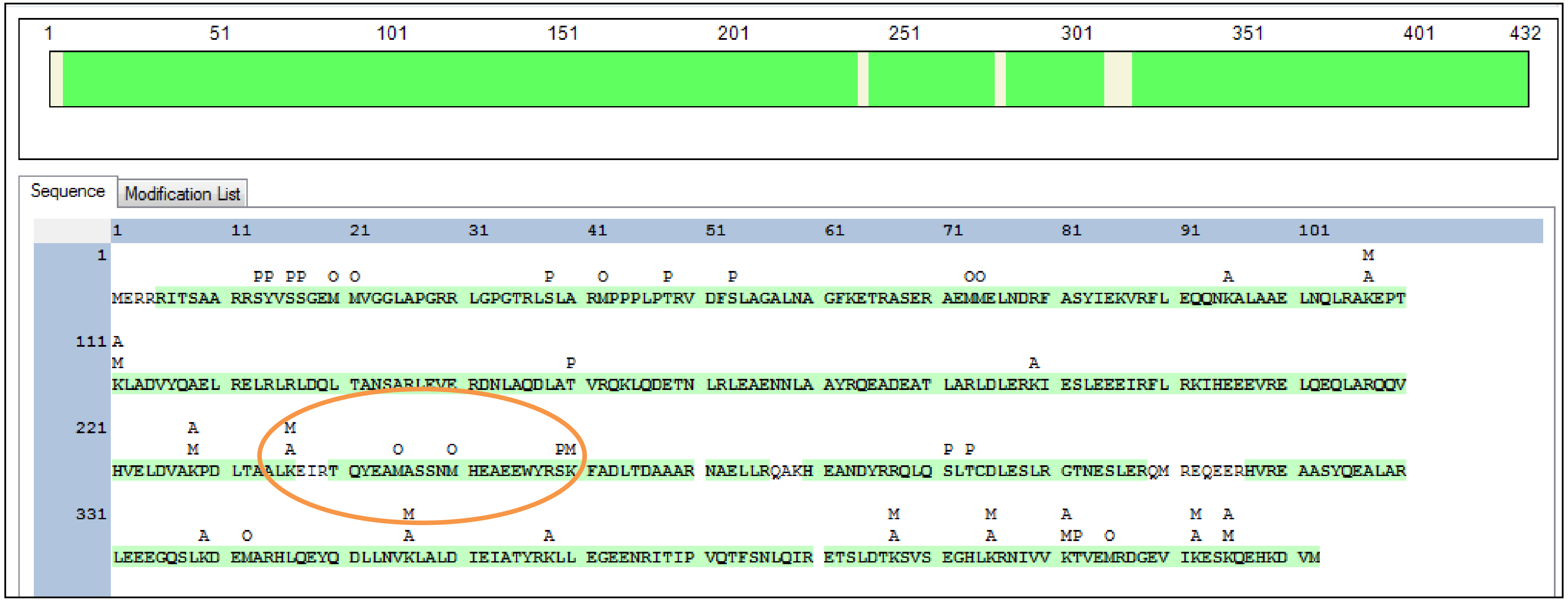


**Fig. S2: Positive control for GFAP sequence coverage.** MS sequence data show full length GFAP sequence with 96% coverage by high confidence peptides (green highlighted peptides, each detected at least twice, typically multiple times). Recombinant human GFAP (500ng) was analyzed using same settings as immunoprecipitated GFAP from TBI patient-derived samples (Methods). Circle: At least two high confidence peptides covered aa 240-260 here that were missing in TBI patient samples.

### Fig. S3

#### GFAP-BDP coil-specific peptide ion peak heights reflect imbalanced abundance

Linear-spaced peptide abundance

Log-spaced peptide abundance

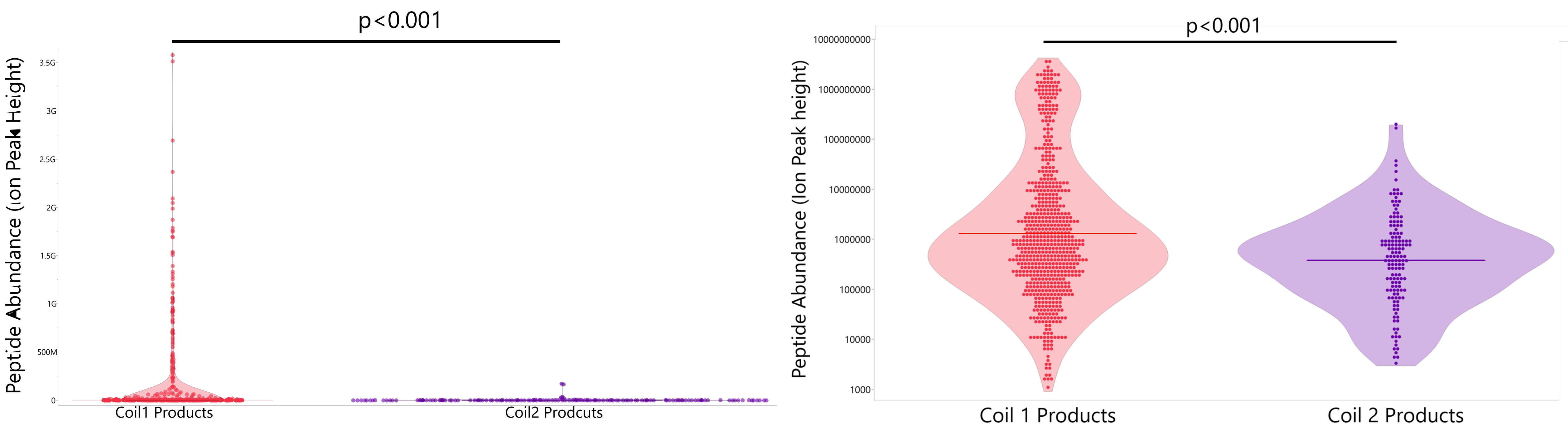


**Fig. S3: Imbalanced of coil1 and coil2 in label-free MS by peptide ion peak abundance in bands of small GFAP-BDPs.** Plotted are 6 TBI CSF samples from 5 TBI patients from 11 MS runs of small GFAP-BDPs from IP cut-out Bands III. and IV. Ion peak height of each recorded peptide sorted by coil1 and coil2 containing peptides document peptide abundance in TBI CSF. Analysis of variance with patient ID as random and fragment coils as fixed effects and undetected peptides excluded (see Fig.1D for peptide frequencies by coils). Ion peak height is graphed using linear y-axis (left) and log-spaced y-axis (right).

Fig. S4

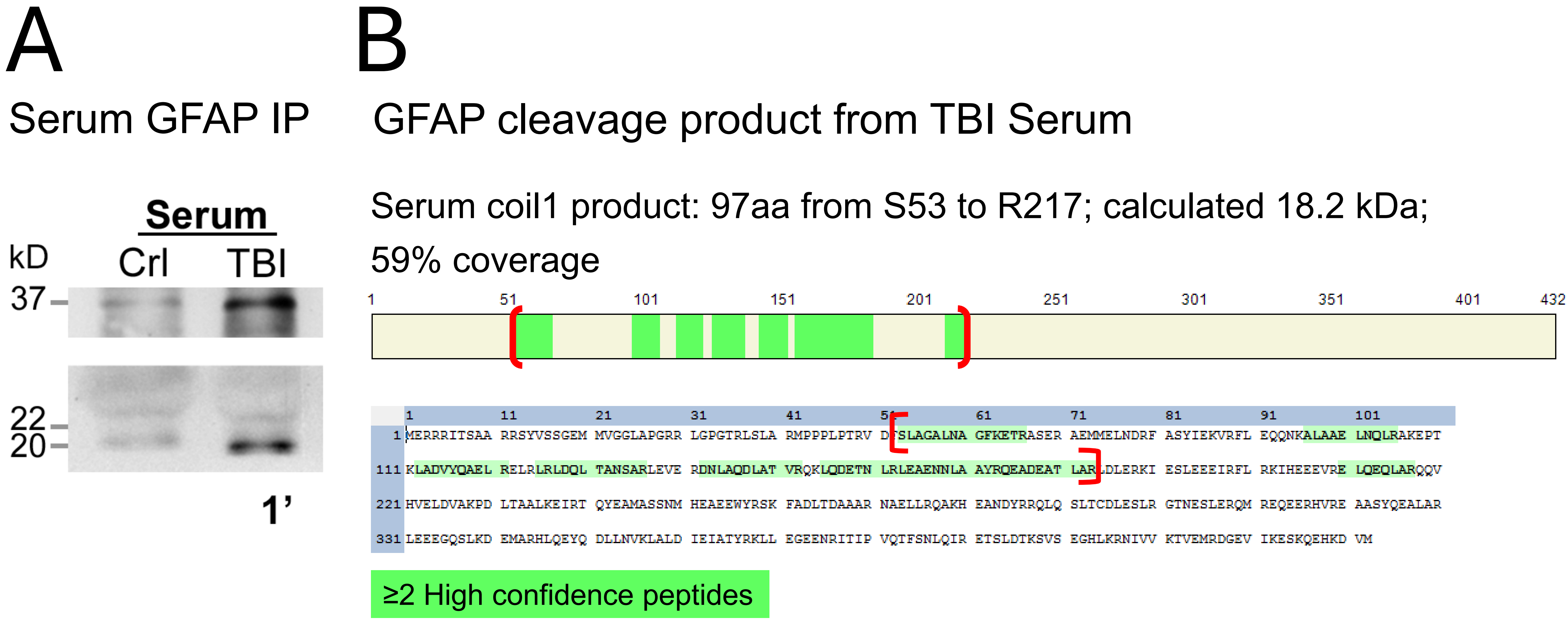


**Fig. S4: TBI Serum-immunoprecipitated GFAP coil1 product. A)** Serum GFAP was immunoprecipitated by a protein G-conjugated rabbit polyclonal anti-GFAP (DAKO) and detected using a mab anti-GFAP (clone 1A244, Abbott; exposure 1min). TBI sera were pooled from i+1and i+2 from a 54-year-old man patient who recovered with moderate disability from a fall; control serum: Precision Med. **B)** MS sequenced high confidence peptide map (green) covering 59% of a 97aa cleavage product from S53 and R217 of a 19-20kDa trypsinized gel slice with estimated mass of 18.2kDa was within coil1 of the GFAP rod. Less immunoprecipitated material than for CSF was run.

### Tables S4

#### A PTMs in COIL 1 products

| Residue with PTM | PTM Validation | Peptides with modification |
|---|---|---|
| “Bands III.” 21-22 kD | | |
| R66(citrullination) | m/z = 130.08534 | 2 of 3 |
| R79(citrullination) | m/z = 130.08534 | 1 of 2 |
| R105(citrullination) | m/z = 130.08505 | 5 of 8 |
| R188(citrullination) | m/z = 130.04985 | 1 of 7 |
| K189(Acetyl) | m/z = 126.09063, m/z = 143.11740 | 1 of 3 |
| K202(Acetyl) | m/z = 126.09151 | 1 of 1 |
| “Bands IV.” 21 kD | | |
| R66(citrullination) | m/z = 130.08563 | 2 of 2 |
| R79(citrullination) | Spectrum mass loss validated | 1 of 1 |
| R88(citrullination) | Likely; and GFAP+PAD2 confirmed* | Possible in 1 of 1 |
| R105(citrullination) | m/z = 130.08517 | 2 of 2 |
| R121(citrullination) | Spectrum mass loss validated | 1 of 7 |
| R136(citrullination) | Likely; and GFAP+PAD2 confirmed* | Possible in 3 of 4 |
| R173(citrullination) | Likely; and GFAP+PAD2 confirmed* | Possible in 1 of 2 |
| R188(citrullination) | m/z = 130.04929 | 1 of 3 |
| R217(citrullination) | Likely; and GFAP+PAD2 confirmed* | Possible in 3 of 7 |
| K63 (Acetyl) | m/z = 126.06519 | 2 of 2 |
| K189(Acetyl) | m/z = 126.09170, m/z = 143.11832 | 2 of 2 |
| K202(Acetyl) | m/z = 126.09063, m/z = 143.11684 | 2 of 2 |

#### B PTMs in COIL 2 products

| Residue with PTM | PTM Validation | Peptides with modification |
|---|---|---|
| “Bands III.” 17.5 kD | | |
| R270(citrullination) | m/z = 130.08548 | 3 of 3 |
| R276(citrullination) | m/z = 130.08548 | 3 of 3 |
| R406(citrullination) | m/z = 130.08659 | 1 of 1 |
| “Bands IV.” 15.3 kD | | |
| R270(citrullination) | m/z = 130.08522 | 3 of 3 |
| R276(citrullination) | m/z = 130.08522 | 3 of 3 |

High confidence peptides; citrullination signature ion(s) present.

Novel, signature ion present GFAP citrullination site

Medium confidence peptides; MS2 ion mass shifts ($b^+$; $y^+$) present.

*High confidence peptides, MS2 ion mass shifts, reported cell-free GFAP deimination (Jin et al., 2013).

High confidence peptides; acetylation signature ion(s) present.

**Table S4: Frequency of coil1 and coil2 PTMs in TBI patients’ CSF.** Listed are peptide residues with Proteome Discoverer-designated and thereafter MS2 spectrum validated post-translational modifications (PTMs) tabulated for **(A)** coil1 and **(B)** coil2 GFAP products. *PTM validation:* via MS2 signature ion mass shifts for arginine (R) citrullination: immonium ion at m/z 130.0975 and/or neutral loss of isocyanic acid at 43.0058 Da; and for lysine (K) acetylation: cyclized immonium ion at m/z 126; (Example spectra: Figure S10). Alternatively, confirmation was by cell-free PAD deamination [65]. Peptides with modification: Number of peptides carrying same modification out of peptides carrying same residue, MS analyses from 6 TBI CSF samples.

### Fig. S5 (A)

# A

MS2 spectrum signature ions for novel GFAP citrullinated R79 in peptide NDRFASYIEK

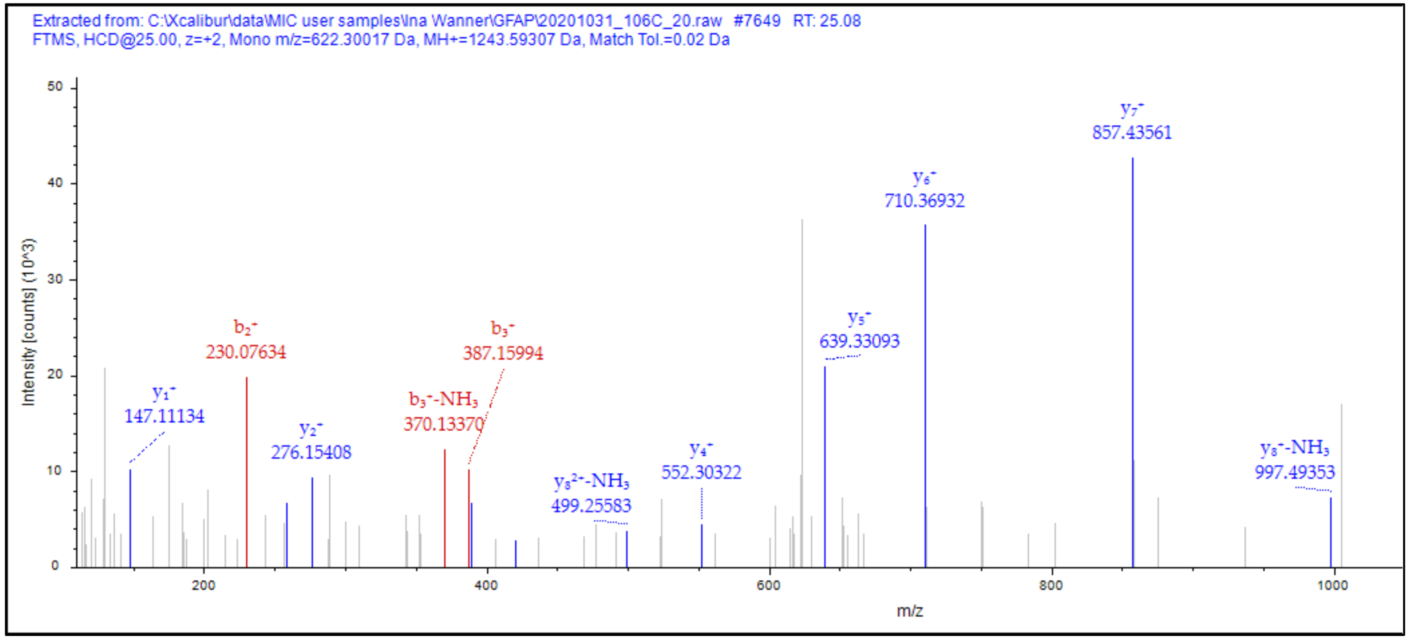


Charge: +2, Monoisotopic m/z: 622.30017Da (-1.16 mmu/-1.87 ppm), MH+: 1243.59307Da, RT: 25.08min

#### Fragment ion series chart

| #1 | $b^+$ | $b^{2+}$ | Seq. | $y^+$ | $y^{2+}$ | #2 |
|---|---|---|---|---|---|---|
| 1 | - | - | N | | | 10 |
| 2 | +3.55 | - | D | - | - | 9 |
| 3 | +6.08 | - | R-citrullination | - | - | 8 |
| 4 | - | - | F | +5.55 | - | 7 |
| 5 | - | - | A | +3.70 | - | 6 |
| 6 | - | - | S | +6.10 | - | 5 |
| 7 | - | - | Y | -0.76 | - | 4 |
| 8 | - | - | I | +7.79 | - | 3 |
| 9 | - | - | E | +4.82 | - | 2 |
| 10 | | | K | +10.00 | - | 1 |

**Figure S5: MS2-spectrum signature-validated GFAP citrullination sites:** Representative examples of three MS spectra from TBI patients' CSF immunoprecipitated GFAP summarized in Table S4 and Figure 4A. *MS2 spectra -higher energy collisional dissociation (HCD) spectra* (left)- show ion intensity/counts over ion sizes in mass/charge, (m/z) of specific monoisotopic ions at the given MS run time plotted are modified ion fragment size (x-axis) and signal intensity (y-axis). Ion mass shifts serve as signature validating citrullinated peptides. Mass shifts are tabulated in *fragment ion series charts* (right) informing on b+ and Y+ charge deviations from regular arginine or lysine identifying that a modification occurred. **A)** Potential novel GFAP citrullination site at arginine position 79 in peptide NDRFASYIEK **B)** Peptide sequence ALAAELNQLRAK with citrullinated arginine at position 105.
**C)** Peptide FADLTDAAARNAELLRQAK with signature ion mass changes identifying citrullinated arginine at positions 270 and 276.

### Fig. S5 (B)

# B

MS2 spectrum signature ions for citrullinated R105 in peptide ALAAELNQLRAK

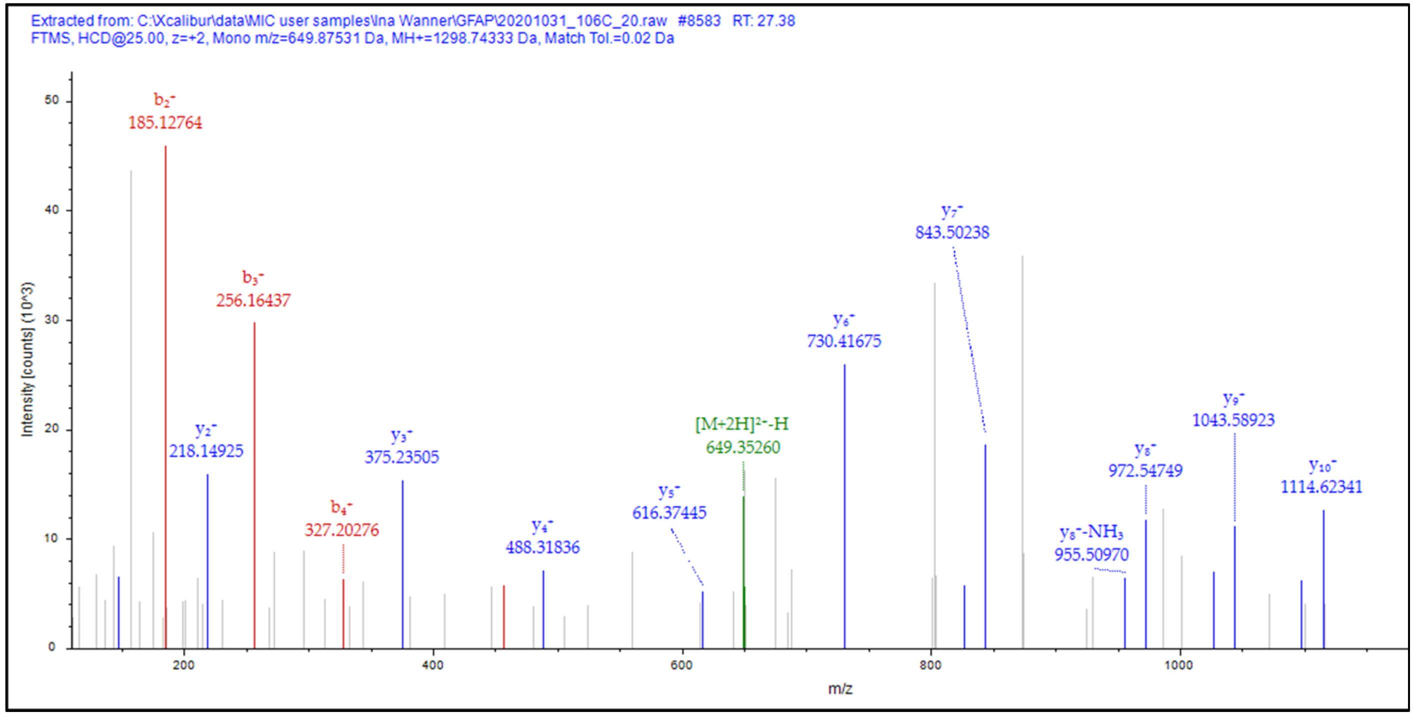


Fragment ion series chart

| #1 | $b^+$ | $b^{2+}$ | Seq. | $y^+$ | $y^{2+}$ | #2 |
|---|---|---|---|---|---|---|
| 1 | - | - | A | | | 12 |
| 2 | +4.47 | - | L | - | - | 11 |
| 3 | +4.75 | - | A | -1.66 | - | 10 |
| 4 | -0.16 | - | A | -4.59 | - | 9 |
| 5 | -7.88 | - | E | -0.18 | - | 8 |
| 6 | - | - | L | +2.77 | - | 7 |
| 7 | - | - | N | +5.34 | - | 6 |
| 8 | - | - | Q | +5.30 | - | 5 |
| 9 | - | - | L | +1.59 | - | 4 |
| 10 | - | - | R-citrullination | +0.05 | - | 3 |
| 11 | - | - | A | +3.12 | - | 2 |
| 12 | | | K | +5.65 | - | 1 |

Charge: +2, Monoisotopic m/z: 649.87531Da (+0.29mmu/+0.45ppm), MH+: 1298.74333 Da, RT: 27.38min,
Identified with: Sequest HT (v1.3); XCorr:3.03,
Fragment match tolerance used for search: 0.02Da
Fragments used for search: b; b-$H_2O$; b-$NH_3$; y; y-$H_2O$; y-$NH_3$

### Fig. S5 (C)

# C

MS2 spectrum signature ion validation of citrullinated R270 and R276 in the peptide: FADLTDAAARNAELLRQAK

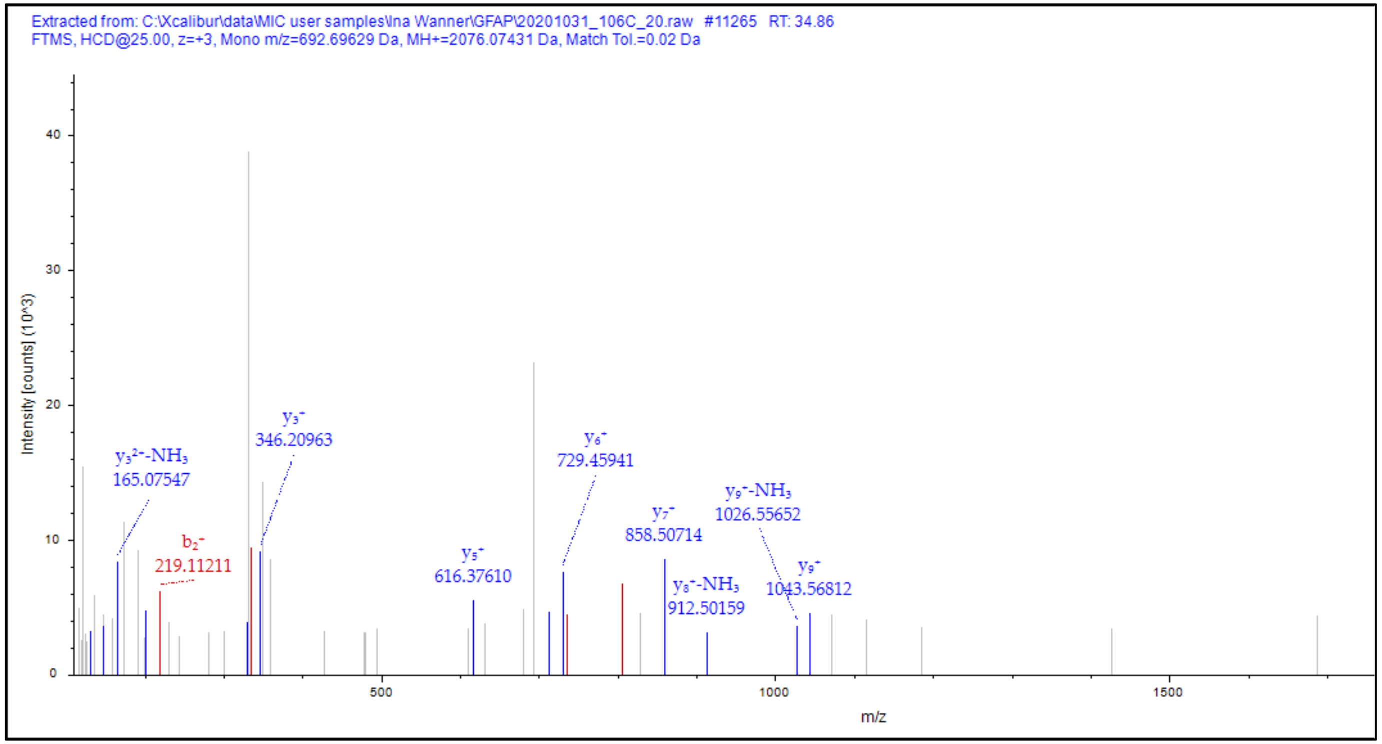


#### Fragment ion series chart

| #1 | $b^+$ | $b^{2+}$ | $b^{3+}$ | Seq. | $y^+$ | $y^{2+}$ | $y^{3+}$ | #2 |
|---|---|---|---|---|---|---|---|---|
| 1 | - | - | - | F | | | | 19 |
| 2 | +3.22 | - | - | A | - | - | - | 18 |
| 3 | -2.37 | - | - | D | - | - | - | 17 |
| 4 | - | - | - | L | - | - | - | 16 |
| 5 | - | - | - | T | - | - | - | 15 |
| 6 | - | - | - | D | - | - | - | 14 |
| 7 | +6.18 | - | - | A | - | - | - | 13 |
| 8 | -3.90 | - | - | A | - | - | - | 12 |
| 9 | - | - | - | A | - | - | - | 11 |
| 10 | - | - | - | R-citrullination | - | - | - | 10 |
| 11 | - | - | - | N | +15.64 | - | - | 9 |
| 12 | - | - | - | A | - | - | - | 8 |
| 13 | - | - | - | E | -3.21 | - | - | 7 |
| 14 | - | - | - | L | +3.26 | - | - | 6 |
| 15 | - | - | - | L | +2.62 | - | - | 5 |
| 16 | - | - | - | R-citrullination | - | - | - | 4 |
| 17 | - | - | - | Q | -3.23 | - | - | 3 |
| 18 | - | - | - | A | - | - | - | 2 |
| 19 | | | | K | +5.11 | - | - | 1 |

Charge: +3, Monoisotopic m/z: 692.69629Da (+0.73 mmu/+1.06 ppm), MH+: 2076.07431 Da, RT: 34.86min

### Fig. S6

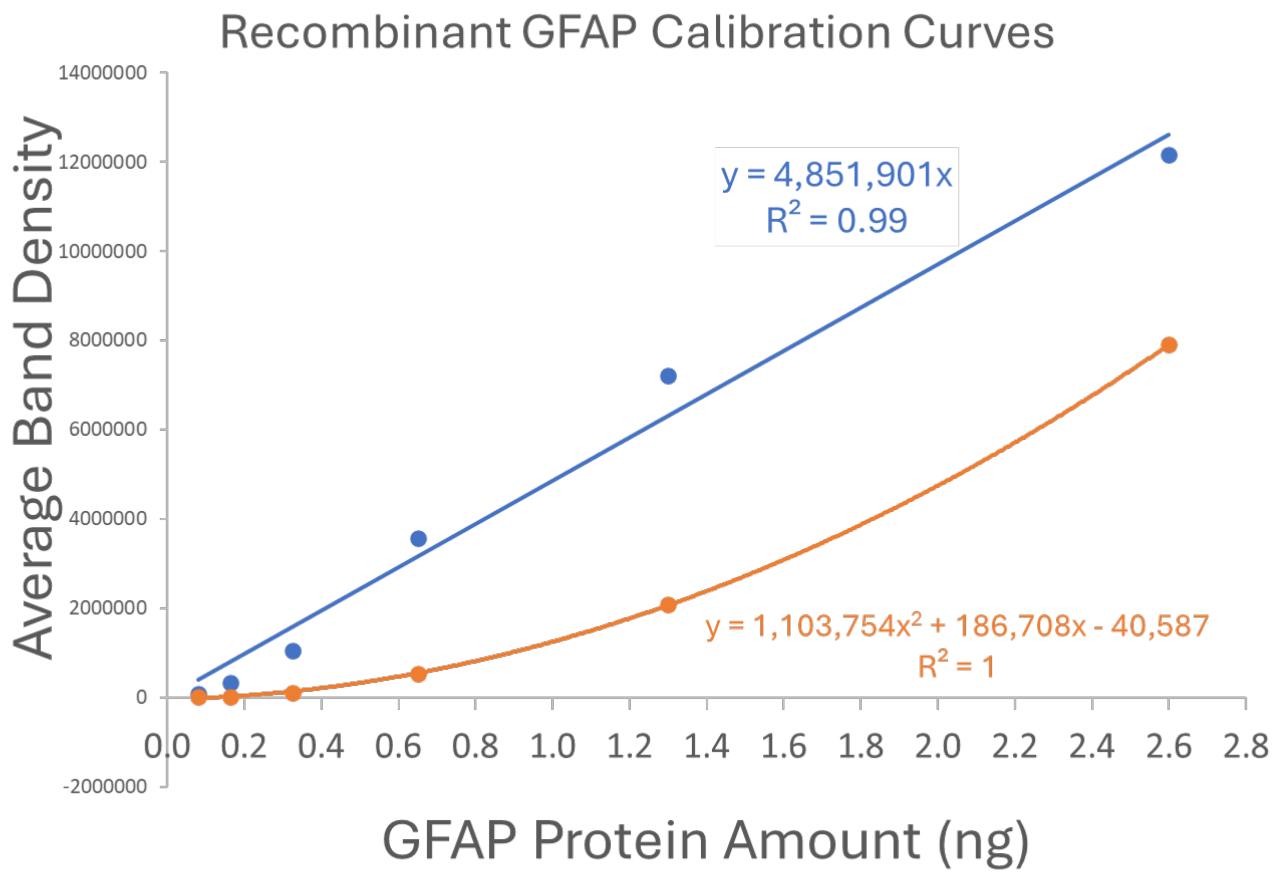


**Figure S6: Recombinant GFAP and GFAP-BDP calibration curves for immunoblot densitometry.** Two recombinant proteins were used to generate two calibration curves. A human recombinant GFAP (EnCor) was used for measuring 50kDa and large GFAP-BDPs (blue, linear formula). For quantification of small GFAP-BDPs a human 25kDa GFAP fragment was made that contained coil1 and a His-tag (EnCor, orange curve, estimated by Desmos.com). Dilutions in duplicates were: 0.08ng; 0.16; 0.33; 0.65; 1.3 and 2.6ng of both calibrants). Subsaturated scaled densitometry of band densities is illustrated in Table S4.

### Table S5

**Original Densities**

| GFAP band Standards | GFAP Amount/lane | <1'' | 2'' | 5'' | 10'' | 20'' | 1' | 5' | 20' |
|---|---|---|---|---|---|---|---|---|---|
| | Exposure | <1'' | 2'' | 5'' | 10'' | 20'' | 1' | 5' | 20' |
| 25 kD | 0.08 ng | | | | | 44372 | 85643 | 321822 | |
| 25 kD | 0.08 ng | | | | | | | 20851 | 89178 |
| 50 kD | 0.16 ng | | 14476 | 36665 | 95416 | 201435 | | | |
| 25 kD | 0.16 ng | | | | | | | 63406 | 184339 |
| 50 kD | 0.33 ng | 33334 | 68795 | 131187 | 218628 | | | | |
| 25 kD | 0.33 ng | | | | 24518 | 73696 | 116086 | 252812 | |
| 50 kD | 0.65 ng | 122632 | 189899 | | | | | | |
| 25 kD | 0.65 ng | | 34173 | 81892 | 159103 | 261980 | | | |
| 50 kD | 1.3 ng | 226501 | | | | | | | |
| 25 kD | 1.3 ng | 80817 | 159068 | 242295 | | | | | |
| 50 kD | 2.6 ng | 382538 | | | | | | | |
| 25 kD | 2.6 ng | 306376 | | | | | | | |

Used densities within range*

**Scalings between exposures**

| GFAP band Standards | GFAP Amount/lane | ud to 2'' | 2'' to 5'' | 5'' to 10'' | 10'' to 20'' | 20'' to 1' | 1' to 5' | 5' to 20' |
|---|---|---|---|---|---|---|---|---|
| 25 kD | 0.08 ng | | | | | 1.93 | 3.76 | |
| 25 kD | 0.08 ng | | | | | | | 4.28 |
| 50 kD | 0.16 ng | | 2.53 | 2.60 | 2.11 | | | |
| 25 kD | 0.16 ng | | | | | | | 2.91 |
| 50 kD | 0.33 ng | 2.06 | 1.91 | 1.67 | | | | |
| 25 kD | 0.33 ng | | | | 3.01 | 1.58 | 2.18 | |
| 50 kD | 0.65 ng | 1.55 | | | | | | |
| 25 kD | 0.65 ng | | 2.40 | 1.94 | 1.65 | | | |
| 50 kD | 1.3 ng | | | | | | | |
| 25 kD | 1.3 ng | 1.97 | 1.52 | | | | | |
| 50 kD | 2.6 ng | | | | | | | |
| 25 kD | 2.6 ng | | | | | | | |

**Scaled Densities to 1'**

| GFAP band Standards | GFAP Amount/lane | <1'' | 2'' | 5'' | 10'' | 20'' | 1' | 5' | 20' | Average scaled all to 1' |
|---|---|---|---|---|---|---|---|---|---|---|
| 25 kD | 0.08 ng | | | | | 77769 | 85643 | 108439 | | 90617 |
| 25 kD | 0.08 ng | | | | | | | 7026 | 8365 | 7696 |
| 50 kD | 0.16 ng | | 247516 | 299980 | 377025 | 353047 | | | | 319392 |
| 25 kD | 0.16 ng | | | | | | | 21365 | 17292 | 19328 |
| 50 kD | 0.33 ng | 1060229 | 1176280 | 1073327 | 863882 | | | | | 1043430 |
| 25 kD | 0.33 ng | | | | 96880 | 129164 | 116086 | 85186 | | 106829 |
| 50 kD | 0.65 ng | 3900463 | 3246957 | | | | | | | 3573710 |
| 25 kD | 0.65 ng | | 584301 | 670012 | 628676 | 459161 | | | | 585538 |
| 50 kD | 1.3 ng | 7204145 | | | | | | | | 7204145 |
| 25 kD | 1.3 ng | 2570485 | 2719798 | 1982375 | | | | | | 2424219 |
| 50 kD | 2.6 ng | 12167095 | | | | | | | | 12167095 |
| 25 kD | 2.6 ng | 9744669 | | | | | | | | 9744669 |

Original calibrant density range: > 26x

Standardize all exposure densities to 1 minute

Final calibrant density range: > 1732x

Average one-step all from shorter to next longer exposure, all scalings multiplied

| Exposure Transitions | <1'' to 2'' | 2'' to 5'' | 5'' to 10'' | 10'' to 20'' | 20'' to 1' | 1' to 5' | 5' to 20' |
|---|---|---|---|---|---|---|---|
| *Scaling Factors* | *1.86* | *2.09* | *2.07* | *2.25* | *1.75* | *2.97* | *3.59* |

*above threshold below saturation

All standardized to 1min

| **Exposure Transitions** | <1'' to 1' | 2'' to 1' | 5'' to 1' | 10'' to 1' | 20'' to 1' | 5' to 1' | 20' to 1' |
|---|---|---|---|---|---|---|---|
| ***Scaling Factors*** | ***31.81*** | ***17.10*** | ***8.18*** | ***3.95*** | ***1.75*** | ***0.34*** | ***0.09*** |
| | *One-step scalings multiplied to 1'* | | | | | *scalings divided to 1'* | |

**Table S5: Scaled Densitometry approach broadened the dynamic range of subsaturated GFAP measurements.** Dynamic range was expanded over 66-fold by scaling different chemiluminescence exposures and standardizing to 1min exposures (scaled densities on right), while all measures were within range (above-detection threshold and below saturation, original densities, left). Scaling ratios between exposure pairs are averaged (center) and scaled to 1min by multiplying transitions of shorter than 1min exposure pairs; and by dividing transitions of longer than 1min exposure pairs (bottom). Established approach, shown for recombinant 50kDa and 25kDa GFAP fragment; used for all GFAP band set measurements of all TBI CSF samples typically 10-13 exposures per serial TBI CSF blot were used, 8 exposures for calibrants (shown here).

### Tables S6

#### Immunoblot densitometry sensitivity and precision

## A

##### GFAP Detection Limits

| Biomarker | Concentration in CSF | | Amount detected on blots | | Dynamic Range |
|---|---|---|---|---|---|
| | LLOD (ng/ml) | ULoQ (ng/ml) | LLOD (ng) | ULoQ (ng) | |
| GFAP (37-50kDa) | 0.027 | 767 | 0.00075 | 21.5 | 28537 |
| GFAP(<37kDa) | 0.0355 | 204 | 0.00065 | 5.7 | 7258 |

## B

##### Inter-experimental technical replicate CV

| Antibody | Sample ID | Band Size | Value 1 (ng) | Value 2 (ng) | Mean (ng) | SD (ng) | CV (%) |
|---|---|---|---|---|---|---|---|
| Rabbit polyclonal anti GFAP (Dako) | 118 i+2 | 50 kD | 0.211 | 0.13 | 0.1705 | 0.057 | 33.6 |
| | | 42-48 kD | 0.127 | 0.118 | 0.1225 | 0.006 | 5.2 |
| | | 37 kD | 0.275 | 0.245 | 0.26 | 0.021 | 8.2 |
| | 118 i+3 | 50 kD | 0.196 | 0.109 | 0.1525 | 0.062 | 40.3 |
| | | 42-48 kD | 0.109 | 0.063 | 0.086 | 0.033 | 37.8 |
| | | 37 kD | 0.298 | 0.162 | 0.23 | 0.096 | 41.8 |
| Average CV between samples (by bandsets) | | 50 kD | | | | | 37.0 |
| | | 42-48 kD | | | | | 21.5 |
| | | 37 kD | | | | | 25.0 |
| Average CV across bands | | | | | | | 27.82 |

**Tables S6: Sensitivity and precision of immunoblot densitometry A) GFAP Detection limits:** The lower limit of detection (LLOD) after background correction and upper limit of quantification (ULoQ) were determined of non-saturated exposures. These were computed separately for full-lengths (50kDa) and large GFAP-BDPs (>/=37kDa) and for small GFAP-BDPs (<37kDa) that were calibrated separately (curve Fig. S6). Limits are given both as concentration in CSF (blue) and as GFAP amount within the 28µl CSF volume loaded (black).

**B) Coefficients of Variation (CVs):** The average between-experiment CVs (SD/mean*100) from technical replicates was 27.8% (n=6 pairs of multiple scaled measurements, specifically 37% for full-length GFAP, 21.5% for fragment bands and 25% for the 37kDa band).

Fig. S7 Distribution and normality of log-transformed GFAP measurements

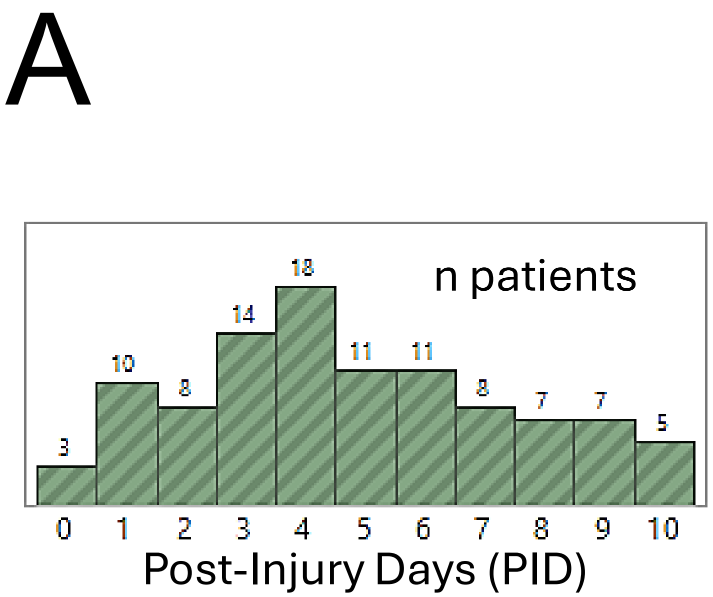


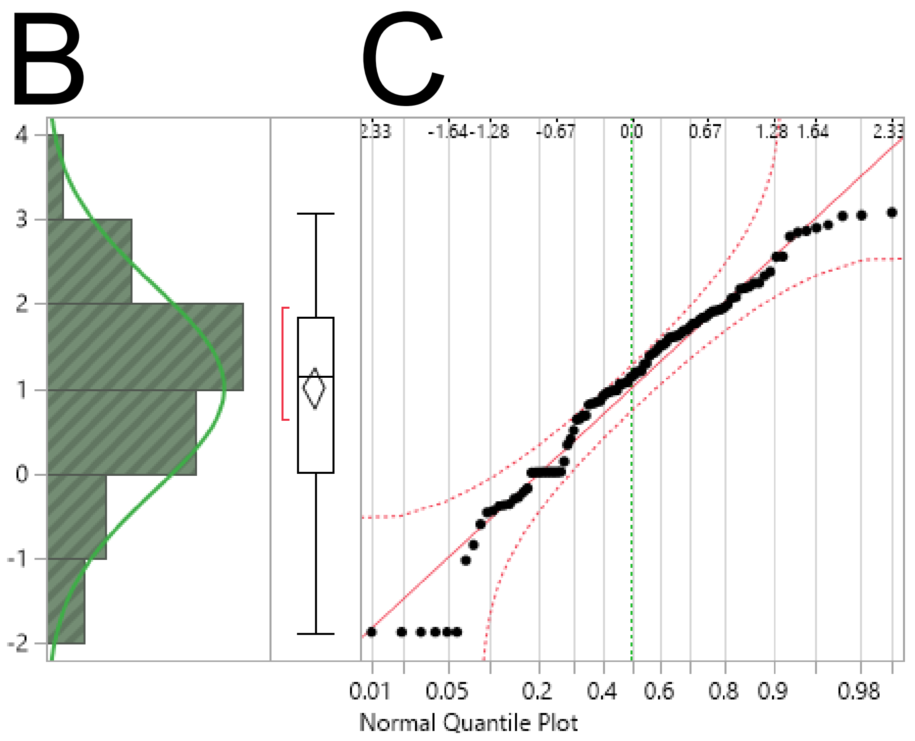


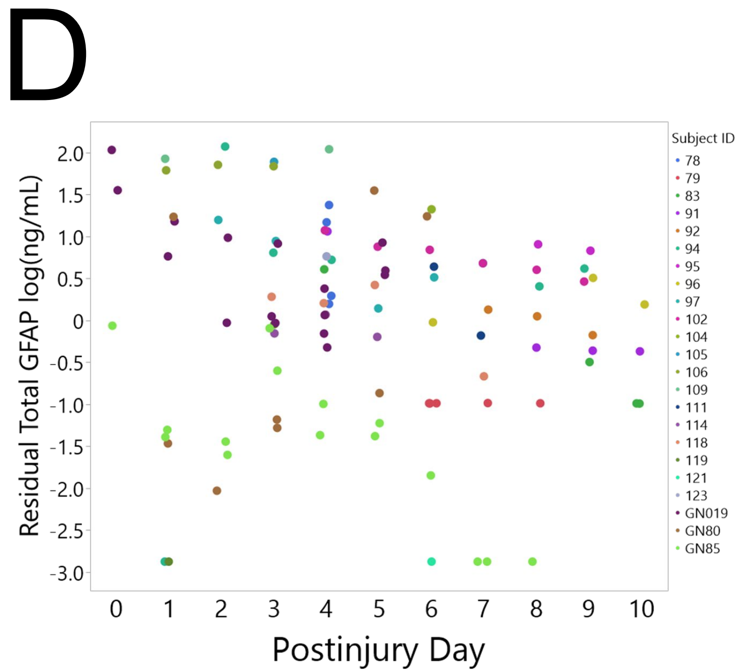


**Fig. S7: Total GFAP TBI CSF data distribution, quantiles and residuals.** **A)** Number of TBI patients' CSF samples collected on each day from injury day (0) to 10PIDs. **B)** Vertical histogram shows frequency distribution of total GFAP log(ng/ml) concentrations binned by log (ng/ml) with fitted normal distribution (green line) that had a Shapiro-Wilk W value of 0.96. Boxplot with densest half (red bracket), median (center line), 25$^{th}$/75$^{th}$ percentiles (box edges) and whiskers extending 1.5x the interquartile range (IQR) of total GFAP being 1.84 log(ng/ml) with outliers beyond. **C)** Normal quantile plot shows quantiles mostly follow normal distribution (red line) and remain within 95% confidence intervals (dotted red lines) documenting acceptable normality. **D)** Residuals of observed minus predicted total GFAP log(ng/ml) are plotted for all patients over 0-10PIDs documents acceptable random variance distribution over time, absence of heteroskedasticity.

### Table S7

| GFAP bandset | Covariate | DF | DFDen | F Ratio | Prob > F | Effect ($\omega_p^2$) |
|---|---|---|---|---|---|---|
| Total GFAP log(ng/mL) | **Age** | 1 | 15.4 | 6.7 | 0.0200 | **0.25** |
| GFAP 50kDa log(ng/mL) | **Age** | 1 | 14.1 | 5.9 | 0.0290 | **0.23** |
| GFAP 45-49kDa log(ng/mL) | **Age** | 1 | 15.7 | 1.9 | 0.1900 | **0.05** |
| GFAP 37-39kDa log(ng/mL) | **Age** | 1 | 18.8 | 13.8 | 0.0015 | **0.38** |
| GFAP 20-26kDa log(ng/mL) | **Age** | 1 | 14.9 | 6.5 | 0.0227 | **0.24** |
| GFAP 15-19kDa log(ng/mL) | **Age** | 1 | 12.4 | 5.3 | 0.0401 | **0.23** |

**Table S7: Covariate effects of age and on GFAP CSF proteoform levels in TBI patients.** Listed are the results of linear mixed model covariate effect testing accounting for time elapsed after TBI and random donor attribute. Age effect on GFAP levels is computed using partial omega squared effect size ($\omega_p^2$), as it adjusts for variable sample sizes (see Methods). Age contributed significantly to uncleaved and total GFAP as well as fragment sets 15-39kD.

### Fig. S8

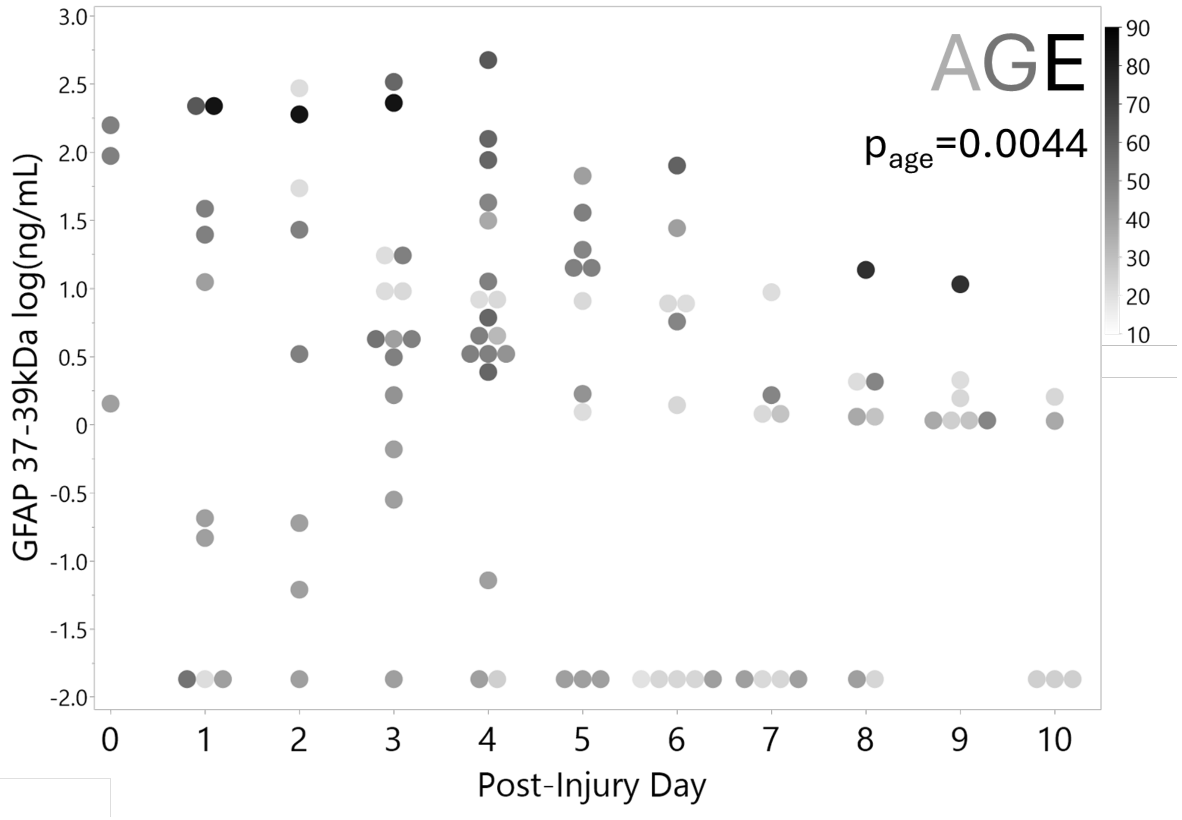


**Fig. S8: Covariate age effect on TBI CSF GFAP-BDPs:** Scatterplot shows 37-39kDa GFAP fragment levels over 10 PIDs with point grayscale for patient age. Older patients tended to have higher GFAP levels (darker points) than younger ones (lighter gray; p=0.004, n=102 observations).

### Table S8

#### GFAP bandset differences with alternate outcome partitioning

| Biomarker GFAP-BDP bandsets | Poor Outcome (GOSE 1-6) *18 patients, 81 CSF samples* | | Good Outcome (strict: GOSE 7+8) *3 patients, 18 samples* | | | *95% Confidence Interval* | | | Effect Size |
|---|---|---|---|---|---|---|---|---|---|
| | **Adjusted Mean (broad-Poor)** | *Std Error-Poor* | **Adjusted Mean (strict-Good)** | *Std Error-Good* | **Difference Poor-Good** | *Lower CI* | *Upper CI* | **p-Value** | **Cohen's d** |
| Total GFAP log(ng/mL) | **1.66** | *0.26* | **1.01** | *0.53* | **0.65** | *-0.53* | *1.83* | 0.260 | 0.92 |
| GFAP 50kDa log(ng/mL) | **0.79** | *0.23* | **0.68** | *0.48* | **0.11** | *-0.99* | *1.21* | 0.835 | 0.20 |
| GFAP 45-49kDa log(ng/mL) | **-0.08** | *0.29* | **-0.71** | *0.59* | **0.63** | *-0.69* | *1.96* | 0.325 | 0.65 |
| **GFAP 37-39kDa log(ng/mL)** | **0.88** | *0.26* | **-0.58** | *0.51* | **1.46** | *0.34* | *2.59* | **0.014** | 1.85 |
| GFAP 20-26kDa log(ng/mL) | **0.37** | *0.39* | **-0.56** | *0.85* | **0.93** | *-1.00* | *2.86* | 0.323 | 0.82 |
| GFAP 15-19kDa log(ng/mL) | **-0.64** | *0.35* | **-1.04** | *0.75* | **0.40** | *-1.32* | *2.12* | 0.623 | 0.37 |

**Table S8: Effect sizes of GFAP bandsets in CSF profiles of alternative dichotomized GOSE scores.**
Similar results with same significance occurrences are obtained when partitioning TBI patients with a stricter good recovery assigned only to those with GOSE 7+8 (3 patients, 18 samples). Poor outcome group included TBI patients with GOSE scores 1-6 (18 patients, 81 observations). Effect size Cohen's d was computed from the difference of the adjusted means derived from the linear mixed model repeated measures with age and time as fixed effects and patient ID as random effect. The difference was divided by the pooled SD, the squareroot of the residual variance component. The 95% confidence interval on the difference and is given in log(ng/ml).

Fig. S9

A

BNPS-skatole: cleavage proteins at tryptophans
(3-bromo-3-methyl-2-(2-nitrophenyl)thiol-3H-indole)

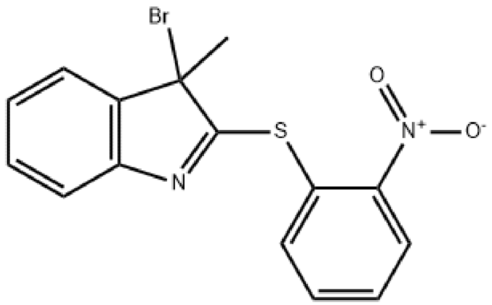


B C D

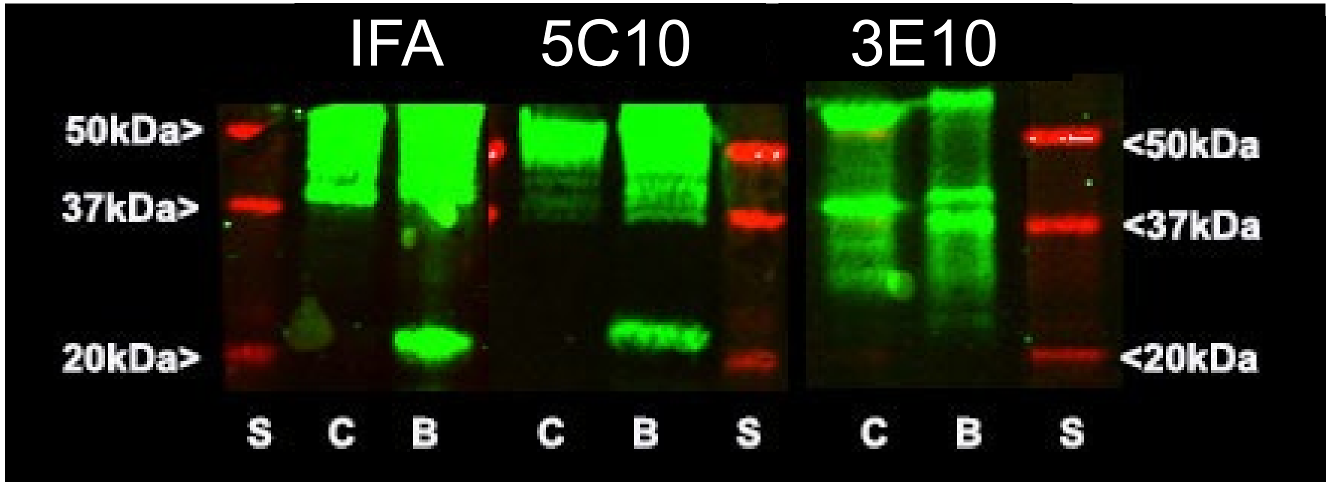


S= Molecular weight standard
C= Control ; B= BNPS-skatole treated

**Fig. S9: Partial chemical GFAP cleavage by BNPS-Skatole determined 5C10 epitope.**
This defined the novel TBI GFAP center cleavage in human traumatized astrocytes and TBI patients. **A)** Formula of BNPS Skatole, cleaving recombinant GFAP at sole tryptophan W256 that generated a N-terminal 34.6kDa fragment (harboring coil1) and a C-terminal 20.5kDa fragment (harboring coil2).
***B-D: Identification of two chemical GFAP fragments by two antibodies and determination of mab 5C10 binding site:*** Immunoblots of control ("C") and BNPS-Skatole ("B") treated recombinant GFAP along with molecular weight standard ("S", see Methods). All three clones detected larger GFAP fragments and full-length GFAP (top parts). **B)** Immunoblot probed by mab anti-pan-intermediate filament, IFA with known epitope in coil2B (aa 358-377, ALDIEIATYRKLLEGEENRI, see Table S3) confirmed that the C-terminal 20.5kDa band contains coil2. **C)** Blot probed using clone 5C10 bound to the IFA-recognized coil2, ~20kDa chemical GFAP cleavage product ("B"). Prior peptide binding studies determined 5C10 bound to a region within Peptide "AALKEIRTQYEAMASSNMHEAEEWYRSKF". The only part of this peptide that resides in the new BNPS-skatole cleaved fragment is YRSKF, thus defining the binding motif of 5C10. **D)** Mabs 3E10 and 2A5 confirmed the N-terminal ~35kDa chemical GFAP cleavage fragment harboring coil1 in "B" that was undetected by 5C10. Clone 5C10 did not detect trauma-generated small BDPs. This experiment determined 5C10 bound to motif 257 YRSK 260, allowing the conclusion that around this region lays the new TBI-induced GFAP center cleavage site (see text).

Fig. S10

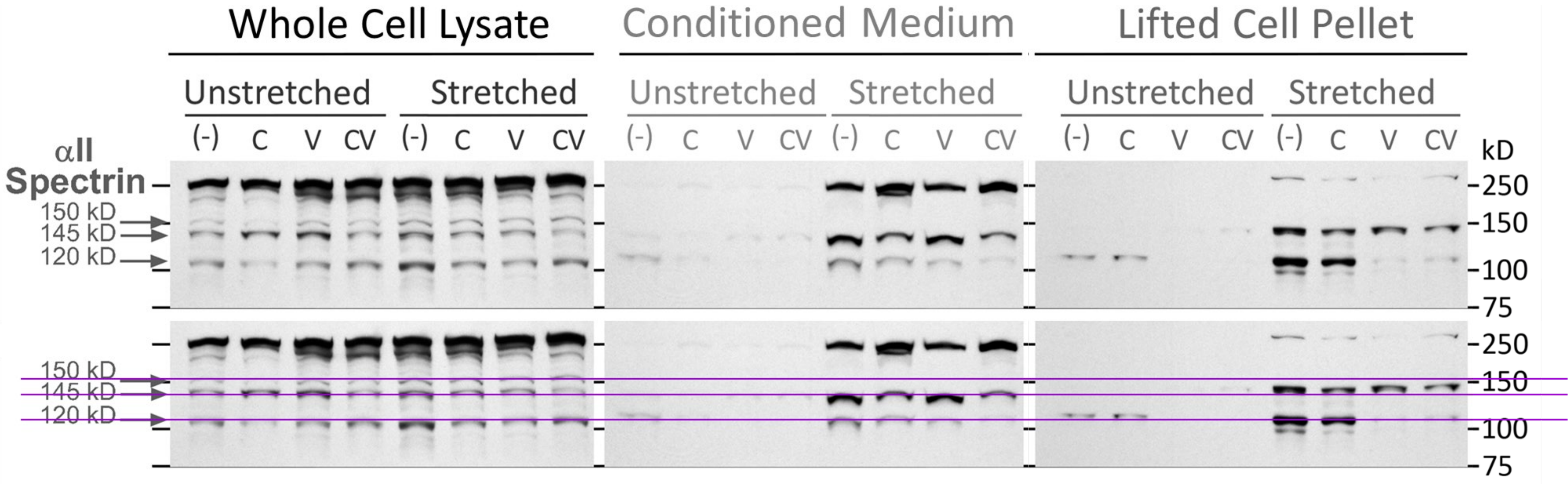


**Fig. S10: αII-spectrin degradation after stretch-injury confirms inhibitor specificity, calpain/caspase activity and shows distinct SBDP localization.**

Immunoblots show αII spectrin breakdown products (SBDPs, Enzo, 1:4,000), validated with protease inhibitors calpeptin (C, 13µM), and ZVAD-FMK (V, 11µM), both (CV) versus undrugged (-) control (Unstretched) and trauma (Stretched) samples in two-day post-trauma astrocyte whole cell lysate (WCL, 20µg/lane), concentrated conditional medium (CM, 6% of original volume/lane), and lifted cell pellet and debris (11µg/lane; detection ECL Pico, 5s exposures WCL, CM, 15s lifted cell pellet). Full-sized αII spectrin (250kDa) was abundant in WCL and CM and lacked in lifted degenerated material. Stretch-injury increased 145/150kD and 120kDa SBDPs. SBDPs migrated at 110 and 140kDa in fluid (see copy with purple lines for comparison). Lysed debris displayed 150 and 120/100kDa SBDPs. In WCL drugs C and V slightly reduced trauma enhanced SBDPs. In lifted/dead material V strongly inhibited the 120kDa and weakened 150kDa fragment, while C only slightly reduced 150kDa SBPD. In CM inhibitor C reduced trauma-released 145kDa SBDP and CV reduced trauma-released 120kDa and 145kDa SBPDs in fluids compared to trauma without inhibitors.

### Fig. S11

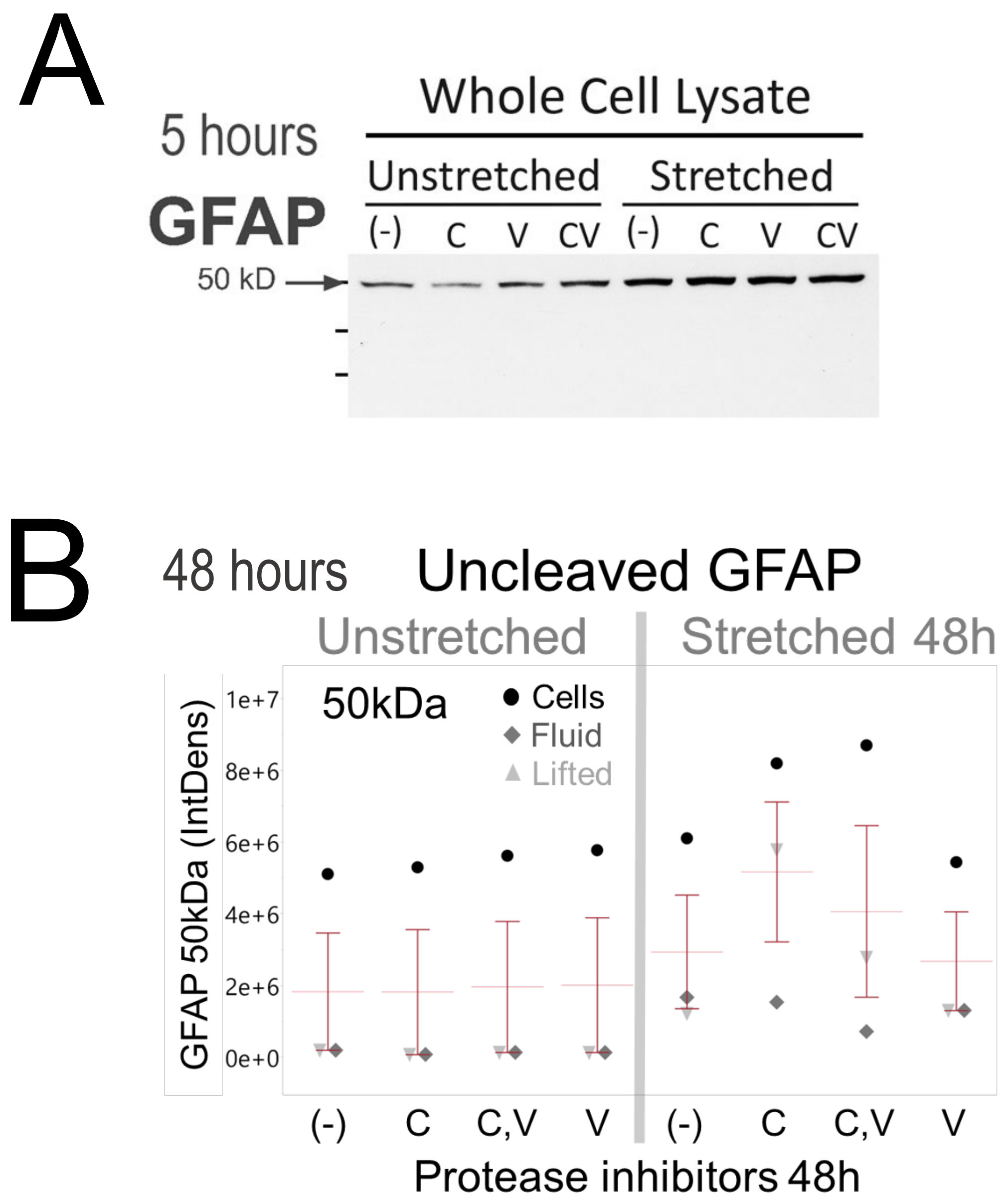


**Fig. S11**

**A) Acute increase in detergent-soluble uncleaved GFAP fraction suggest filament disassembly.** 2sec exposed immunoblot probed shows acute full-length GFAP increase within 5h after stretch injury in Cells, documents increased NP40-solubility of the GFAP cytoskeleton.

**B) Densitometry of** uncleaved GFAP signals at 48h postinjury increased in presence of calpeptin (C, C,V) across compartments, mainly due to levels in cells and lifted material. Scaled densitometry of subsaturated bands measured in a range of exposures (Table S5).

Fig. S12

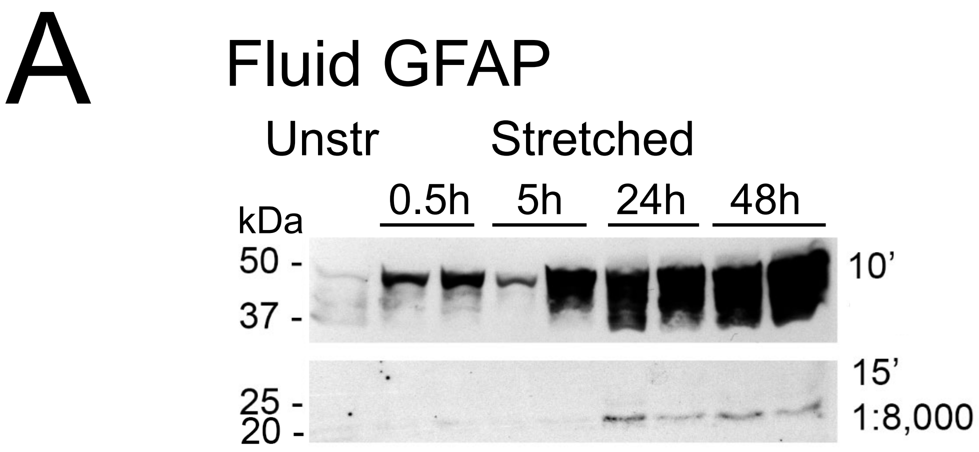


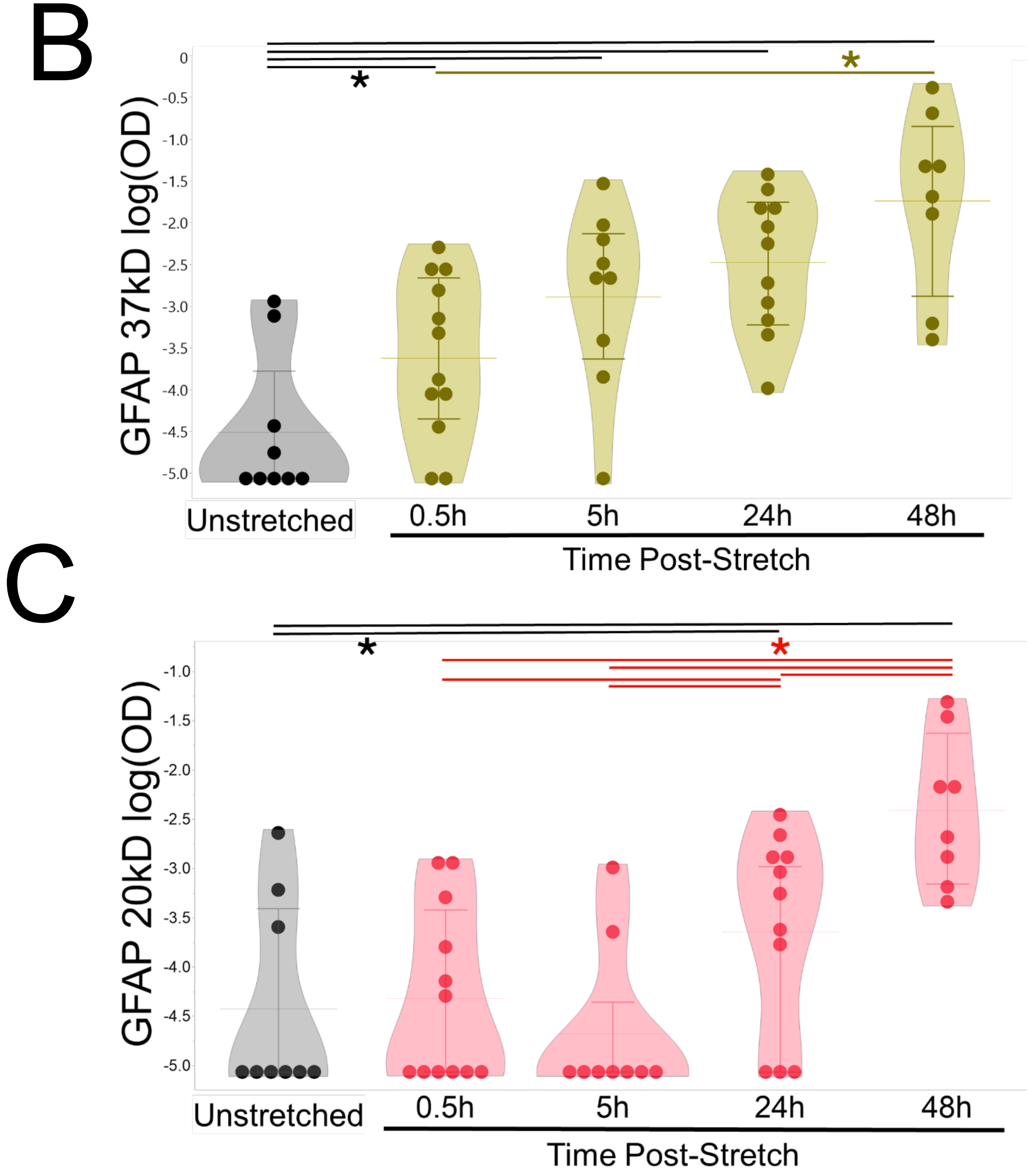


**Fig. S12: Timecourse of GFAP breakdown product release shows delayed small product elevation.**

**A)** GFAP immunoblot of concentrated conditioned medium (Fluid) at two exposures shows increasing GFAP signals between 0.5h and 48h after stretch injury (two samples per timepoint). The 37kDa band increased earlier than the 20kDa band. See Halford et al., 2017 for details. **B+C)** Violine-jitter plots show signal amounts (log optical densities, OD) for the 37kDa **(B)** and 20kDa **(C)** bands in control and stretch-injured cultures at 0.5, 5, 24, and 48 hours post-injury. Data are from 4-9 donors, with 7-12 samples per condition. Stretch severities ranged from 3-5psi. **B)** The 37kDa fragment was elevated versus control at all timepoints postinjury (0.5h: $p<0.05$ at 0.5h, Cohen’s d= 1.1SD units; 5-48h: $p<0.01$ d: 2-3.4). By 48h means were increased versus 0.5h by d=2.3 ($p<0.05$).
**C)** The 20kDa product increased at 24h ($p<0.05$, d=1.6) and more robustly at 48h ($p<0.001$, d=3) versus baseline. At 48h 20kDa levels differed from all earlier timepoints ($p<0.001$ versus 0.5 and 5h; $p<0.05$ versus 24h).
Error bars are means with interquartile range. Data were analyzed by linear mixed model repeated-measures with time as fixed effect and donor as random effect; effect sizes are differences standardized by residual variance reported as Cohen’s d.

### Fig. S13

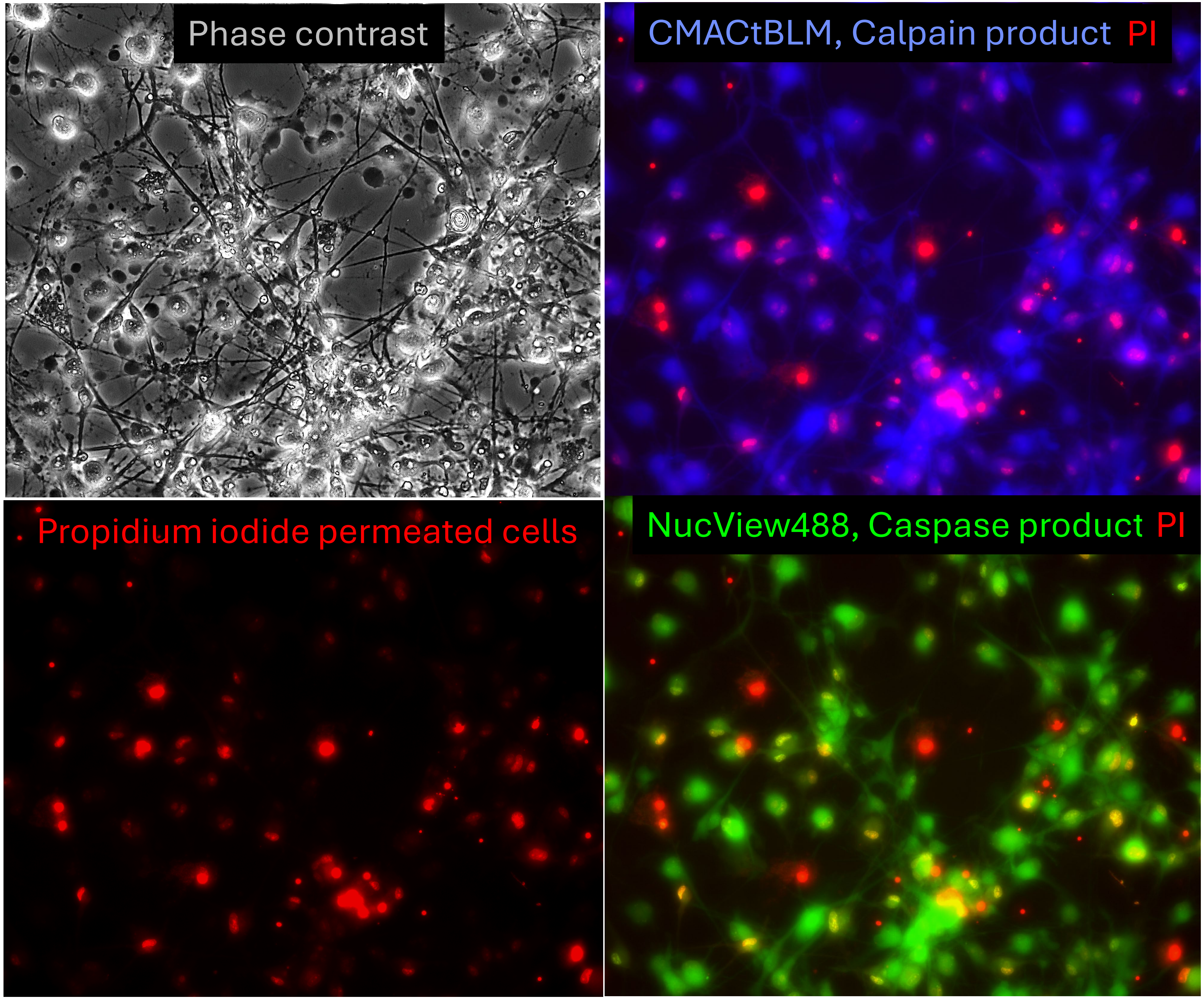


**Fig. S13: Separate channel documentation of Fig.7B for stretch-injured astrocytes.**

Live fluorescence imaging shows bushy (protoplasmic) astrocytes at 5h postinjury with concurrent calpain (blue) and caspase (green) activity in subpopulations of membrane-intact (PI-) or membrane-porated (PI+, red) astrocytes. Phase contrast shows membrane blebbing (phase-dark) without discernible protease activity. Integrity-compromised, sealed or re-sealed cells. Calpain reporter, CMACtBLM (blue) caspase reporter, NucView488 (green).

Fig. S14(A)

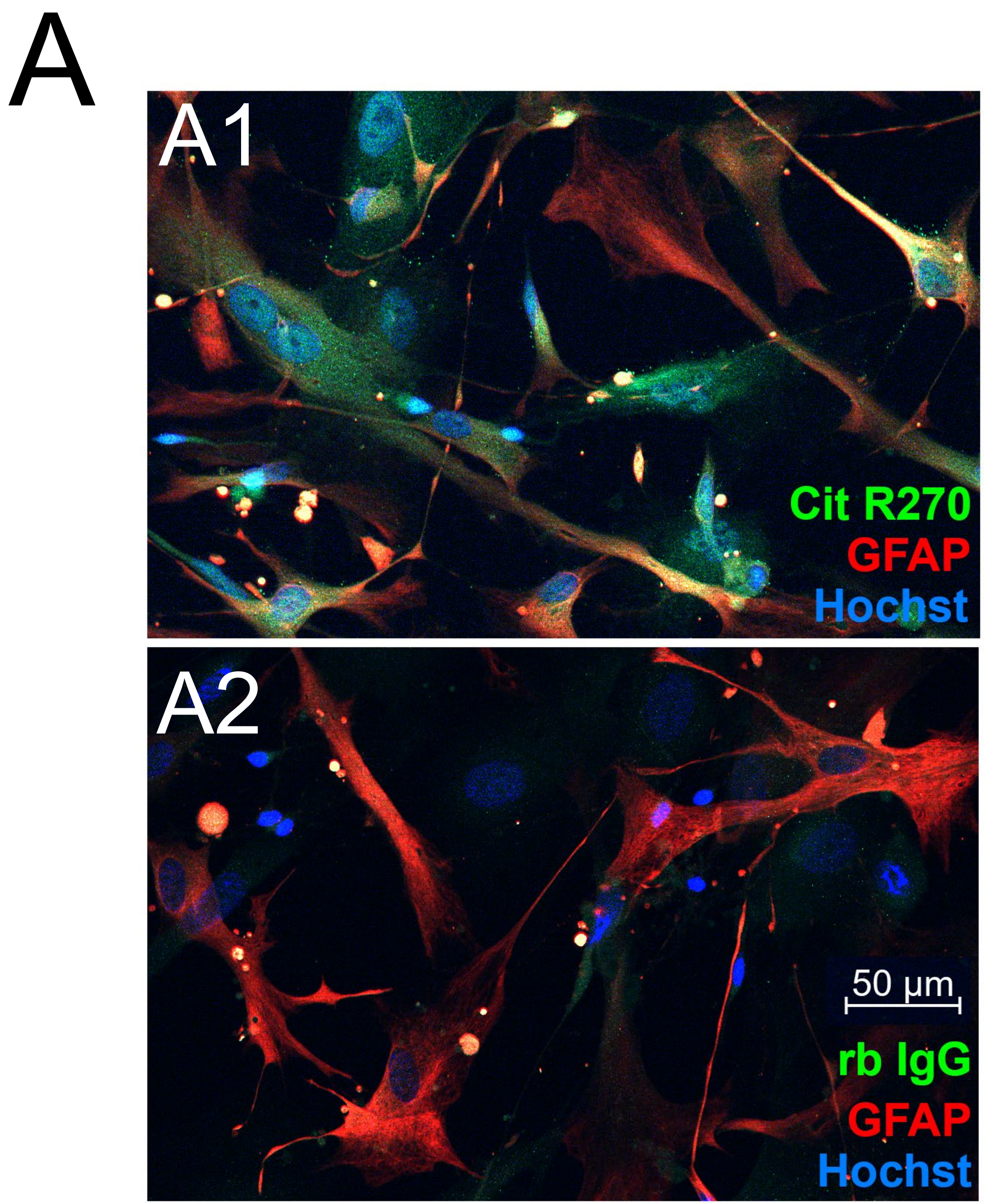


**Fig. S14: Citrullinated GFAP antibody control**

**A)** Rabbit antibody control for Cit270-GFAP counterstained for unmodified GFAP (polyclonal chicken anti-GFAP, red).

**A1:** Rabbit monoclonal antibody Cit-R270-GFAP (green) shows punctate non-filamentous signal in human astrocytes. **A2:** Rabbit immunoglobulins (rbIgG) incubated with same anti rabbit-Al488 secondary antibody (green) served as negative control.

Fig. S14 (BC)

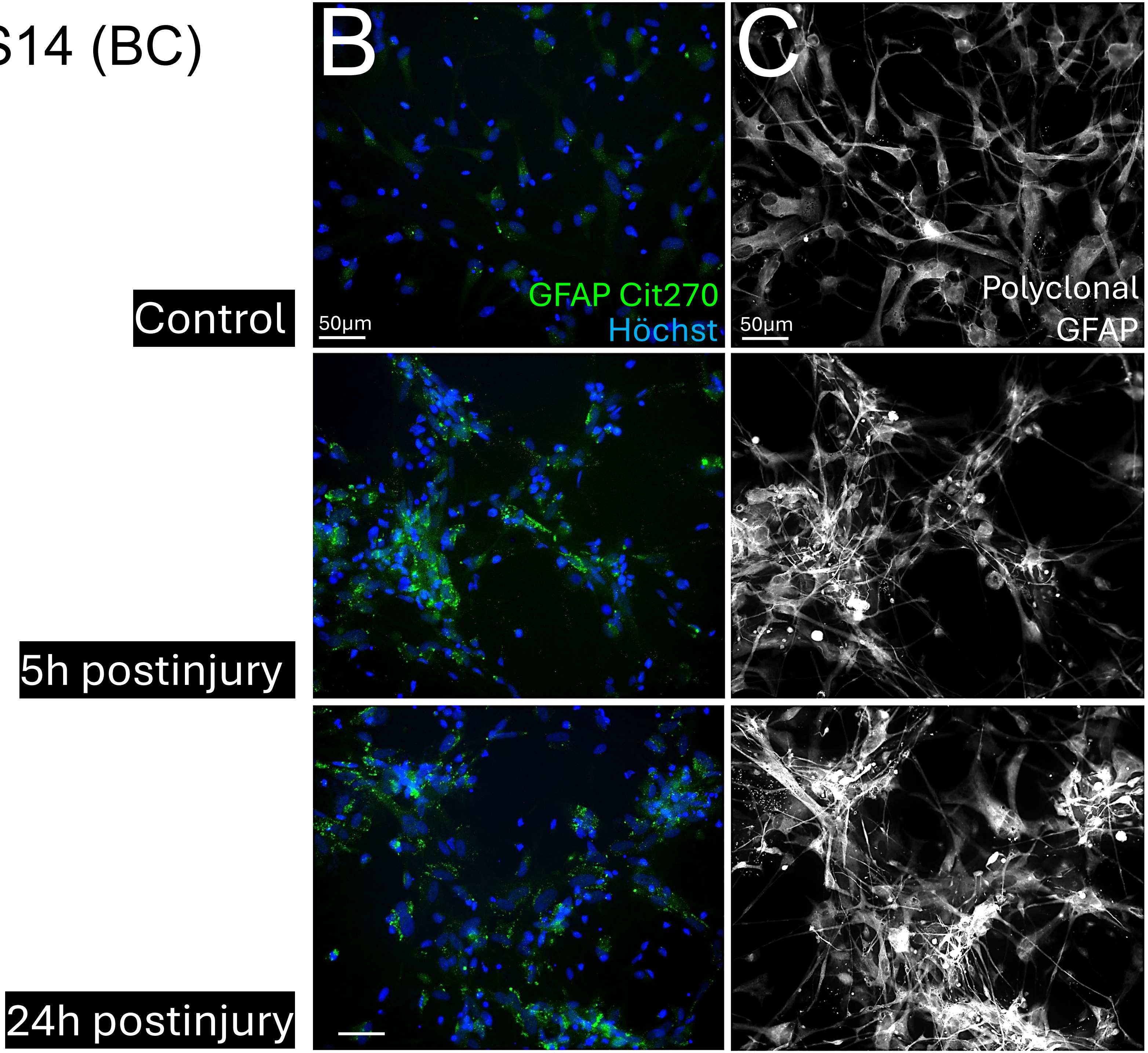


**Fig. S14 continued:**

*Citrullinated GFAP (Cit270), and unmodified GFAP (white) images without red channel shown in Fig.8.*

**B)** Low Cit-270-GFAP in unstretched human astrocytes (top, green; shown with nuclei, blue) compared with trauma-increase of Cit270 granules at 5h (middle) and 24h (bottom) postinjury.

**C)** Same images as (B) showing control astrocytes with faint cytoskeletal GFAP and increased GFAP at 5 and 24h postinjury using a chicken polyclonal anti-GFAP with anti-chicken Cy5 (white).

### Fig. S14 (DE)

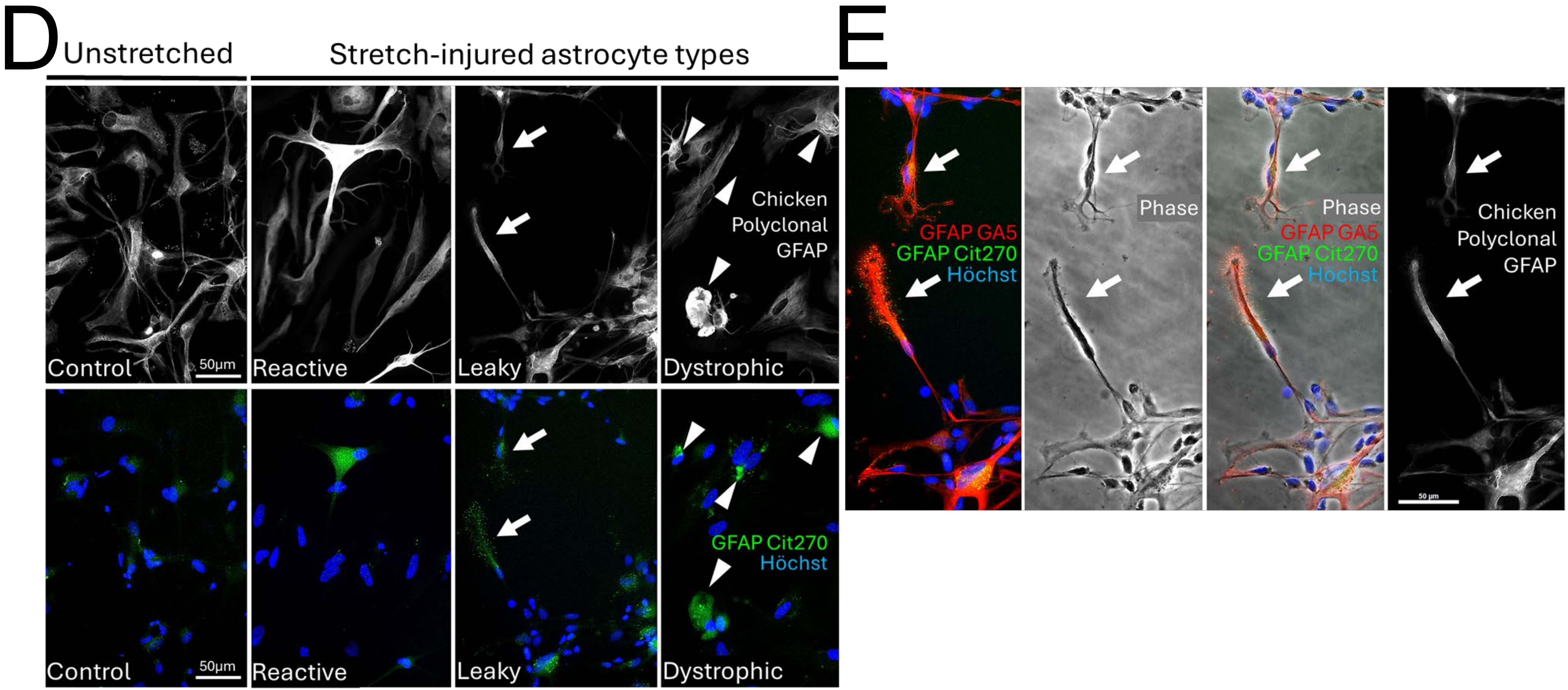


**Fig. S14** *continued: Cit270-GFAP and unmodified GFAP in astrocyte injury states.* **D)** Same as Fig.8B, here documenting unmodified GFAP (white, top) and Cit270-GFAP (green with nuclei, blue, bottom) signals for control, reactive, leaky and dystrophic astrocytes (from left to right) that document increase in unmodified and modified GFAP after stretch-injury. **E)** Same as Fig.8B, here showing GA5-stained astrocyte processes (red) surrounded by Cit270-GFAP particles (green) beyond lamellipodial process limits (phase) and unrelated to polyclonal unmodified GFAP structures (white).
